# *De novo* design of ligand binding proteins using large language models alone

**DOI:** 10.64898/2026.09.02.748987

**Authors:** Nam Hyeong Kim, A. Katherine Hatstat, Hyunil Jo, Yibing Wu, William F. DeGrado

## Abstract

Protein design has rapidly advanced with the advent of sequence- and structure-based machine learning models. However, reasoned design, which applies physicochemical principles and rules derived from sequence-structure-function relationships, has not seen the same benefits from generative machine learning models. Here, we test the ability of common large language models (LLMs; e.g. Claude, ChatGPT, and Gemini) to consider design principles to generate *de novo* proteins that bind metals and lipophilic small molecules without copying existing sequences. Common LLMs alone are able to 1) generate protein sequences to adopt a desired fold and bind the target ligand and 2) explain the principles that motivate the design choices. Following structure prediction and filtering, we selected a small set of designs (6 to 12 designs per query) for experimental validation, affording metal binders in one round of LLM-based design (25% hit rate) and perfluorooctanoic acid binders in two rounds (25% hit rate in the second round of design). Importantly, the LLMs produce detailed justification to accompany the *de novo* designed sequences, providing a conceptual framework on which designs can be evaluated. While the successful designs have some deviations from the prompted parameters and LLM-articulated design rationale, these campaigns provide a case study that highlights the utility of LLMs in making protein design more comprehensible and accessible to users without sophisticated design expertise.

## Introduction

Computational protein design has made brilliant progress in recent years,^1,2^ and scientists have gained the ability to design proteins that bind small molecules and protein targets. It has also been long possible to design proteins that bind metal ions as catalysts^3^ and dynamical switches to control protein dynamics^4–7^. There are several approaches to protein design in widespread use: sequence-based design, structure-based design, and rational design. In sequence design, one can use protein language models^8–11^ to draw from the wealth of sequence data to help design proteins intended to fold and function. These models learn statistical representations of sequences from evolutionary-scale protein sequence databases to generate new sequences independent of three-dimensional coordinates. Structure-based design focuses first on generating a three-dimensional structure using parametric equations^12,13^ or, more recently, hallucination, flow or diffusion^14,15^, which then serves as a template for the automated selection of a sequence^16,17^ that can fold into the given structure. The choice of the backbone can also be conditioned upon the structural requirements compatible with a given function such as metal-binding. In rational design, one applies physicochemical principles and rules derived from bioinformatic and empirical data to design proteins; for example, when the pattern of hydrophobic and polar residues along a protein sequence matches that of a secondary structure (e.g., 3 – 4 residues for the α-helix) the hydrophobic residues will line up along one face of the helix. The association of the apolar faces of the helices leads to a folded helical bundle. By varying the steric and physicochemical properties of the interacting sidechains, a variety of topologies of helical bundles, ranging from two-helix to eight-helix bundles have been accessed.^18–20^ Moreover, through reasoned design, humans can readily design sites capable of binding metal ions^7^ or organic small molecules^21–23^ based on simple rules augmented with detailed computational structural models.

Both sequence- and structure-based design have seen significant advances from generative models trained on the wealth of sequence and structural data now available, but rational design has not seen the same benefits. We postulate that large language model (LLM) inference provides an opportunity to formalize and apply established protein design principles without explicit structural generation or energy-based optimization.

To this end, we ask whether large language models (LLMs) alone are capable of reasoned design of proteins with two relatively simple functions: binding metal ions or forming a cavity to bind perfluorooctanoic acid (PFOA), a major member of the perfluorinated alkane substances class of pollutants. We ask whether one can simply query a common LLM such as ChatGPT,^24^ Gemini,^25,26^ or Claude^27^ to autonomously design proteins that bind these molecules without copying known sequences. Emerging reasoning-enhanced language models, which are designed for systematic problem solving, should, in principle be able to achieve this task by breaking it down into smaller, discrete tasks and applying encoded physicochemical priors to each subproblem. First, the models must synthesize information about apolar patterning to generate sequences that fold into helical bundles. Indeed, Ovchinnikov and co-workers show that reasoning models can generate amino acid sequences that fold into a helical bundle using a simple natural language prompt.^28^ To generate ligand-binding proteins, the models must capture various chemical principles: for example, Zn^2+^ is a d^10^ ion with a preference for tetrahedral geometry (or less favorably, octahedral or square pyramidal geometry) and carboxylate (Glu/Asp), imidazole (His) and thiolates (Cys) are good ligands. Motifs for binding metal ions including His-XXX-His/Glu figure largely in metal-binding sites. Similarly, perfluoro-alkanes prefer hydrophobic interactions, and the principles for designing helical bundles with an elongated channel that accommodate ligands like PFOA have been reported in the literature^22,23^. Such information is readily available and, in principle, might be assimilated by a language model.

To test this, we composed natural language prompts that detail our design goals and use Claude,^27^ Gemini,^25,26^ and ChatGPT^24^ to generate a limited number of sequences that satisfy the design criteria we outline. For metal binding proteins, the models articulate reasonable physicochemical principles for the designated metal binding task and generate sequences that fold into helical bundles with moderate confidence. We then experimentally characterized LLM-generated designs and found that several candidates express, have the desired secondary structure, and show evidence of metal binding. Initial

PFOA binders (specifically, five-helix bundles with hydrophobic central cavities) showed no appreciable PFOA binding. Thus, we returned to the LLM with a hypothesis about an alternate structural topology informed by work from the Woolfson laboratory.^22^ With this single iteration on the prompt based on relevant literature, we obtained improved PFOA binders with an experimental hit rate of 25%. While we obtain hits in both design campaigns, the LLM-generated designs do not always satisfy the structural features delineated in the prompt, highlighting limitations in the structural reasoning of the LLMs. Further, redesign of LLM-generated proteins with state-of-the-art design tools affords different sequences with improved structure prediction confidences, indicating that the LLMs can generate feasible but suboptimal sequences. These relatively modest accomplishments demonstrate both the strengths and limitations of LLMs to apply computer reasoning to the construction of functional proteins from first principles, and they bring valuable information that might be advantageously combined with structure-based approaches in a modern agentic workflow.

## Results

### LLMs generate metal binding, helix-turn-helix homodimers in a single round of design

In this work, we first gave several LLMs the task of designing *de novo* proteins with Cu^2+^- and Zn^2+^-binding sites. Because previous studies have designed proteins to bind these ions,^3,7,29–36^ we prompted for the LLM to not copy previous work or use motifs that had been used previously for constructing metal-binding proteins. We were interested in LLM-guided design independent of structural input, so we articulated a prompt and explored the sequences designed by the following LLMs: ChatGPT (5 and 5 advanced “thinking” model)^24^, Claude (Sonnet 4.5)^27^, and Gemini (model 2.5).^25,26^ The entire prompt was ~800 words (**Supplemental Note 1**) and was written in about 10 minutes, and the same prompt was presented to each LLM. The prompt architecture contains four key components, summarized below (**Fig. 1B**):

- **Motivation and context:** how proteins bind specific metals and how metal binding information can be transmitted to other domains in multidomain proteins
- **Structural features**: homodimeric four helix bundle comprised of two helix-turn-helix motifs of approximately 60 residues/monomer, with N- and C-termini within 15Å
- **Chemical logic**: bind two metal ions, with binding sites comprised of His residues instead of Asp/Glu/Cys in the first shell
- **Deliverables**: generate 10 sequences following the design parameters described and explain the rationale for the design choices made

**Figure 1.**
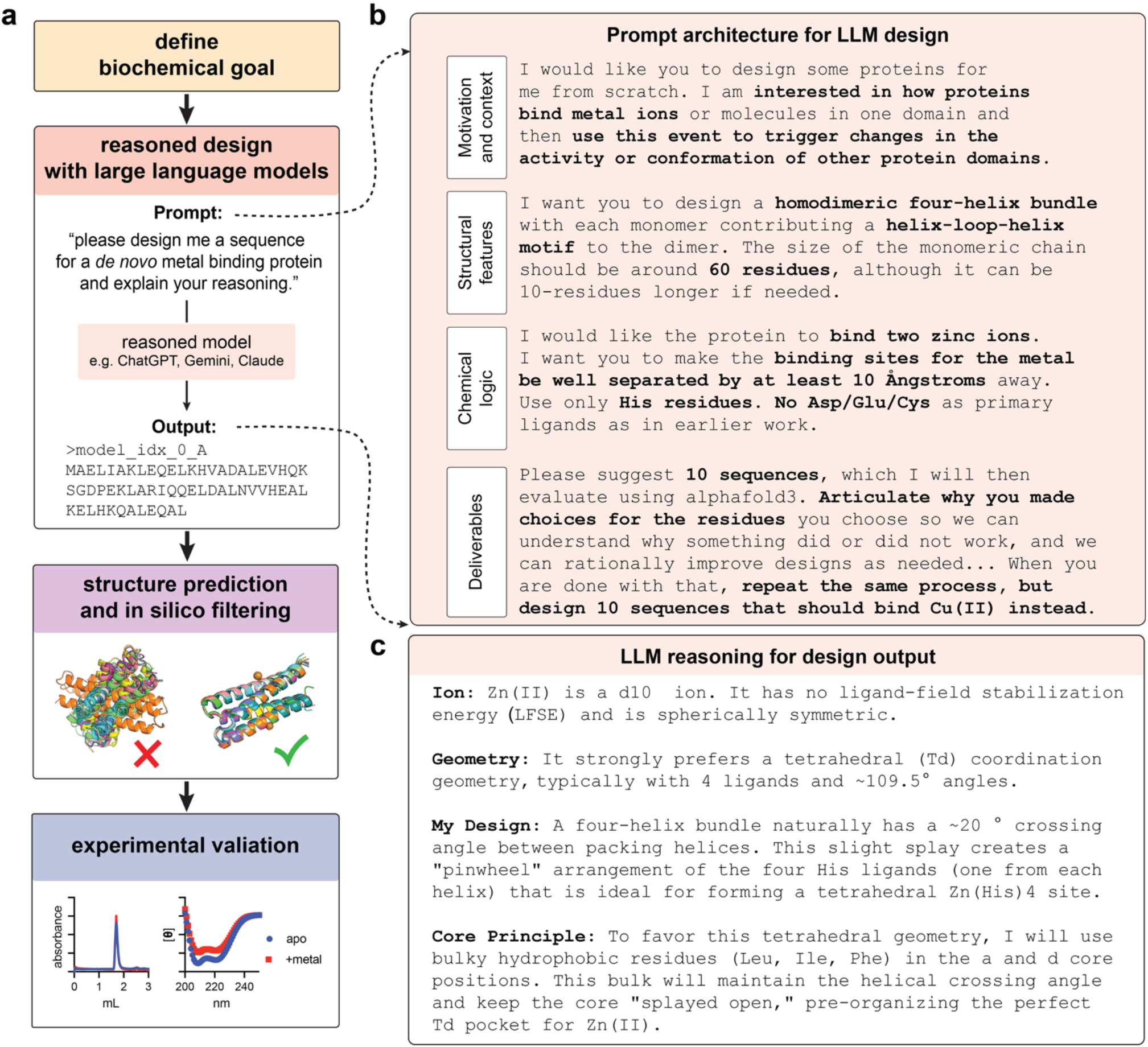
Natural-language models enable design of *de novo* proteins from first principles. (**A**) A design campaign starts with defining a biochemical goal, followed by computational design, candidate selection, and experimental validation. In this work, we complete all computational design (culminating in *de novo* sequence candidates) using off-the-shelf natural language models alone instead of the traditional structure-based design campaign. (**B**) Here, we construct a prompt to articulate the biological motivation and context of the design task, define general structural features and chemical principles desired in the designed proteins, and delineate the deliverables we expect from the NLM, namely a specific number of *de novo* designed sequences and articulated rationale for the design choices made. (**C**) An excerpt from NLM outputs that detail the rationale for the design of metal binding bundles. See Supplemental Notes 1–4 for full LLM prompt and responses for metal binding protein design.

In all cases, the LLMs provided designed sequences and rationale in only a few minutes (**Fig. 1C;** full LLM responses provided in **Supplementary Notes 2–4**). While the format of the responses varied, the models uniformly captured physicochemical concepts important for both *de novo* helical bundles (e.g. heptad repeat, helix crossing angle, loop architecture) and metal binding. The LLMs were able to reflect on metal electronic configuration and ligation preferences, as well as the design of scaffolds that would support residues for the desired binding function. For example, Gemini rationalized that Zn^2+^ prefers a tetrahedral coordination geometry, prioritized a 4His binding site, and sought to preferentially place bulky hydrophobic residues at positions *a* and *d* of the helical heptad repeat to maintain core packing and a helical crossing angle favorable for forming the desired Zn(His)_4_ binding site (see **Supplementary Note 3**). The models were also able to rationalize features that would enforce selectivity of designs for Zn^2+^ vs Cu^2+^: To drive the binding site toward Cu^2+^ selectivity over Zn^2+^, His pairs should be positioned to favor square planar geometry with second-shell residues that penalize tetrahedral geometry (**Fig. 1C; Supplemental Notes 2**,**3**). In total, we generated 10 designs per metal (Zn^2+^ and Cu^2+^) with four LLM models (Gemini, Claude, ChatGPT base, ChatGPT advanced “thinking” model), affording 10 sequences*2 metals*4 LLMs = 80 total sequences.

We sought to evaluate the sequences on several metrics: 1) sequence novelty, 2) designability, 3) propensity for homodimer formation, and 4) metal binding capacity. To assess sequence novelty, the designed sequences were first queried via both protein BLAST (BLASTp)^37,38^ and sequence similarity searches in the Protein Data Bank (PDB) to ensure that the models were not simply copying natural or designed sequences that have been previously reported. For all sequences generated across all models, neither BLASTp nor PDB sequence queries returned any similar sequences. While all models articulated similar design principles and chemical logic, each sampled different sequence spaces in the designs (**Fig. 2A; Tables S1,2**). For the Zn^2+^ binding designs, Gemini preferred highly charged and repetitive sequences with repetitive Lys, Glu, Leu motifs, while Claude generated the most diverse sequences with varied lengths and with asymmetry in the helices of the helix-turn-helix motifs. Interestingly, ChatGPT (free base model) generated sequences with high Asn, Thr, and Ser content while the more advanced thinking model generated sequences that converged on a well-defined sequence pattern with limited variation between designs (**Fig. S1**).

**Figure 2.**
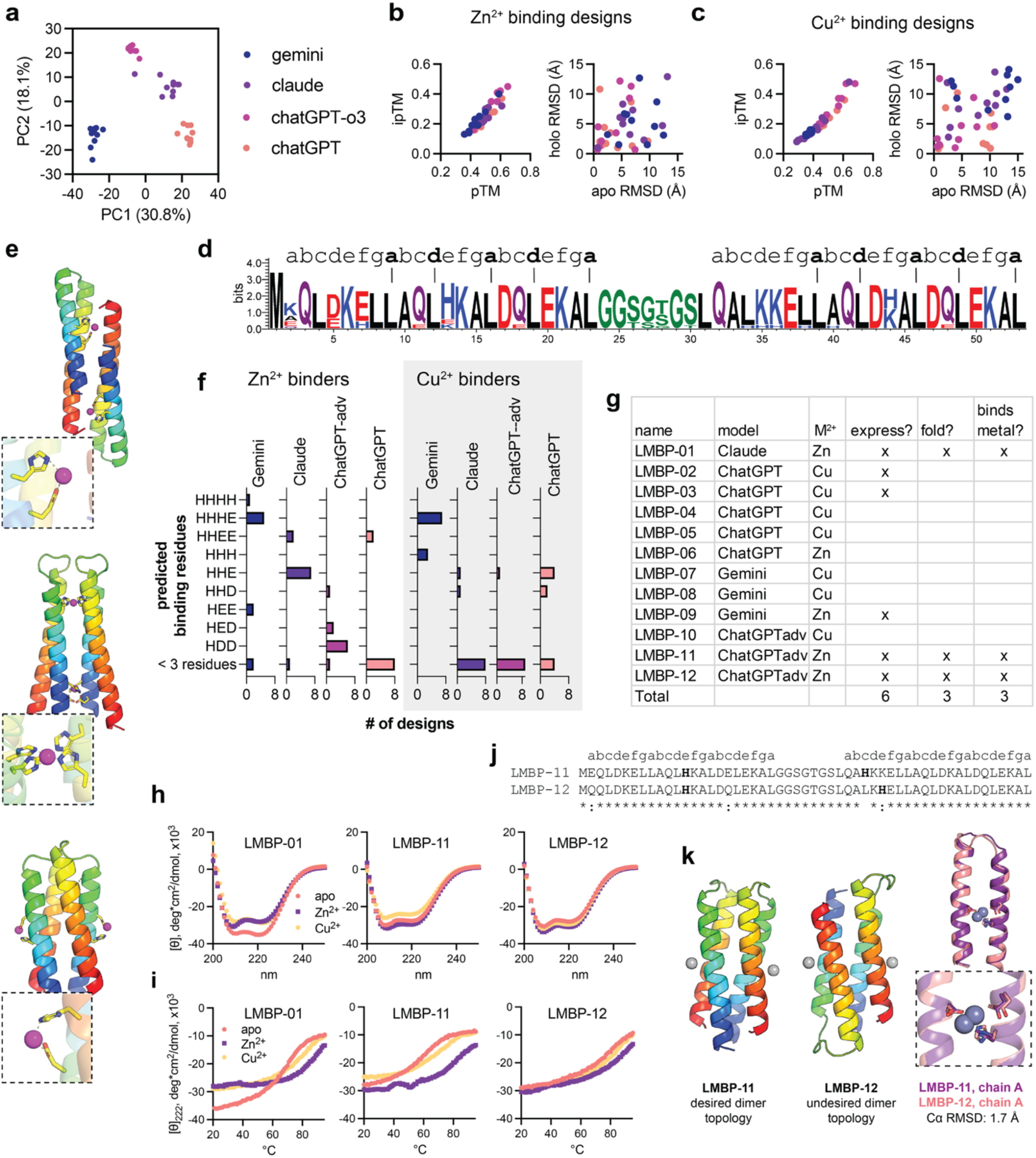
LLMs generate *de novo* metal binding proteins. (**A**) Principal component analysis of ESM2 embeddings show that different LLMs sample different sequence space. (**B**,**C**) Structure prediction confidence scores and inter-model RMSD for Chai-1 models of LLM-generated metal binding proteins (LMBPs). (**D**) Sequence logo of zinc binding designs generated by ChatGPT-advanced show that designs follow the principles established for design of four helix bundles. Heptad repeat (*abcdefg)* pattern is annotated above the sequence logo, with bold/dashes indicating key hydrophobic residues at *a* and *d* positions. (**E**) Structure prediction models of representative Cu^2+^ binding designs, highlighting the diversity of helical bundle topology and (inset) metal binding sites generated. (**F**) Distribution of predicted metal binding motifs by model and target divalent metal. Predicted binding motifs determined by visual inspection of Chai-1 models for each design. (**G**) Summary of experimental characterization effort for the 12 selected design candidates, termed LMBPs. (**H**) Far-UV circular dichroism spectra of apo (10µM LMBP) and holo (10µM LMBP with 40µM Zn^2+^) samples of LMBP-01, LMBP-11, and LMBP-12 show all three have helical character. (**I**) Thermal denaturation circular dichroism experiments show that the LMBP-01, LMBP-11, and LMBP-12 are selectively stabilized by cognate ion Zn^2+^ relative to non-cognate ion Cu^2+^. (**J**) Sequence alignment of LMBP-11 and LMBP-12, which vary by 4 residues. Despite high sequence similarity, the bundles have (**K**) highly similar structure in the monomeric helix-turn-helix motif (right) but different predicted dimer topologies (left).

Following sequence generation, designs were refolded with AlphaFold3 (AF3)^39^ and Chai-1^40^ to assess predicted fold and metal coordination (**Fig. 2B, Fig. S2**). We assess designs by model confidence and self-consistency (measured by inter-prediction root mean square deviation (RMSD)) where higher confidence scores and low inter-prediction RMSD indicate that a design is more likely to adopt a given structure. Across all designs, the confidence scores were low to moderate (e.g. average Chai-1 pTM = 0.49, ipTM = 0.25 across Zn^2+^ binders; average Chai-1 pTM = 0.45, ipTM = 0.21 across all Cu^2+^ binders) (**Fig. 2B**,**C; Fig. S2**). Confidence scores for design predictions differed between AF3 and Chai-1 (**Fig. S2C**). Zn^2+^ binders had higher confidence in Chai-1 predictions than in AF3 predictions, with three designs with Chai-1 pTM scores > 0.6 (sequences from ChatGPT, ChatGPT-advanced, and Claude). Cu^2+^ binders showed the opposite trend, with higher confidence AF3 models than Chai-1 models, with five designs with pTM scores between 0.6–0.8 (sequences from ChatGPT, ChatGPT-advanced, or Gemini). Notably, the confidence scores are well below the thresholds we typically consider reasonable in structural biology or when using confidence metrics for *in silico* selection of design candidates, with confidence cutoffs of >0.9 (pLDDT or pTM) in typical design campaigns.

Across the design candidates, the sequences are predicted to adopt the desired helix-turn-helix topology, favoring helix-breaking glycine, serine, and proline residues in the interhelical loop. The LLMs capture design principles that should drive homodimer formation by preferentially placing hydrophobic residues at the *a* and *d* position of the heptad repeat to produce a hydrophobic core in the assembled four-helix bundle (**Fig. 2D**). However, not all designs are predicted to have the desired topology (parallel assembly of helix-turn-helix hairpins with all N and C termini within 15 Å of each other). The structural models also display a diversity of predicted metal binding motifs, despite specifically prompting the LLMs to design 4His binding sites (**Fig. 2E**). Across the designs, there are both internal and surface binding motifs (**Fig. 2E**), and sequences from different LLMs have different distributions of predicted binding site residues (**Fig. 2F**). Only Gemini produced sequences that are predicted to have requested 4 residue binding motifs, and only one design is predicted to contain the desired 4His binding site. Between Zn^2+^ and Cu^2+^ binder designs, the Zn^2+^ binders have more reasonable metal binding motifs, while a majority of the Cu^2+^ binder models have predicted metal binding sites containing fewer than 3 amino acids (**Fig. 2F**). Thus, while the LLMs can rationalize design principles (**Fig. 1C; Supplemental Notes 2**,**3**), there are discrepancies between user-described design parameters, the rationale articulated by the LLMs, and the LLM-generated sequences.

After filtering for self-consistency (low RMSD between multiple structure prediction samples) and manual inspection, we selected twelve design candidates for experimental validation, herein termed LLM-designed Metal Binding Proteins (LMBPs) (**Fig. 2G**). The design candidates selected consist of both Zn^2+^ and Cu^2+^ binders across multiple LLMs with various predicted topologies and metal binding motifs. Of the twelve designs, six expressed in *Escherichia coli* and were successfully purified. Three of the six purified designs displayed the desired helical secondary structure by circular dichroism (CD) (**Fig. 2H**) and formed monodisperse species in analytical size exclusion chromatography (SEC), with an elution volume consistent with that expected for a dimeric complex (**Fig. S3**). Metal binding was first assessed by measuring the thermal stability of each design in the presence and absence of the cognate metal ion (Zn^2+^ for LMBP-1, LMBP-11, and LMBP-12). In all cases, addition of the cognate divalent cation, Zn^2+^, stabilizes the designs to a greater degree than addition of non-cognate cation (Cu^2+^) (**Fig. 2H**,**I**), indicating that the designs preferentially bind the desired divalent cation. Zn^2+^ binding to LMBP-11 was further validated by isothermal titration calorimetry, affording an apparent dissociation constant (K_D_) in the double-digit µM range (K_D_ = 30.1 µM ± 7.67 µM; **Fig. S4**). This is consistent with the expected affinity given that LMBP-11 is predicted to coordinate Zn^2+^ through a surface three-residue site (His/Asp/Asp).^29,34,36,41–47^

Two of the successful designs, LMBP-11 and LMBP-12 were both generated with ChatGPT-advanced and vary by only four amino acids (**Fig. 2J**). Despite good agreement between the predicted structure of the helix-turn-helix monomer in LMBP-11 and LMBP-12 (1.7Å Cα RMSD; **Fig. 2K**, right), the designs show different predicted homodimer topologies, with LMBP-11 predicted to assemble with the desired parallel dimer topology and LMBP-12 with the undesired antiparallel dimer topology (**Fig. 2K**, left). In the predicted structures, the topology of LMBP-12 results in an intra-chain salt bridge between D5 and K35’ (both *d* positions of the heptad repeat, facing the dimer interface) that is not present in LMBP-11 (**Fig. S5**). This additional salt bridge should serve to stabilize the dimer.^48,49^ Indeed, LMBP-12 exhibits higher thermal stability than LMBP-11 in thermal denaturation experiments (**Fig. 2H**). In both LMBP-11 and LMBP-12, the predicted metal binding site is comprised of one His and two Asp residues. This binding motif deviates from the prompt in which we specified that the model use only His residues and not Asp/Glu/Cys residues. Indeed, the residues that the LLM indicated as key Zn^2+^ binding residues are in positions too far apart in 3D space to accommodate metal binding (**Fig. S6**). In both cases, the deviations in topology and metal binding motif may be more carefully constrained in structure-based design pipelines.

As a final test of backbone designability and sequence novelty, we selected the highest confidence design from each LLM and subjected each backbone to redesign with ProteinMPNN^16^ and refolding with AlphaFold2.^50^ AF2 models of MPNN-designed sequences have higher confidence than the input model for all LLMs except ChatGPT (**Fig. S7A**). Further, ProteinMPNN-designed sequences cluster away from the LLM-generated designs in a principal component analysis of ESM2^8^ embeddings, indicating that ProteinMPNN samples different sequence space for the top backbone than that sampled in LLM-based design (**Fig. S7B**). Thus, while LLMs generate sequences that fold into the desired topology, the models are not sampling the most designable sequence space for the given backbone.

### LLM generated helical bundles bind perfluorooctanoic acid

As metal binding protein design is well-established in the literature, we next sought to test the performance of LLMs on a more challenging design task. Ligand binding design is still an outstanding challenge in the field, with successful design campaigns relying on structural informatics, robust scaffold and binding site motif selection, and often, large-scale screening campaigns to identify a few lead designs.^17,51–55^ Here, we tested the ability of LLMs to generate ligand binding proteins without structural informatics or high-level user input in the design process.

For this task, we sought to design binders for perfluorooctanoic acid (PFOA), a persistent “forever chemical”^56^ for which a protein binder would be useful for adsorption and bioremediation strategies. Here we proposed a simple binding hypothesis: the binder should contain a hydrophobic cavity to accommodate the perfluoroalkyl tail and charged residues to stabilize the charged carboxylate head group of PFOA. We reasoned that a helical bundle with a hydrophobic central cavity and charged residues in interhelical loops near the openings of the cavity would satisfy these requirements. We queried LLMs with a brief prompt (<200 words) that included the SMILES string for PFOA, a short description of the desired structural features (e.g. five helix bundle, apolar groups in the protein core, charged residues in the loops, binding two molecules of PFOA) and the desired output (10 design sequences) (**Fig. 3A; Supplemental Note 4**). Again, the LLMs were able to rationalize design choices consistent with established principles for design (**Supplemental Notes 5**,**6**). For example, Gemini reasoned that the designs should include features such as:

- `Helices use a coiled-coil-like heptad pattern with L/I/V at putative core positions and mostly Glu/Lys surface`.
- `Loops are short, enriched in Lys/Arg/His for the PFA carboxylate, and include Gly/Pro to favor hairpins`.
- `Each design has at least one Trp and one Tyr (most have more) for concentration determination and potential fluorous/π interactions near the ends`.

**Figure 3.**
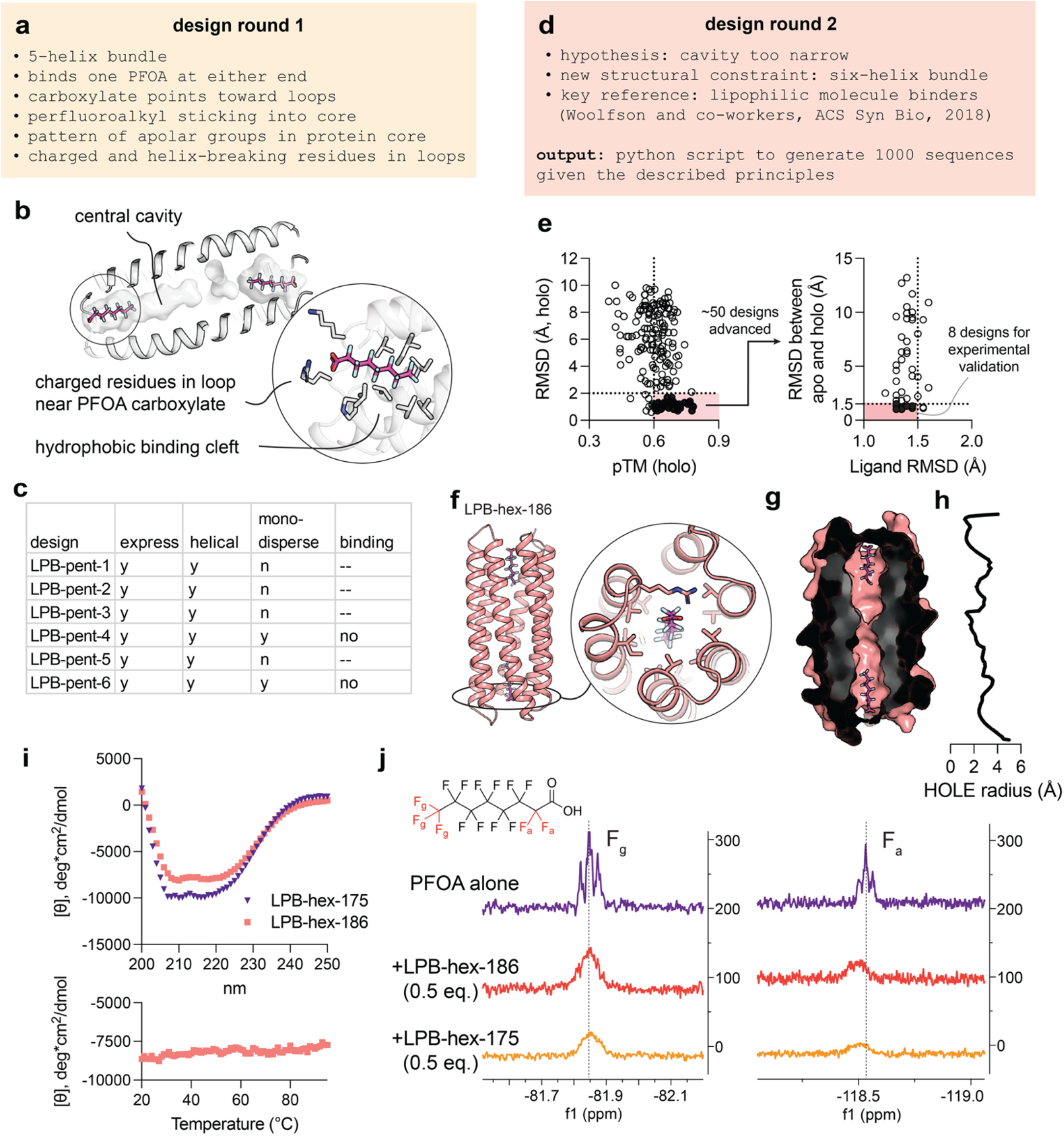
LLM designed perfluorooctanoic acid binding proteins. (**A**) Design round 1 involved prompting for a five-helix bundle with a central cavity that binds PFOA at either end, with the perfluoroalkyl tail in a hydrophobic central cavity and the carboxylate group stabilized by charged residues in interhelical loops. (**B**) Structure prediction reveals designs with the desired features. (**C**) Summary of experimentally characterized candidates from the first round of design. (**D**) Based on established principles from Woolfson and co-workers,^22^ we revised our design prompt and generated a series of six-helix bundles with the same biochemical features. (**E**) design candidates were filtered in two steps based on confidence score, RMSD of holo models, RMSD between apo and holo models, and ligand RMSD across models. (**F**,**G**) Top design candidates are predicted to have the desired structural features: six helix bundle topology with hydrophobic central cavity and charged residues on interhelical loops to stabilize the PFOA carboxylate. (**H**) HOLE analysis of the pore radius of LBP-hex-175. (**I**) Top candidates LBP-hex-175 and LBP-hex-186 have helical character and high thermostability. (**J**) 19F NMR of PFOA +/− sub-stoichiometric concentrations of LBP-hex-175 and LBP-hex-186, where addition of protein induces broadening of resonances corresponding to the terminal trifluoromethyl group (left) and alpha fluorine atoms adjacent to the carboxylate head group (right).^57^

We then asked the model to generate a python script that would produce 1,000 sequences that fit the established design rationale (**Supplemental Note 6**). We generated sequences and predicted structures with Chai-1 of 100 of the 1,000 designs without and with ligands (two equivalents of PFOA as per the design prompt). While the models had moderate confidence for the fold accuracy (average pTM = 0.49), they had low confidence for the complex (average ipTM = 0.2) (**Fig. S8**). Nonetheless, we filtered the designs for self-consistency in both apo and holo structure prediction models (<5 Å backbone RMSD), affording 6 candidates for experimental validation (termed LPB-pent for LLM-generated PFOA binding pentamer). The final designs showed the desired design features (**Fig. 3B**) but sample different sequence space (30% and 62% average pairwise sequence identity and similarity, respectively, in all vs all comparison). All six designs expressed and displayed helical secondary structure, and two of the six (LPB-pent-4 and LPB-pent-6) showed a major, monodisperse peak at the expected elution volume by analytical size exclusion chromatography (**Fig. S9**). We then screened LPB-pent-4 and LPB-pent-6 for PFOA binding via 19F NMR, where we expect resonances corresponding to the perfluoroalkyl chain of PFOA to shift or broaden upon PFOA:protein binding.^57^ We observed minimal broadening of the resonances corresponding to the terminal trifluoromethyl (~80 ppm) and alpha-fluorine atoms adjacent to the carboxylate head group (~117 ppm) of PFOA upon titration of the designed binders at sub-stoichiometric ratios (**Fig. 3C, Fig. S10**).

Returning to our design rationale, we hypothesized that the cavity of the designed five-helix bundles may be too constrained to allow PFOA binding (**Fig. 3D**). Indeed, previous efforts by Woolfson and co-workers^22^ toward design of helical bundles showed that hexameric bundles with open central cavities bound small hydrophobic dyes and lipophilic molecules with higher affinity than pentameric bundles with narrower cavities. To build on these principles, we provided the reference and prompted:

~~~
I would like you to read the paper I attached and apply the principles
of α-helical barrels with a central, accessible channel to the Python
code for sequence generation. The authors expanded the traditional
heptad repeat.
~~~

Using the generated python script (**Supplemental Note 6**), 1000 sequences were produced and 200 were selected for modeling with Chai-1 in apo and holo states (2 equiv. PFOA). The six-helix bundle designs had higher confidence overall relative to the five helix bundles (average pTM for holo models = 0.62 across 200 designs, versus average pTM = 0.49 for five helix bundles). Designs were filtered in two steps: first, candidates were selected by holo model confidence (pTM > 0.6) and self-consistency (Cα RMSD < 2.0Å). Then, this subset was further filtered for candidates with high ligand placement consistency (ligand RMSD <1.5Å) and low RMSD between the apo and holo structure (Cα RMSD < 1.5Å) to select designs in which the binding cavity would be pre-organized, and binding would incur a minimal entropic cost. After filtering, eight candidates were advanced to experimental validation (termed LPB-hex for LLM-generated PFOA binding hexamer) (**Fig. 3E**). Inspection of models of the final candidates showed that the designs adhere to the desired principles: the bundles are predicted to accommodate a PFOA ligand at either end of the central cavity, with charged residues on loops near the carboxylate head groups and patterned apolar groups along the pore lining (**Fig. 3F**), and the central cavity size is comparable to that of previously reported lipophilic molecule-binding bundles^22^ (LPB-hex-186 average pore diameter = 5.1 Å; **Fig. 3G,H**).

Two designs (LPB-hex-175 and LPB-hex-186) induced broadening of NMR resonances corresponding to the terminal trifluoromethyl group (~82 ppm) and alpha fluorine atoms (~118 ppm) of PFOA^57^ upon addition of 0.5 equivalents of the protein (**Fig. 3J**). LPB-hex-175 and LPB-hex-186 reduced the peak area of the trifluromethyl resonance by 90.8% and 87.3%, respectively. These top candidates have the same overall structural topology (Cα RMSD = 1.8Å; **Fig. S11**) but low pairwise sequence identity. While both candidates have helical character (**Fig. 3I**), neither was monodisperse by SEC (**Fig. S12A**). In fact, LPB-hex-175 formed a species with a molecular weight consistent with that of a trimer which persisted even after boiling and SDS-poly acrylamide gel electrophoresis (**Fig. S12B**). Structure prediction models of trimeric assemblies with both ligand-aware models AlphaFold3^39^ and Chai-1^40^ (3 protein chains ± 6 PFOA ligands) suggest various domain swapped oligomer states but have very low confidence scores (pTM scores between 0.2 and 0.4 for all apo and holo models) and thus are not reliably interpretable (**Fig. S13**).

To better understand the designability of the LLM-generated candidates, we queried the backbone of LPB-hex-175 against the non-redundant PDB60 database with TM-align^58^ to assess the prevalence of the fold. We filtered hits for TM-score > 0.5 (indicative of same global fold) and query coverage > 0.7, affording 343 hits out of 45,177 non-redundant structures. Of these, no structures had RMSD < 3.0 Å. Thus, the backbone of the LLM-generated designs is not a prevalent structure in the PDB. We next compared the LLM-generated sequence space to that explored by structure-based methods for the same backbone. To this end, we used the model of the lead candidate, LPB-hex-175, as input for LigandMPNN re-design with a moderate temperature (t = 0.5) to sample more sequence diversity. MPNN-generated designs sample different sequence space than those generated by the LLMs (**Fig. S14**), showing that the LLMs can enforce sequence-based design principles that may not be inherently favored in structure-based methods. We then refolded the LigandMPNN redesigned sequences with Chai-1, affording higher confidence models relative to the input design LPB-hex-175 (average pTM of re-designed models = 0.834 and 0.537, respectively; LPB-hex-175 pTM = 0.700, ipTM = 0.356). The top scoring redesign (pTM = 0.929, ipTM = 0.702) had a Cα RMSD of 0.97 relative to LPB-hex-175 but only 26% sequence identity (51/194 residues conserved). As with the LMPB designs, the LLM-generated designs are feasible but not optimal. Overall, with only two iterative natural language prompts (<250 words total) and 14 designs tested (6 LPB-pent designs and 8 LPB-hex designs), we obtained a lead candidate that expresses, has helical secondary structure, binds the desired small molecule target and exhibits different biochemical features than those preferred by structure-based methods alone.

## Discussion

The field of protein design seeks to derive rules from natural proteins to enable the design of new-to-nature proteins from first principles. While structure-based generative pipelines have enabled increasingly sophisticated design tasks, these models require a high degree of user expertise at all stages: in the selection of design target, operation of design tools, and evaluation of design candidates. We envision that integration of LLMs will lower this barrier to make design an agentic process wherein the user can articulate a design goal in natural language and obtain design candidates and evaluation metrics with only a few clicks.

While future agentic workflows will integrate structure and sequence-based methods, our goal here was to see how much a reasoned LLM could achieve when used in isolation. We explored two design tasks of varying complexity: the design of metal binding dimeric helical bundles and lipophilic ligand binding proteins. In these tasks, the models must synthesize several chemical and biochemical principles to satisfy chemical and structural restraints articulated in the prompt without explicit prompting for 3D coordinates or structural information. While performance varies by LLM and task, we obtained designs that show the desired structural features and exhibit metal or ligand binding in both design campaigns. In both cases, LLM-designed sequences vary from those generated by structure-based models for the same backbone, indicating that LLMs may sample a less “designable” sequence space than state-of-the-art generative design models. Importantly, the models articulate design rationale in their output, communicating the physicochemical principles that were considered in the process. While the designs generated in the campaigns did not satisfy all requirements delineated in the prompts, this rationale and principles synthesized by the LLMs provides a conceptual foundation on which designs can be evaluated.

To further evaluate the utility of LLMs in *de novo* design campaigns, we prompted the LLMs to evaluate our top designs (selected for experimental validation) in the metal-binding design campaigns (**Supplemental Note 7**). Providing sequence alone, we asked the LLM to identify the likely metal binding residues, metal selectivity, and dimer topology and identify the best candidate. The analysis varies as a function of model sophistication: the more advanced Claude Opus 5 provides a more detailed analysis than Claude Sonnet 4.6, including more reflection on putative 3D arrangements of residues, helix propensity and core packing, and the effect of loop length on hairpin stability and dimer topology. Thus, even since the outset of this study, the ability of LLMs to rationalize design principles has improved dramatically.

We anticipate that protein design will rapidly become more facile when natural language prompts are integrated with structure-aware methods. Indeed, the most recent LLM models have achieved agentic workflows that can run entire structure-based design pipelines to autonomously generate *de novo* protein binders to a number of biological targets.^59^ This effort, however, required a 15,000+ word prompt, a full suite of state-of-the-art generative design models, and $50,000 cloud compute budget with unrestricted LLM tokens. Thus, we are still far from fully autonomous design being widely accessible. Despite this, as the models continue to advance and become more efficient, we envision that LLMs will play a critical role in making design rationale more comprehensible and enabling new design and biological discovery.

## Conclusions

Large language model inference provides an opportunity to synthesize design principles articulated in the literature to generate new-to-nature proteins using natural language prompts alone. The use of LLMs alone lack some of the control permitted in structure-based design campaigns but allow sequence generation from first principles without sophisticated user expertise. Further, LLMs provide detailed justification for principles applied in the design process. We envision that LLMs will continue to increase in utility in *de novo* design, making design campaigns more accessible and comprehensible without user intervention at each design step.

## Acknowledgements

The authors thank Maggie Horst, Ian Bakanas, Sam Mann, and Kehan Chen for thoughtful feedback on the manuscript and members of the DeGrado lab for discussions throughout the project. We thank Rose Yang for assistance in running TMAlign analyses. This work was supported by NIH-R35-GM122603 (W.F.D), NSF-MCB-2306190 (W.F.D), NIH-K99GM155611 (A.K.H), and RS-2023-00252972 (N.H.K).

