## Supporting Information for "*De novo* design of ligand binding proteins using large language models alone"

[#] Authors contribute equally to this work.

| Content | Page |
|---|---|
| **Supplemental materials and methods** | 3 |
| **Supplemental Figures** | |
| Figure S1. Sequence logos for metal binding sequences generated by different ChatGPT models | 6 |
| Figure S2. Confidence metrics for structure prediction of metal binding helical bundles | 7 |
| Figure S3. Analytical size exclusion chromatography traces for LMBP candidates show monodisperse species | 8 |
| Figure S4. Isothermal titration calorimetry of $ZnSO_4$ into LMBP-11 show a LMBP:$Zn^{2+}$ binding affinity in the double-digit μM range | 9 |
| Figure S5. Models of LMBP-11 vs LMBP-12 are predicted to have opposite topologies | 10 |
| Figure S6. Histidine residues designated as metal binding residues are not in binding competent positions in LMBP-11 | 11 |
| Figure S7. ProteinMPNN re-design of top LMBP candidates from each LLM sample different sequence space and have higher confidence metrics than LLM-generated designs | 12 |
| Figure S8. Confidence metrics for LPB-pent structure prediction models | 13 |
| Figure S9. Experimental characterization of LPB-pent designs | 14 |
| Figure S10. LPB-pent designs show minimal change to PFOA resonances in 19F NMR | 15 |
| Figure S11. LPB-hex-175 and LPB-hex-186 have the same overall topology despite low pairwise sequence identity | 16 |
| Figure S12. Size exclusion chromatography and SDS-PAGE of LPB-hex design candidates | 17 |
| Figure S13. Structure prediction models of LPB-hex-175 trimers | 18 |
| Figure S14. LPB-hex backbone redesign and sequence analysis | 19 |
| **Supplemental Tables** | |
| Table S1. Sequences for LLM-generated metal binding proteins for Zn(II) | 20 |

Table S2. Sequences for LLM-generated metal binding proteins for Cu(II) 22

Table S3. Sequences LLM-generated metal binding proteins selection for experimental validation 24

Table S4. Sequences generated for penatmeric LLM-generated PFOA binding proteins (LPB-pent) 25

Table S5. Sequences penatmeric LLM-generated PFOA binding proteins (LPB-pent) selected for experimental validation 27

Table S6. Sequences hexameric LLM-generated PFOA binding proteins (LPB-hex) selected for experimental validation 28

**Supplemental Notes**

Supplemental Note 1. Natural language prompt for metal binding protein design 29

Supplemental Note 2. Output for NLM metal binding protein design, ChatGPT (advanced “thinking” model) 30

Supplemental Note 3. Output for NLM metal binding protein design, Gemini 40

Supplemental Note 4. Natural language prompt for design of PFAS binding proteins 50

Supplemental Note 5. Output for initial PFOA binder prompt, ChatGPT (advanced thinking model) 51

Supplemental Note 6. Gemini output for generation of LPB-pent and LPB-hex designs. 53

Supplemental Note 7. LLM evaluation of LLM-generated metal binding designs 64

Supplemental References 73

#### Methods

Sequence design, structure prediction and sequence selection

Prompts were generated for three different design tasks: metal binding helical bundles, metal binding beta-hairpin peptides, and perfluorooctanoic acid (PFOA) binding helical bundles (Supplemental Notes 1-3). Three different models were queried with identical natural language prompts: ChatGPT (OpenAI; 5 and advanced 5 "thinking" model)[1], Gemini (Google; model 2.5)[2], and Claude (Anthropic; Sonnet 4.5 model)[3]. Models were queried between October 2025 and March 2026. In each prompt, the desired features and constraints were defined, and the model was prompted to generate a specific number of sequences that fit the desired criteria and to explain the rationale for design choices. The designed sequences were scraped from the NLM output and used as input for structure prediction.

All sequences (Supplemental Tables 1-6) were used for structure prediction with AlphaFold3[4,5] and Chai-1[6]. For the metal binding helical bundles, sequences were modeled as dimers both with and without two metal ions (Zn(II) or Cu(II). Beta-hairpins were modeled with one copy of the peptide sequence and one metal ion, and PFOA binders were modeled with the desired equivalents of PFOA using the following SMILES string: C(=O)(C(C(C(C(C(C(C(F)(F)F)(F)F)(F)F)(F)F)(F)F)(F)F)(F)F)O. The confidence scores of each model were extracted and compared across models, and a targeted set of design candidates were selected for experimental validation through a combination of filtering by confidence metric and by manual inspection of the structural models. Sequence space was analyzed by inputting all sequences into ESM2[7], extracting the embeddings, and running a principal component analysis on the ESM2[7] embeddings.

Retrospective analysis of LLM-generated designs with LLMs (Claude Sonnet 4.6 and Opus 5) were performed as incognito searches to limit any influence of model memory or context on evaluation of the designs.

Redesign of top candidates

To assess the designability of the backbones of LLM-generated design candidates, the structure prediction models of top candidates were used as input for redesign with ProteinMPNN[8] and/or LigandMPNN[9]. 32 sequences were designed per backbone and refolded with AlphaFold2 (apo structure) or Chai-1 (holo structure). Sequence space was analyzed by inputting all sequences (LLM-designed and MPNN redesigned) into ESM2, extracting the embeddings, and running a principal component analysis on the ESM2 embeddings.

Sequence and structural similarity search

Sequence similarity was assessed via BLASTp[10,11] search and by sequence search in the Protein Data Bank. Structural similarity was assessed by TM-align[12] search against a non-redundant PDB60 database. For TM-align, candidates were filtered by TMscore (>0.5) and query coverage (>0.7) and assessed by backbone RMSD.

Gene fragment design and plasmid construction

The amino acid sequences for metal binding helical bundles and PFOA binders were appended with a N-terminus TEV protease cut site and hexahistidine tag, then back translated to nucleotide sequences and codon optimized for expression in *Escherichia coli.* Homology arms for Gibson assembly were appended onto the nucleotide sequence and gene fragments were ordered from Twist Bioscience. Upon receipt, gene fragments were resuspended and cloned into a linearized pET28a vector via Gibson assembly with 20 ng insert, and 20 ng vector. Following 1h incubation at 50°C, Gibson assembly mix was transformed into chemically competent BL21(DE3) *E. coli* by heat shock (30s, 42 °C) and recovered for 1h at 37°C in LB before plating on LB agar plates with kanamycin selection with overnight growth at 37 °C.

Protein expression and purification

Expression cultures (500 mL TB media with kanamycin selection) were inoculated from overnight cultures and grown at 37°C with shaking at 220 rpm to $OD_{600}$ = 0.6 before induction with 0.5 mM isopropyl-β-D-1-thiogalactopyranoside. Upon induction, cultures were grown overnight at 30 °C, 220 rpm. Expression cultures were harvested by centrifugation (400 rpm) and cell pellets were resuspended in 1X phosphate buffered saline (PBS) with additives before lysis by sonication (Sonic Dismembrator 500, Fisher Scientific; 2 sec on, 2 sec off, 40% amplitude, 15 mins total time). Lysate was clarified by centrifugation to remove cell debris (Thermo Scientific LYNX 4000, 16000 rpm for 30 mins) and the protein of interest was purified by immobilized metal affinity chromatography (IMAC) as follows: Lysate was applied to HisPur Ni-NTA resin (Thermo Fisher) pre-equilibrated with 1X PBS with 20 mM imidazole via gravity flow chromatography. Following lysate exposure, the resin was washed with 3 column volumes of 1X PBS with 20 mM imidazole) and hexahistidine tagged proteins were eluted with 7 mL elution buffer (1X PBS with 250 mM imidazole). The hexahistidine tag was cleaved with $His_6$-TEV protease overnight at 4°C and untagged proteins were isolated by reverse IMAC to separate untagged protein from $His_6$-TEV and any uncleaved protein. Initial protein purity was assessed by SDS-polyacrylamide gel electrophoresis using protein samples prepared in 4X NuPAGE LDS sample buffer (Thermo Fisher). Samples were heated to 95°C for 5 min before loading into NuPAGE Bis-Tris Mini Protein Gel (Thermo Fisher) and running at 200V for 30 min in 1X MES SDS running buffer (Thermo Fisher). Gels were stained with SimplyBlue SafeStain Coomassie gel stain (Thermo Fisher).

Proteins were purified with a final size exclusion chromatography step using a Superdex 75 10/300 column on an AKTA fast protein liquid chromatography system (1X PBS, 0.5 mL/min). Fractions containing the protein of interest were pooled and concentrated using Amicon centrifugal filter device (3 kDa MWCO). For metal binding helical bundles, an additional purification step was added in which the proteins were concentrated and purified by reverse phase preparative high performance liquid chromatography (HPLC) to ensure no trace metals were bound to the protein. HPLC-purified samples were then lyophilized and resuspended in Chelex-treated 10mM MOPS pH 7.5, 100 mM NaCl before downstream analysis.

Circular dichroism

*Metal binding helical bundles*

Following HPLC purification and lyophilization, purified proteins were resuspended in 10 mM 3-(N-morpholino)propanesulfonic acid (MOPS) pH 7.5 with 100 mM NaCl. Stock concentration was determined by UV-Vis spectroscopy and samples were prepared for circular dichroism at a final concentration of 10μM (250μL sample). All designs were screened for secondary structure features in a 0.1 mm cuvette using a JASCO J-810 spectrophotometer using the following settings for spectrum measurement: scan from 250-190nm, 8sec response, 50nm/min scanning speed, 1nm bandwidth, 6 accumulations.

For the samples that showed helical secondary structure, the samples were subsequently incubated with 4 equivalents of ZnSO4 or CuSO4 (prepared in water at 5mM concentration and diluted into the protein sample to a final concentration of 40μM). Samples (apo and holo) were then subjected to thermal denaturation (SETTINGS) to assess protein stability and metal binding-induced stabilization.

*PFOA binders*

PFOA binder concentration was measured by UV-Vis spectroscopy and samples were prepared to a final concentration of 10μM in 1X PBS prior to CD spectrum measurement using the settings reported above.

*Data analysis*

For all CD data, mdeg was converted to molar ellipticity using the following equation: (mdeg*M)/(10*L*C) where M is the average molecular weight of the amino acids in the protein (in g/mol), C is the concentration of the protein in g/L and L is the path length of the cell. All data was plotted in GraphPad (Prism).

<u>Analytical size exclusion chromatography</u>

The oligomeric state of all proteins was assessed by analytical size exclusion chromatography using a Superdex 75 5/150 on an AKTA fast protein liquid chromatography system (50μM sample, 50uL, in 1X PBS, running at 0.2 mL/min).

<u>Isothermal titration calorimetry</u>

Isothermal titration calorimetry was performed on a Malvern MicroCal PEAQ-ITC instrument. To prepare samples for ITC, reverse-phase HPLC purified LMBP-11 was lyophilized and reconstituted in Chelex-treated buffer (10mM MOPS, pH 7.4 with 50 mM NaCl) before extensive dialysis against the same buffer. Zinc stocks were prepared by dissolving $ZnSO_4$ in dialysis buffer after protein dialysis was completed. Protein concentration was confirmed by UV-Vis using the following equation as the design does not have A280 signal: 144(A215–A225). Protein was diluted to the desired concentration and a final volume of 280μL was loaded into the ITC sample cell. Independent titration experiments of $ZnSO_4$ into LMBP-11 were performed (1 x 0.4μL injection followed by 34 x 1μL injections), with [Zn(II)] determined for each experiment with a UV-vis assay using spectrophotometric probe 4-(2-pyridylazo)resorcinol (10mM MOPS, pH 7.4, using reported extinction coefficients)[13]. Binding isotherms were fit using a one set of site model with fitted offset correction. Ligand-into-buffer control titrations (1 x 0.4μL injection followed by 34 x 1μL injections) were performed to assess heats of dissolution during ligand titration.

<u>19F NMR to assess protein:perfluorooctanoic acid interactions</u>

LLM-generated perfluorooctanoic acid binding designs were screened by 19F NMR to assess perfluorooctanoic acid (PFOA) binding. Proteins were purified as described above and protein concentration was determined via UV-Vis spectroscopy. NMR samples were prepared in 1× PBS containing 5% $D_2O$, with a final PFOA concentration of 150 μM. Proteins were added in either 0.2 or 0.5 molar equilvalents and incubated for 2 hrs before analysis. All NMR spectra were acquired at 298.1 K using a Bruker Avance III HD 400 MHz spectrometer and processed using TopSpin 3.6.3 (Bruker).

#### Supplemental Figures

chatGPT advanced “thinking” model

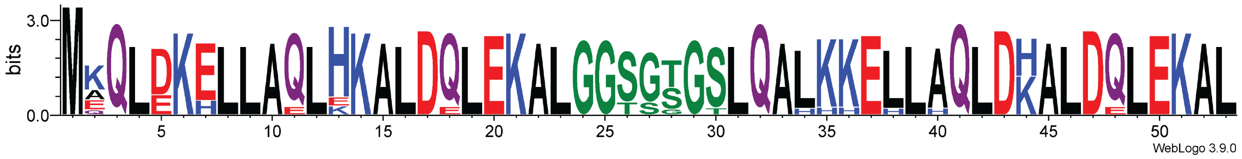


chatGPT (free base model)

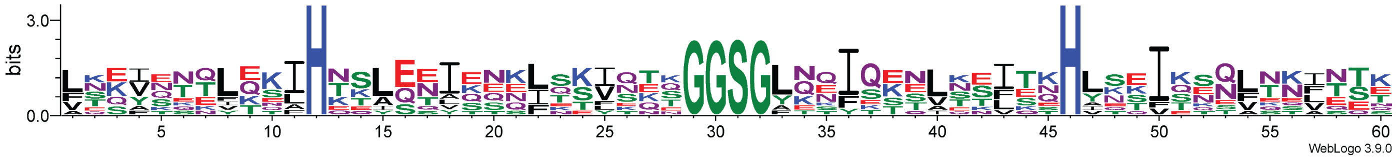


**Figure S1. Sequence logos for metal binding sequences generated by different ChatGPT models.** Sequence logos show that the more advanced ChatGPT model (top) converged on a highly similar sequence pattern, while the free base model (bottom) sampled more sequence diversity.

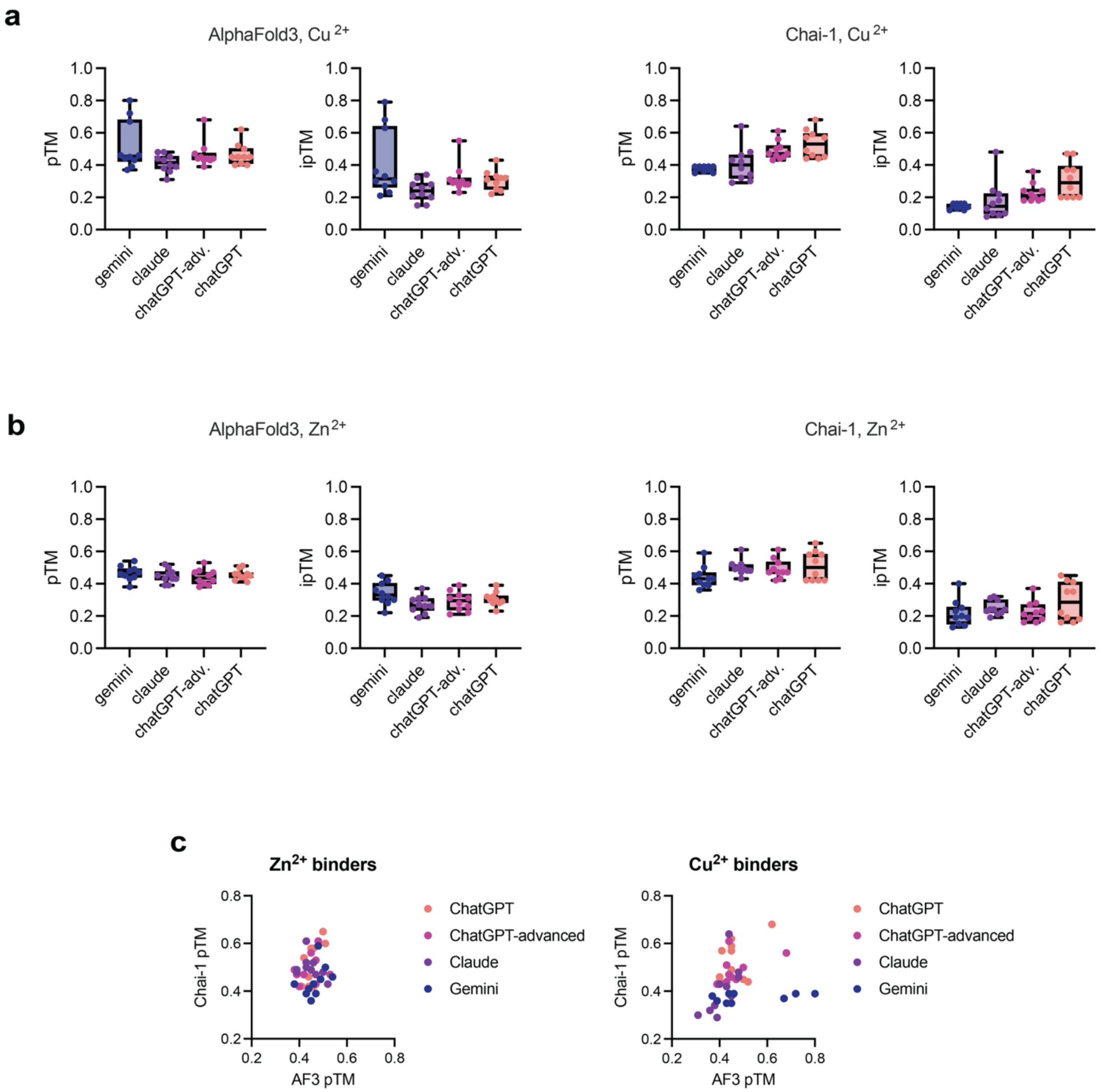


**Figure S2. Confidence metrics for structure prediction of metal binding helical bundles.** Confidence metrics (pTM and ipTM) for AlphaFold3 and Chai-1 metrics for (A) Cu2+ and (B) Zn2+ binding metals designed by ChatGPT, ChatGPT (advanced “thinking” model), Claude, and Gemini. Chai-1 metrics are also presented in Figure 2B as pTM vs ipTM plots. (C) comparison of average pTM per design in AF3 vs Chai-1 predictions show that the structure prediction models have varying confidences across designs.

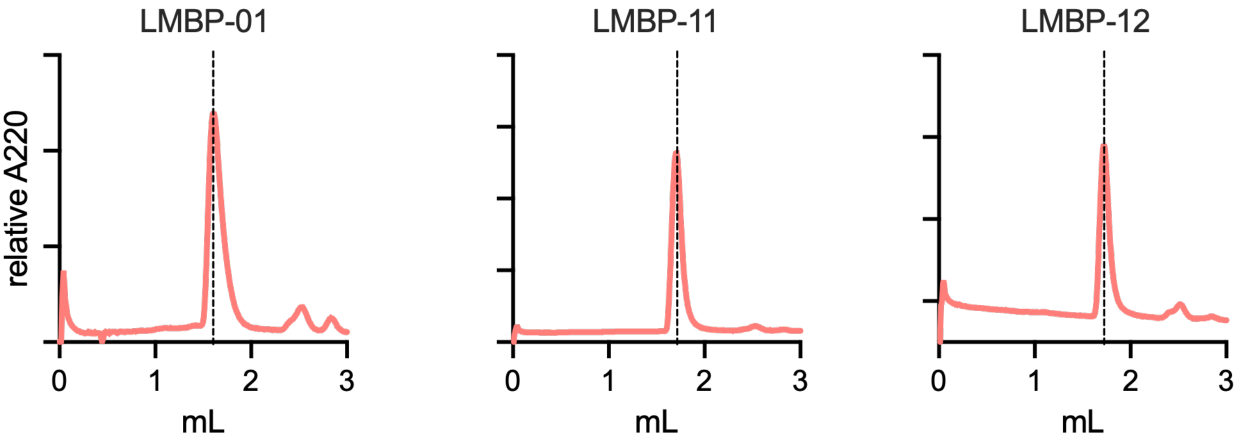


**Figure S3. Analytical size exclusion chromatography traces for LMBP candidates show monodisperse species.**

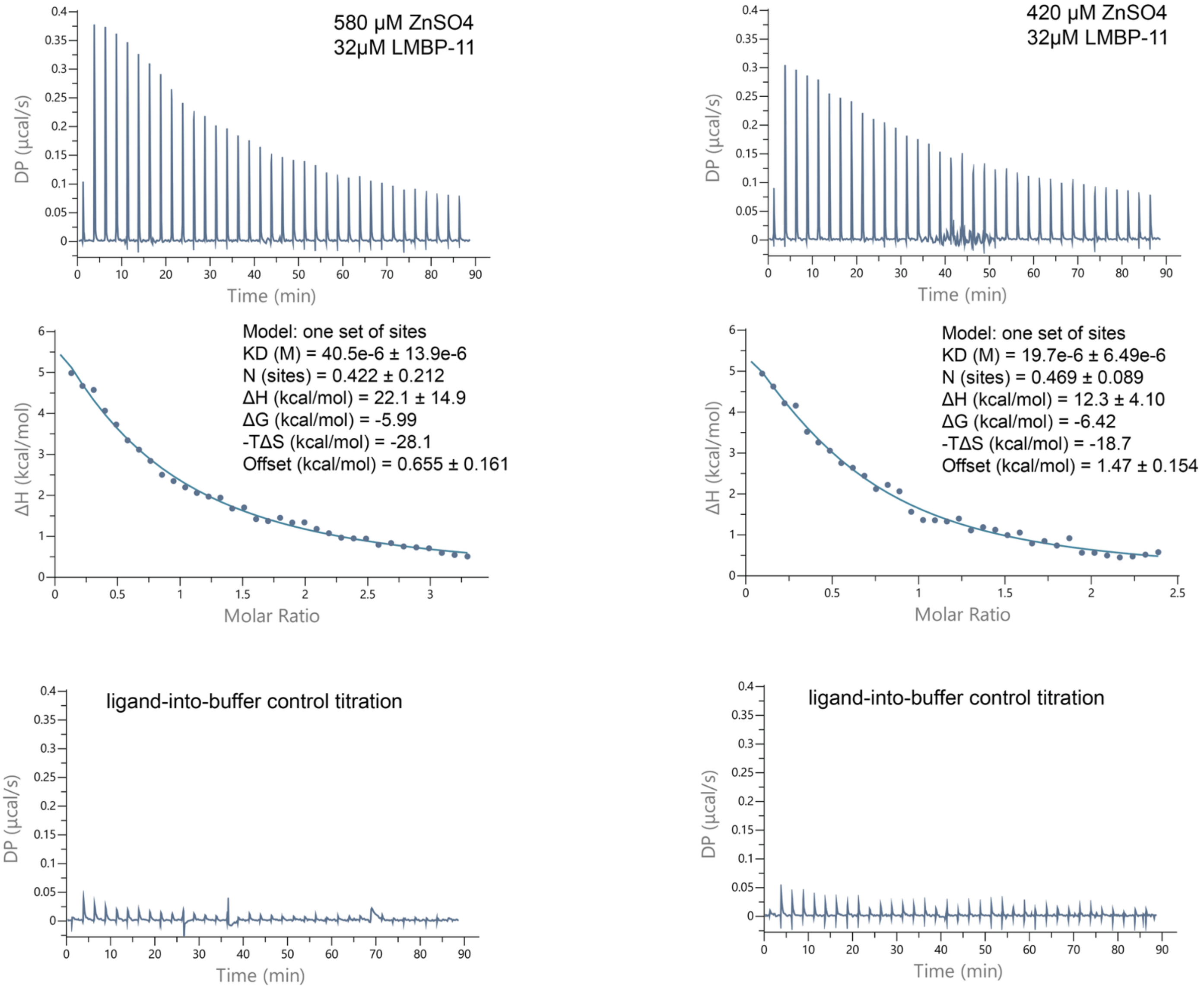


**Figure S4. Isothermal titration calorimetry of $ZnSO_4$ into LMBP-11 show a LMBP:$Zn^{2+}$ binding affinity in the double-digit µM range.** Experiments were performed with 35 injections (1 x 0.4µL initial injection followed by 34 x 1 µL injections) of ligand into protein (280µL sample cell). Concentration of protein and ligand were independently determined for each replicate, with 580µM and 420µM ZnSO4 titrated into 32µM protein for experiments presented on the left and right, respectively. Top: Thermograms for each experiment; middle: binding isotherms (normalized heat vs molar ratio of ligand); bottom: thermograms for ligand-into-buffer control titrations. Binding affinities calculated for each experiment are 40.5 µM ± 13.9 µM and 19.7µM ± 6.49 µM for left and right experiments, respectively, with an average $K_D$ of 30.1 µM ± 7.67 µM across experiments.

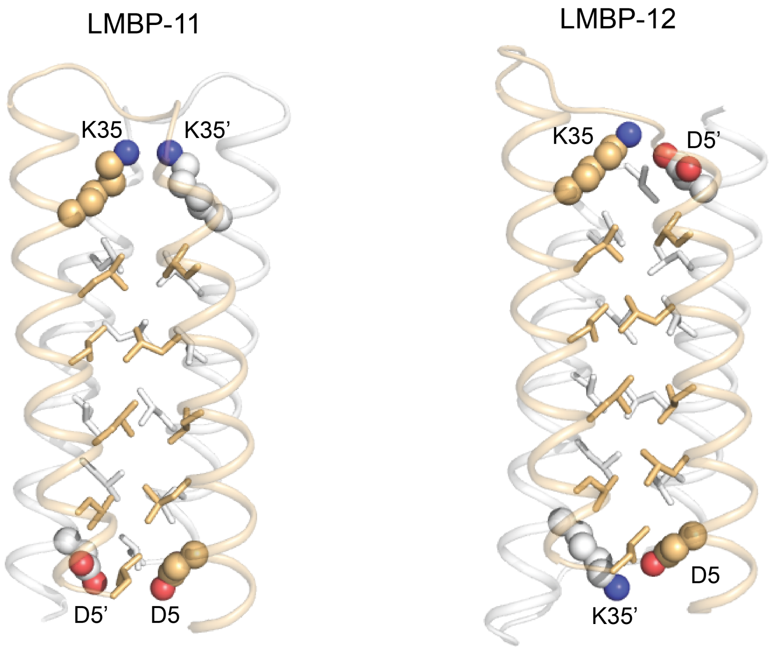


```
         abcdefgabcdefgabcdefga        abcdefgabcdefgabcdefga
LMBP-11 MEQLDKELLAQLHKALDELEKALGGSGTGSLQAHKKELLAQLDKALDQLEKAL
LMBP-12 MQQLDKELLAQLHKALDQLEKALGGSGTGSLQALKHELLAQLDKALDQLEKAL
        *:***************:*************** *:*****************
            ↑                             ↑
            D5                            K35
```

**Figure S5. Models of LMBP-11 vs LMBP-12 are predicted to have opposite topologies.** Models of LMBP-11 and LMBP-12 (holo models, predicted with Chai-1, but $Zn^{2+}$ omitted for clarity) shown as cartoon ribbons, with each chain of the dimer colored in wheat and white, respectively. Side chains are shown as sticks or spheres for residues in *a* and *d* positions of the heptad repeat, forming the the core of the helix-turn-helix dimers. LMBP-11 is predicted to form a parallel dimer of helix-turn-helix hairpins, while LMBP-12 is predicted to form an anti-parallel dimer topology. As a result, the LMBP-12 model indicates salt bridges between D5 and K35’ on opposite chains of the dimer. These interchain salt bridges would not be formed in the LMBP-11 topology. D5 and K35 in each chain highlighted in sphere representation.

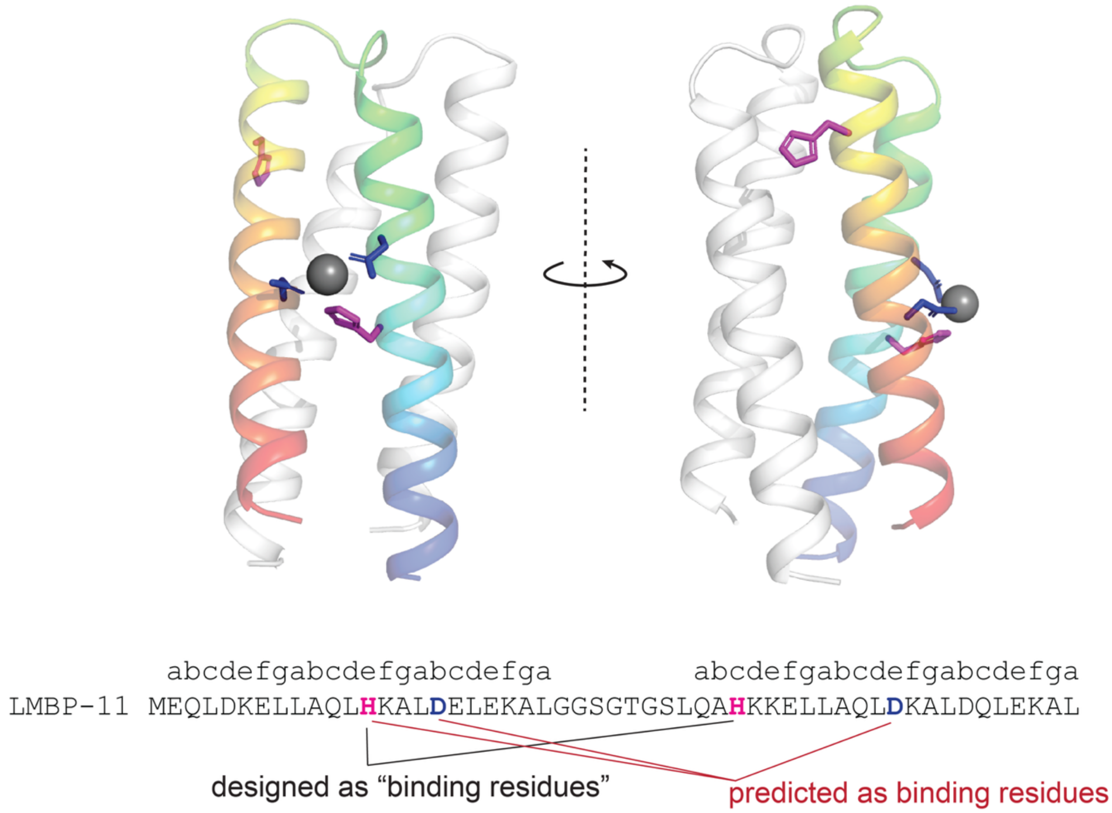


**Figure S6. Histidine residues designated as metal binding residues are not in binding competent positions in LMBP-11.** Model of LMBP-11 (chain A in chainbow, chain B in white), with designed “binding” histidine residues (e.g. the residues the LLM denotes as part of the binding interaction) shown in pink sticks and additional residues participating in predicted metal binding site (as determined in Chai-1 prediction) in blue sticks. One of the histidine residues designated by the LLM as part of the binding site is separated from the predicted binding site by ~2 helical turns and is on the opposite face of the helix-turn-helix hairpin.

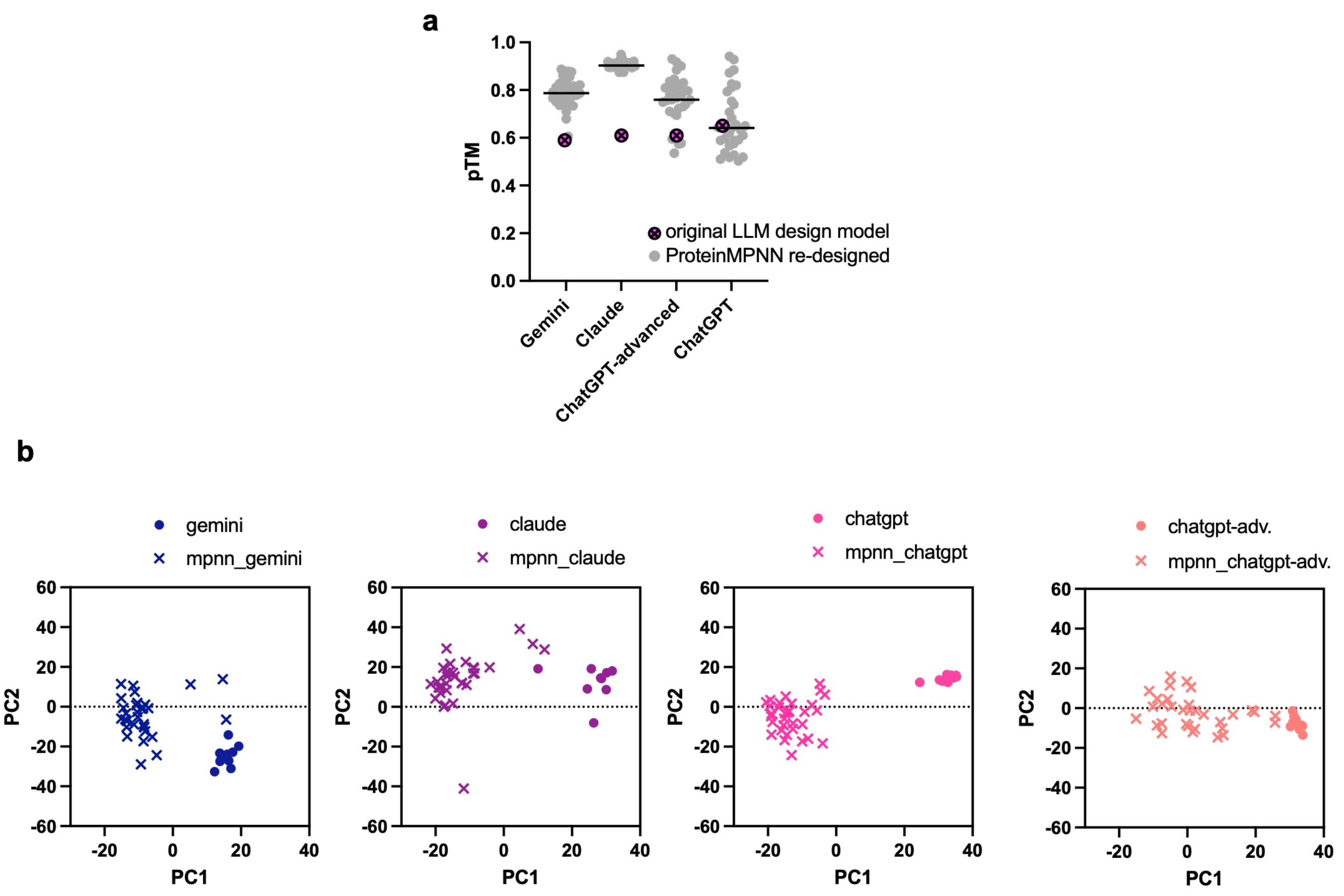


**Figure S7. ProteinMPNN re-design of top LMBP candidates from each LLM sample different sequence space and have higher confidence metrics than LLM-generated designs.** (A) confidence metrics for AlphaFold models of designs generated by subjecting the LMBP design candidate from each LLM output to redesign with ProteinMPNN. (B) Sequence space of ProteinMPNN redesigns is compared to that of original LLM designs via principal component analysis of ESM2 embeddings. Embeddings and PCA are computed across all sequences (all LLM designed sequences and all ProteinMPNN redesigned sequences) but distributions are shown by LLM for clarity.

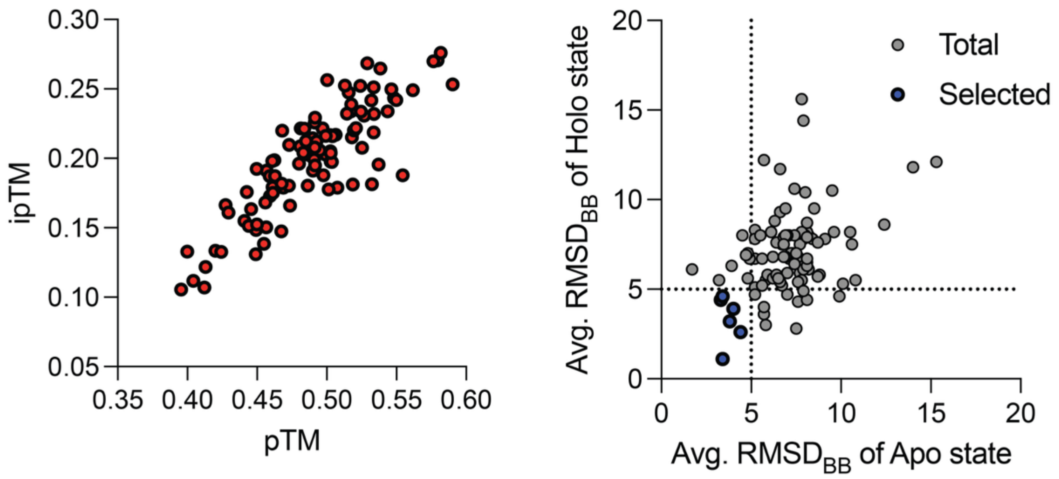


**Figure S8. Confidence metrics for LPB-pent structure prediction models.** Structure prediction of LPB-pent design candidates with Chai-1 gave low to moderate confidence models (left). Design candidates were filtered by self-consistency metrics in both apo and holo state, affording six designs for experimental validation.

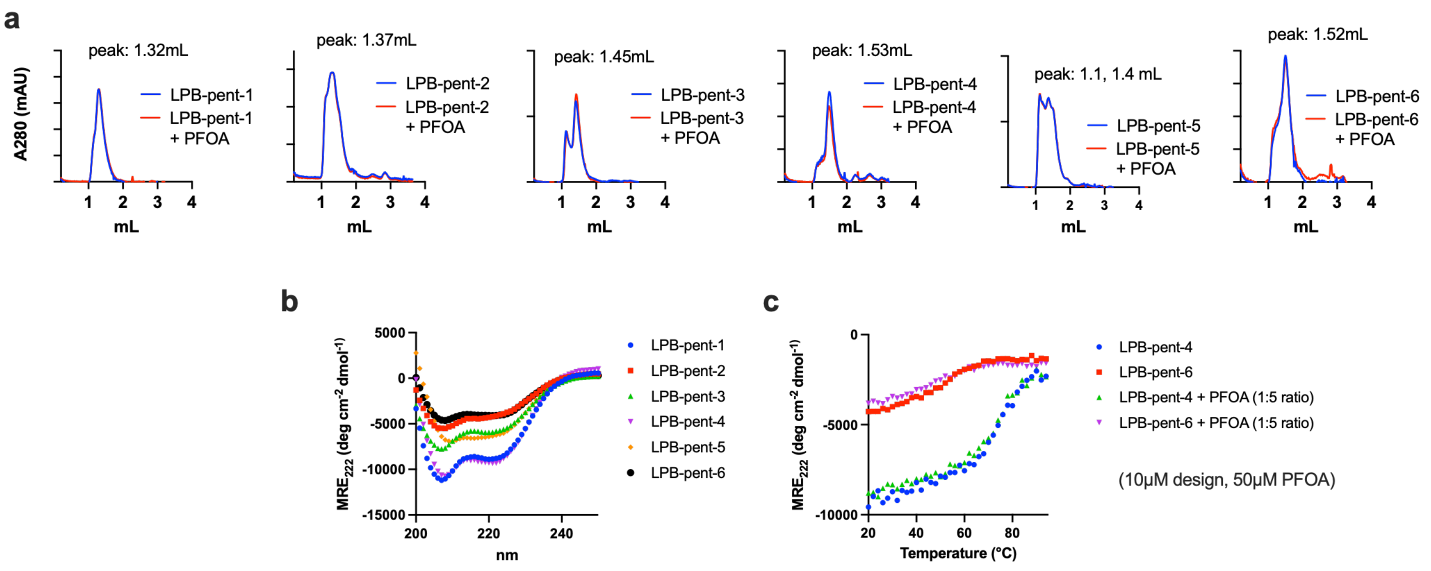


**Figure S9. Experimental characterization of LPB-pent designs**. (a) Analytical size exclusion chromatography traces for LPB-pent 1-6 in the presence and absence of excess perfluorooctanoic acid. (b) Far-UV circular dichroism spectra for LPB-pent 1-6 show that all designs have helical character. (c) LPB-pent-4 and LPB-pent-6, which showed the best monodisperse character by SEC, do not show PFOA-dependent changes in thermal stability.

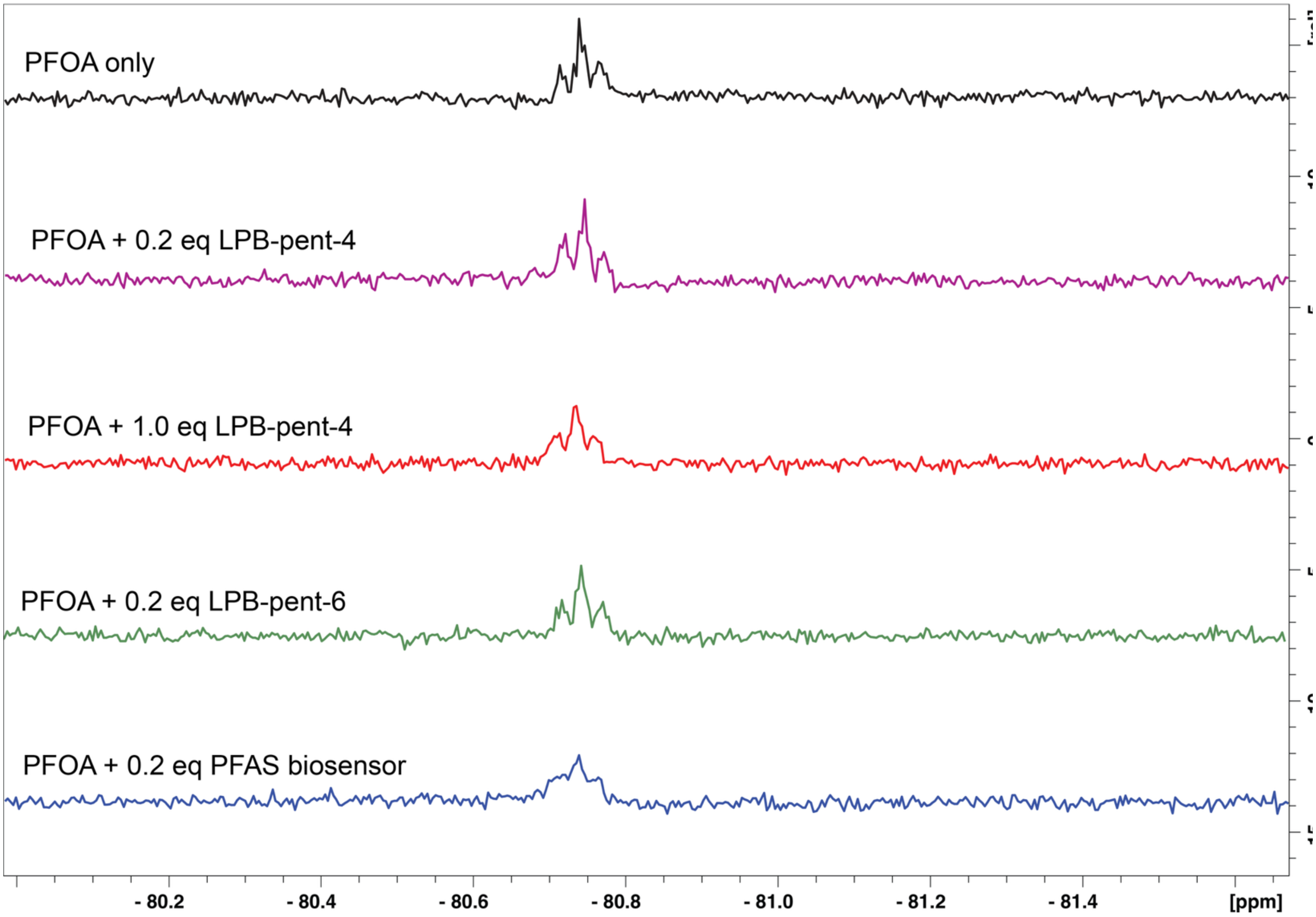


**Figure S10. LPB-pent designs show minimal change to PFOA resonances in 19F NMR.** PFOA was titrated with sub- or stoichiometric equivalents of LPB-pent-4, LPB-pent-6, and a previously reported PFAS biosensor[14] (positive control). Protein:PFAS binding should shift or broaden PFOA resonances[15] (shown here at ~ –81 ppm, corresponding to the terminal trifluoromethyl group), but LPB-pent designs show minimal broadening at the equivalents tested.

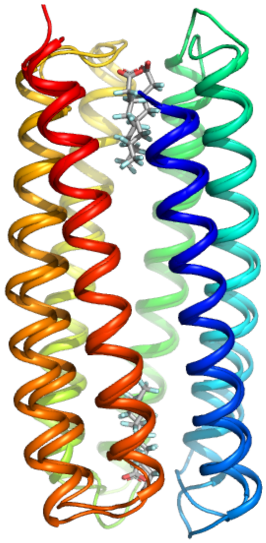

**Figure S11. LPB-hex-175 and LPB-hex-186 have the same overall topology despite low pairwise sequence identity**. Alignment of structure prediction models of LPB-hex-175 and LPB-hex-186 (holo models generated with Chai-1) give a Cα RMSD = 1.8Å. Both models colored in chainbows with PFOA ligands in gray. Models are aligned with the cealign algorithm in PyMol.

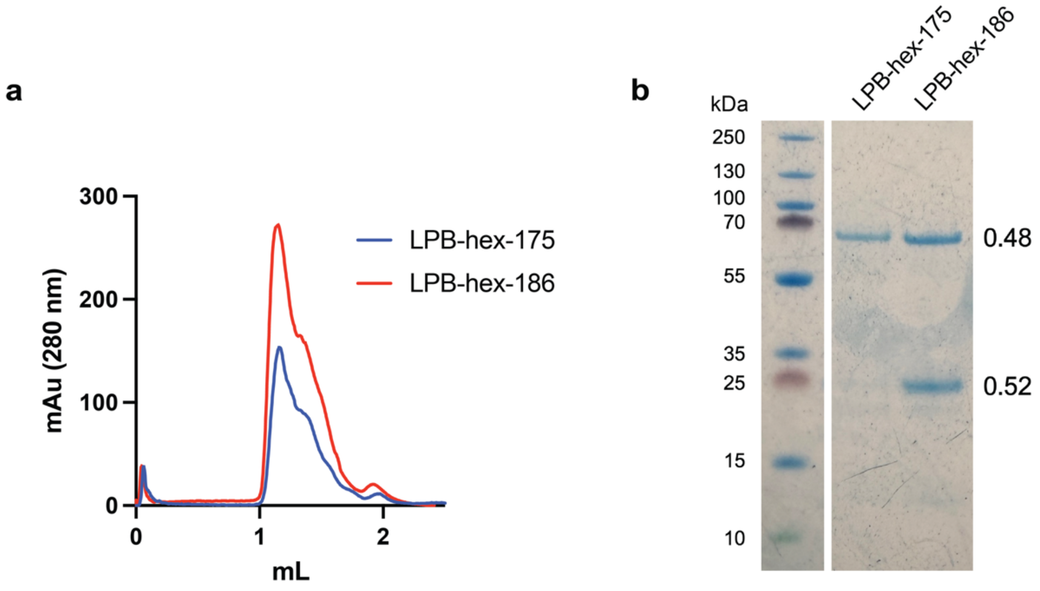


**Figure S12. Size exclusion chromatography and SDS-PAGE of LPB-hex design candidates**. Six-helix bundle designs LPB-hex-175 andLPB-hex-186 are (a) not monomer disperse by analytical size exclusion chromatography and (b) form trimers that persist even after boiling in LDS loading buffer (expected monomer MW ~25kDa). LPB-hex-175 persists as a trimer, while LPB-hex-186 forms a mixture of trimer and monomer in SDS-PAGE.

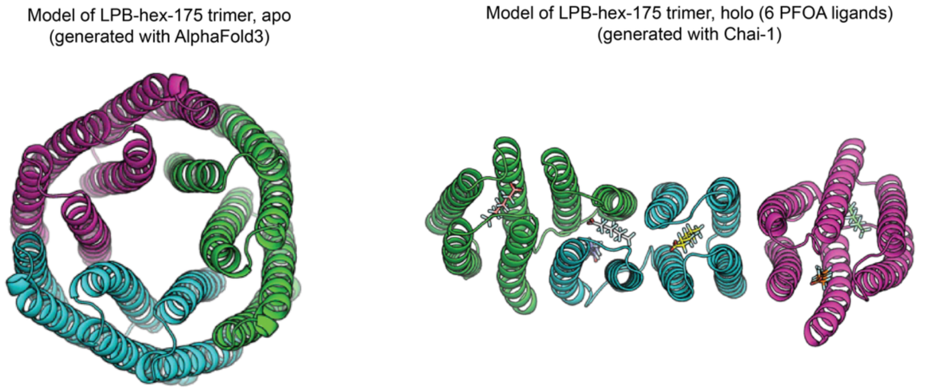


**Figure S13.** Structure prediction models of LPB-hex-175 trimers in apo and holo states generated with AlphaFold3 and Chai-1, respectively.

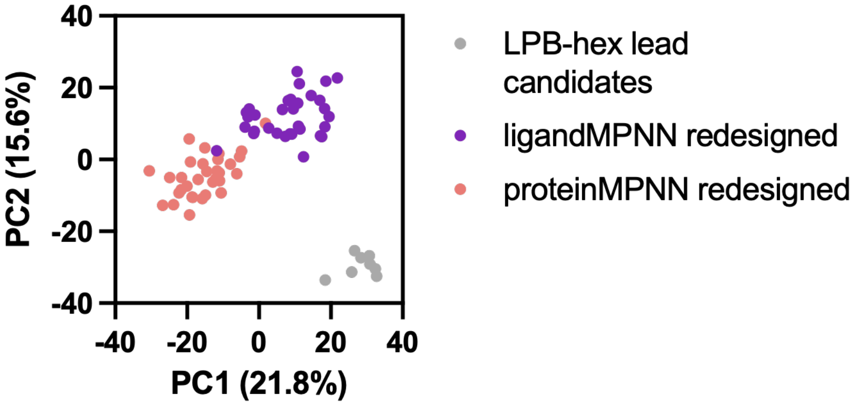


**Figure S14. LPB-hex backbone redesign and sequence analysis.** Principal component analysis of ESM2 embeddings for LPB-hex designs compared to LigandMPNN and ProteinMPNN re-design of lead LPB-hex-175 backbone

**Table S1. Sequences for LLM-generated metal binding proteins for Zn(II), labeled by LLM**

| id | sequence |
|---|---|
| gemini_1 | LEKLEEKHLEKIQEKLEKIQEKHLEEKIGGPGGPGGKELEKIHKEQLEKIKEALEKIKEKHLEEKI |
| gemini_2 | AEKAEEKAEKIQEKAEKIQEKAEEKAGGPGSGPGGEAEKAHKEQAEKIKEQAEKIKEKAEKA |
| gemini_3 | FEKFEEKFEKIQEKFEKIQEKFEEKFGGPGSGPGGEFEKFHKEQFEKIKEQFEKIKEKFEEKF |
| gemini_4 | LEKLEEKHLEKIQEKLEKIQEKHLEEKIGGPGAPGAGKELEKIHKEQLEKIKEALEKIKEKHLEEKI |
| gemini_5 | IEKLIEKHIEKQEEIIEKQEEHIEKLIIGGPGSGPGGEIEKIHKEQIEKKQEQIEKKQEKHIEKII |
| gemini_6 | VEKVEEKVVEKIQEKVEKIQEKVEEKVGGPGSGPGGEVEKAHKEQVEKIKEQVEKIKEKVEEKV |
| gemini_7 | LEKLEEKHLEKIQEKLEKIQEKHLEEKIGGPEGKEGPGLEKIHKEQLEKIKEALEKIKEKHLEEKI |
| gemini_8 | FEKLEEKHFEKIQEKFEKIQEKHLEEKIGGPGSGPGGEFEKIHKEQFEKIKEAFEKIKEKHLEEKI |
| gemini_9 | LEKYEEKHLEKIQEKLEKIQEKHLEEKIGGPGSGPGGELEKIHKEQLEKIKEALEKIKEKHLEEKI |
| gemini_10 | AEKAEEKHAEKIQEKAEKIQEKHLEEKIGGPGSGPGGEAEKIHKEQAEKIKEAAEKIKEKHLEEKI |
| claude_1 | MAELIAKEQEIKHVALEKHQSGKDELLARIAVEIHEALEIAHKDLEELKAQLEEALE |
| claude_2 | MVEVIAQLEQIHEALKEHQSGPDKLLVEVAQEIHKALEFAHKDLEQLVKQLEEAL |
| claude_3 | MAELIAKLEQELHKVADALEHKQSGDETKLLAEIVKQLHEALQIAHKDLEELKAQLELQAL |
| claude_4 | MKELIAYEQRIHEALEKHQSGKDKLLFDLVQQIHEALKFAHKDLEELKAQLEQALE |
| claude_5 | MAEVIAKVEQILHEIAQELHQSGPEELLVRIAQEIHEALEVAHKDLEELVKQLEQA |
| claude_6 | MKELIAFLEQRIHQALEKHQDSGETKLLFEIVQRLHDALQFAHKDLEELKAQLEQALE |
| claude_7 | MAELIAKLEQELKHVADALEVHQKSGDPEKLARIQQELDALNVVHEALKELHKQALEQAL |
| claude_8 | MKELIAELEKRIHEALKEHKDSGKDELLAKIVEEIHKALEIAYKDLEELKAQLEQALE |
| claude_9 | MAELIAELEQALHEALKEHQSGKDKLLAEIAQELHEALEIAHKELEQLKAELEQAL |
| claude_10 | MKELIAFLEQELHKVAQELHKDSGKEELLARIAQEIHDALQFAYKDLEELKAQLEQALE |
| chatgpt_1 | LKEINNTVKNAHNSAETLKEKLSKVNTKGGSGLNNIQTQVKNFKQHYKTITSNFTQINSN |
| chatgpt_2 | FNQYNTQITTIHTSASNIKNKISSIQQSGGSGLQSISESLETFTKHYQSIKNEFSNVNEK |
| chatgpt_3 | FSEVQTQFKSIHKKLEQISQELQEFTETGGSGLTEIKENLQEITSHLKQIKTQVTNFTTS |
| chatgpt_4 | LTKVETTLEQLHKKIETIEQKFKSLSNQGGSGYKNIQSTVNKISNHINEIESQLETASET |
| chatgpt_5 | LSQAKQQLQEIHEELQSYQSSLNKVQQEGGSGANKFSKEISSIEKHLTKIKQQLKNLQTK |
| chatgpt_6 | VNSITNELEKIHTSLENIESNLSSIEEQGGSGLQQIEKNATEIEKHLSEISSSLQKIESK |
| chatgpt_7 | LTKIENNLEKFHQQLEEASEQLTKISTSGGSGYQQIQKTLKTVSEHVNKVQENANSIESQ |
| chatgpt_8 | LEEVQSKLQSIHNTYQEVQTNITTIQKKGGSGFEQFQEELSELTNHASSIQENLTKVTTE |
| chatgpt_9 | VQSYSKELTKLHTTLQNIENQFQKVNKTGGSGLNTIQSNLTSLQQHLEEFTNQLNKLNQE |
| chatgpt_10 | AKKFSETYQKIHNSLSEITNEIETYKTNGGSGLKEYTTSLNSITTHLSTISKTVNEFTTT |
| chatgpt_thinking_1 | MKQLEKHLLAQLEKALDQLEKALGGSGTGSLQALHKELLAQLDKALDQLEKAL |
| chatgpt_thinking_2 | MAQLDKHLLAELKKALDQLEKALGGSGSGSLQALKKELLHQLDKALDQLEKAL |
| chatgpt_thinking_3 | MKQLDKELLAQLHKALDQLEKALGGSSTGSLQALKKEHLAQLDKALDQLEKAL |
| chatgpt_thinking_4 | MEQLDKELLAQLHKALDELEKALGGSGTGSLQAHKKELLAQLDKALDQLEKAL |
| chatgpt_thinking_5 | MKQLDKELLAQLHKALDQLEKALGGSGSGTLQALKKELLAQLDHALDQLEKAL |
| chatgpt_thinking_6 | MQQLDKELLAQLHKALDQLEKALGGSGTGSLQALKHELLAQLDKALDQLEKAL |
| chatgpt_thinking_7 | MKQLDKELLAQLHKALDQLEKALGGTGSGSLQALKKELLAQLDHALDQLEKAL |

| | |
|---|---|
| chatgpt_thinking_8 | MAQLDKELLAQLHKALDQLEKALGGSGTGSLQALKKELLAQLDHALDELEKAL |
| chatgpt_thinking_9 | MKQLEKELLAQLHKALDQLEKALGGSGSGSLQALKKELLAQLDHALDQLEKAL |
| chatgpt_thinking_10 | MEQLEKELLAQLHKALDQLEKALGGTSGGSLQALKKELLAQLDHALDQLEKAL |

#### Table S2. Sequences for LLM-generated metal binding proteins for Cu(II), labeled by LLM

| id | sequence |
|---|---|
| gemini_1 | AEKIEEKHAEKIQEKAEKIQEKHAEEKIGGPGGPGGKEAEKIHKEQAEKIKEAAEKIKEKHAEEKI |
| gemini_2 | AEKAEEKHAEKAQEKAEKAQEKHAEEKAGGPGGPGGKEAEKAHKEQAEKAQEAAEKAQEKHAEEKA |
| gemini_3 | VEKIEEKHVEKIQEKVEKIQEKHVEEKIGGPGSGPGGEVEKIHKEQVEKIKEAVEKIKEKHVEEKI |
| gemini_4 | AEKIEEKHAEKIQEKAEKIQEKHAEEKIGGPGAPGAGKEAEKIHKEQAEKIKEAAEKIKEKHAEEKI |
| gemini_5 | VEKVAEKHVEKVAEKVEKVAEKHVEEKVGGPGSGPGGEVEKAHKEQVEKVAEQVEKVAEKHVEEKV |
| gemini_6 | AEKSEEKHAEKSIEKAEKSIEKHAEEKSGGPGSGPGGEAEKSHKEQAEKSAEQAEKSAEKHAEEKS |
| gemini_7 | AEKIEEKHAEKIQEKAEKIQEKHAEEKIGGPEGKEGPGAEKIHKEQAEKIKEAAEKIKEKHAEEKI |
| gemini_8 | VEKIEEKHVEKIQEKVEKIQEKHVEEKIGGPGSGPGGEAEKIHKEQAEKIKEAAEKIKEKHVEEKI |
| gemini_9 | IEKAEEKHIEKAQEKIEKAQEKHIEEKAGGPGGPGGKEIEKIHKEQIEKAQEAIEKAQEKHIEEKA |
| gemini_10 | AEKAEEKHAEKAQEKAEKAQEKHAEEKAGGPGGPGGKEAEKAHKEQAEKAQEAAEKAQEKHAEEKA |
| claude_1 | MSEIAALESQLKHADQLLEKESGPGDKIHAELAKVLESHIAELEIAQQKLESELKG |
| claude_2 | MQLIAEELSKQAHINALKQSEGNPSDAHIEQLVAKLQSHLAELEVAKNLQAELKS |
| claude_3 | MSEILAKELQHAIQNLKESDHGPGQSHIQELAANLESHVAELEIAKQVLQKELQSKG |
| claude_4 | MAQILAELSKQHAINQLKESDGPGERHLAEIANKLESHIPELEIAKVLQSELQAKG |
| claude_5 | MSQILAEQLKAHIEQLNKSDGPGDKHIQELAAKLESHVAELEVAKQILQKELNAELKG |
| claude_6 | MAEILKQLSQHAIENLIKQDGPGSKHLQEIAKVLANHIEELEVAKILQQELQSKG |
| claude_7 | MSELIAKQLQEHAIKLNQSDGPGDRHIQELAANVLKHISELEIAKQVLQAELKSKG |
| claude_8 | MAQILAEELSKHAINQLQESDGPGQRHLAEIAVKLNHIPELEIAKQVLQSELKAKG |
| claude_9 | MSEILAKQLQSHAIENLKQSDGPGEKHLQELAAVLKSHINELEIAQQVLQAELKSEKG |
| claude_10 | MAELIAQKLSQHAINELKQSDGPGSKHIQELAAKVLNHISELEVAKILQQELQSKKG |
| chatgpt_1 | LKEINNTVKNAHNSAETLKEKLSKVNTKGGSGLNNIQTQVKNFHQTYKTITSNFTQINSN |
| chatgpt_2 | FNQYNTQITTIHTSASNIKNKISSIQQSGGSGLQSISESLETFHKNYQSIKNEFSNVNEK |
| chatgpt_3 | FSEVQTQFKSIHKKLEQISQELQEFTETGGSGLTEIKENLQEIHSQLKQIKTQVTNFTTS |
| chatgpt_4 | LTKVETTLEQLHKKIETIEQKFKSLSNQGGSGYKNIQSTVNKIHNKINEIESQLETASET |
| chatgpt_5 | LSQAKQQLQEIHEELQSYQSSLNKVQQEGGSGANKFSKEISSIHKKLTKIKQQLKNLQTK |
| chatgpt_6 | VNSITNELEKIHTSLENIESNLSSIEEQGGSGLQQIEKNATEIHKELSEISSSLQKIESK |
| chatgpt_7 | LTKIENNLEKFHQQLEEASEQLTKISTSGGSGYQQIQKTLKTVHESVNKVQENANSIESQ |
| chatgpt_8 | LEEVQSKLQSIHNTYQEVQTNITTIQKKGGSGFEQFQEELSELHNSASSIQENLTKVTTE |
| chatgpt_9 | VQSYSKELTKLHTTLQNIENQFQKVNKTGGSGLNTIQSNLTSLHQELEEFTNQLNKLNQE |
| chatgpt_10 | AKKFSETYQKIHNSLSEITNEIETYKTNGGSGLKEYTTSLNSIHTELSTISKTVNEFTTT |
| chatgpt_thinking_1 | MKQLEKHLLAQLEKHLDQLEKALGGSGTGSLQALKHELLAQLDHALDQLEKAL |
| chatgpt_thinking_2 | MAQLDKHLLAELKHALDQLEKALGGSGSGSLQALHKELLAQLDHALDQLEKAL |
| chatgpt_thinking_3 | MKQLDKHLLAQLKHALDQLEKALGGSSTGSLQALHKELLAQLDHALDELEKAL |
| chatgpt_thinking_4 | MEQLDKHLLAQLKHALDELEKALGGSGTGSLQALHKELLAQLDHALDQLEKAL |
| chatgpt_thinking_5 | MKQLDKHLLAQLKHALDQLEKALGGSGSGTLQALHKELLAQLDHALDQLEKAL |
| chatgpt_thinking_6 | MQQLDKHLLAQLKHALDQLEKALGGSGTGSLQALHKELLAQLDHALDQLEKAL |
| chatgpt_thinking_7 | MKQLDKHLLAQLKHALDQLEKALGGTGSGSLQALHKELLAQLDHALDELEKAL |

| | |
|---|---|
| chatgpt_thinking_8 | MAQLDKHLLAQLKHALDQLEKALGGSGTGSLQALHKELLAQLDHALDQLEKAL |
| chatgpt_thinking_9 | MKQLEKHLLAQLKHALDQLEKALGGSGSGSLQALHKELLAQLDHALDQLEKAL |
| chatgpt_thinking_10 | MEQLEKHLLAQLKHALDQLEKALGGTSGGSLQALHKELLAQLDHALDQLEKAL |

**Table S3. Sequences LLM-generated metal binding proteins selection for experimental validation**

| ID | Sequence |
|---|---|
| LMBP-01 | MAELIAKLEQELKHVADALEVHQKSGDPEKLARIQQELDALNVVHEALKELHKQALEQAL |
| LMBP-02 | VNSITNELEKIHTSLENIESNLSSIEEQGGSGLQQIEKNATEIHKELSEISSSLQKIESK |
| LMBP-03 | LTKIENNLEKFHQQLEEASEQLTKISTSGGSGYQQIQKTLKTVHESVNKVQENANSIESQ |
| LMBP-04 | VQSYSKELTKLHTTLQNIENQFQKVNKTGGSGLNTIQSNLTSLHQELEEFTNQLNKLNQE |
| LMBP-05 | AKKFSETYQKIHNSLSEITNEIETYKTNGGSGLKEYTTSLNSIHTELSTISKTVNEFTTT |
| LMBP-06 | LTKVETTLEQLHKKIETIEQKFKSLSNQGGSGYKNIQSTVNKISNHINEIESQLETASET |
| LMBP-07 | AEKIEEKHAEKIQEKAEKIQEKHAEEKIGGPGGPGGKEAEKIHKEQAEKIKEAAEKIKEKHAEEKI |
| LMBP-08 | AEKIEEKHAEKIQEKAEKIQEKHAEEKIGGPGAPGAGKEAEKIHKEQAEKIKEAAEKIKEKHAEEKI |
| LMBP-09 | IEKAEEKHIEKAQEKIEKAQEKHIEEKAGGPGGPGGKEIEKIHKEQIEKAQEAIEKAQEKHIEEKA |
| LMBP-10 | MKQLEKHLLAQLKHALDQLEKALGGSGSGSLQALHKELLAQLDHALDQLEKAL |
| LMBP-11 | MEQLDKELLAQLHKALDELEKALGGSGTGSLQAHKKELLAQLDKALDQLEKAL |
| LMBP-12 | MQQLDKELLAQLHKALDQLEKALGGSGTGSLQALKHELLAQLDKALDQLEKAL |

**Table S4. Sequences generated for penatmeric LLM-generated PFOA binding proteins (LPB-pent), labeled by LLM** (initial 10 sequences/LLM, before additional sequences generated with LLM-produced python script)

| ID | sequence |
|---|---|
| claude_pfas_1 | MELEIQKLAEIQDKLEEILEQLKGPRGNELEIQKVAEILEKLQEIWEQLRGSRKNQLEIARLAEIQDKLEEILEKLQGPRGKQLEIQRLAEIQDKLEEILEQLKGSRGNEIEIQKLAEILEKLQEIWQQLK |
| claude_pfas_2 | MQLEIQKLVEILDKLQEVLEQLKGSKGNELEIQKVVEVLDKLQEILEQLRGSKENQLEIARLAEVQDKLQEVLEQLKGSKENQLEVQRLAEVLDKLQEILEQLKGSKGNEIEIQKLVEILDKLQEVWQQLK |
| claude_pfas_3 | MEIEIQKLAFILEKLARILEQIKGGRKNEIEIQKLAFILEKLARILEQIKGGKHNQIEIARLAFILDKLAKILEQIKGGRKNEIEIQRLAFILDKLAKILEQIKGGRKNQIEIQKLAFILDKLWRILEQIK |
| claude_pfas_4 | MELEIQKLAEVLEKVQEILEQLKGPHRNELEIQKVAEVLEKIQEIWEQLRGSHKNQLEIARLAEVLEKVQEILEQLKGPHRNQLEIQRLAEVLEKIQEILEQLKGSHRNQIEIQKLAEVLEKIWQEILEQLK |
| claude_pfas_5 | MEIEIQKFAEILDKLQEILEQIKGSRKNQIEIQKFAEVLDKLQEILEQIKGSKENQIEIARLAFILDKLWEILEQIKGSRKNQLEIQRLAFILDKLQEILEQIKGSKGNQIEIQKFAEVLDKLQEILEQIK |
| claude_pfas_6 | MQIEIQKVAEVIDKVQEILEQIKGSPRNEIEIQKVAEVIDKVQEILEQIKGSPRNQIEIARLAEVIDKVQEILEQIWGSPRNEIEIQRVAEVIDKVQEILEQIKGSPRNEIEIQKVAEVIDKVWEILEQIK |
| claude_pfas_7 | MEIEIQKLAFILEKVQEIWLQIKGGRKNELEIQKLAFILEKVQEILLQIKGGHSNQIEIARLAFILDKVQEILLQIKGGRKNELEIQRLAFILDKVQEILLQIKGGHKNQLEIQKLAFILEKVQEILLWIK |
| claude_pfas_8 | MELEIQKVVEILEKLQEVLEQIKGPKHNEIEIQKVVEVLEKLQEVLEQIKGPKHNQLEIARLAEVLEKLQEVLEQWKGPKHNEIEIQRVAEVLEKLQEVLEQIKGPKHNQIEIQKVVEVLEKLQEVWEQIK |
| claude_pfas_9 | MQLEIQKFAEILDKLQEIWEQIKGPRKNEIEIQKFAEILDKLQEILEQIKGPRKNQIEIARLAFILDKLQEILEQIKGPRKNEIEIQRFAEILDKLQEILEQIKGPRKNQIEIQKFAEVLDKLQEILEQWK |
| claude_pfas_10 | MEIEIQKLAEVLDKVQEILEQIKGSHGNEIEIQKVAEVLDKVQEILEQIKGSHGNQIEIARLVEVLDKVWEILEQIKGSHGNEIEIQRVVEVLDKVQEILEQIKGSHGNQIEIQKLAEVLDKVQEILEQIK |
| gemini_pfas_1 | MGEIQKLEQEIQKLEQAIQKLEQGKGEIQKLEQEIQKLEQAIQKLEQGKGEIQKLEQEIQKLEQAIQKLEQGKGEIQKLEQEIQKLEQAIQKLEQGKGEIQKLEQEIQKLEQAIQKLEQW |
| gemini_pfas_2 | MVEIQKLEQLIQKLEQVIQKLEQPRGVEIQKLEQLIQKLEQVIQKLEQPRGVEIQKLEQLIQKLEQVIQKLEQPRGVEIQKLEQLIQKLEQVIQKLEQPRGVEIQKLEQLIQKLEQVIQKLEQY |
| gemini_pfas_3 | MAIQKLEQLIQKLEQAIQKLEQKRGEIQKLEQLIQKLEQAIQKLEQKRGEIQKLEQLIQKLEQAIQKLEQKRGEIQKLEQLIQKLEQAIQKLEQKRGEIQKLEQLIQKLEQAIQKLEQW |
| gemini_pfas_4 | MIIQKLEQIIQKLEQIIQKLEQGPKIIQKLEQIIQKLEQIIQKLEQGPKIIQKLEQIIQKLEQIIQKLEQGPKIIQKLEQIIQKLEQIIQKLEQGPKIIQKLEQIIQKLEQIIQKLEQY |
| gemini_pfas_5 | MALEQLEQLLKQLEQALKQLEQGKGALEQLEQLLKQLEQALKQLEQGSPKGALEQLEQLLKQLEQALKQLEQGKGALEQLEQLLKQLEQALKQLEQGSPKGALEQLEQLLKQLEQALKQLEQW |
| gemini_pfas_6 | MLLEQKEQLLEQKEQLLEQKEQRGSMLLEQKEQLLEQKEQLLEQKEQRGSMLLEQKEQLLEQKEQLLEQKEQRGSMLLEQKEQLLEQKEQLLEQKEQRGSMLLEQKEQLLEQKEQLLEQKEQY |
| gemini_pfas_7 | MAIQKLEQLLQKLEQAIQKLEQNGKAIQKLEQLLQKLEQAIQKLEQNGKAIQKLEQLLQKLEQAIQKLEQNGKAIQKLEQLLQKLEQAIQKLEQNGKAIQKLEQLLQKLEQAIQKLEQW |
| gemini_pfas_8 | MVEQKLKQLEQKLKQVEQKLKQKPGEVEQKLKQLEQKLKQVEQKLKQKPGEVEQKLKQLEQKLKQVEQKLKQKPGEVEQKLKQLEQKLKQVEQKLKQKPGEVEQKLKQLEQKLKQVEQKLKQY |
| gemini_pfas_9 | MGEIQKQEQIIQKQEQGIQKQEQGRPGEIQKQEQIIQKQEQGIQKQEQGRPGEIQKQEQIIQKQEQGIQKQEQGRPGEIQKQEQIIQKQEQGIQKQEQGRPGEIQKQEQIIQKQEQGIQKQEQW |
| gemini_pfas_10 | MLIQKLEQILQKLEQLIQKLEQSKGLIQKLEQILQKLEQLIQKLEQSKGLIQKLEQILQKLEQLIQKLEQSKGLIQKLEQILQKLEQLIQKLEQSKGLIQKLEQILQKLEQLIQKLEQY |
| chatgpt_pfas_1 | LKALEEKIKALEEKLKAIEEKGDKPRHKIKALEEKLKALEEKVKALEEKGQDPKGLKALEEKLKWLEEKIKALEEKGEPRKQGLKAIEEKVKALEEKIKALEEKGHRPKKGVKALEEKLKYLEEKLKALEEK |
| chatgpt_pfas_2 | IKALEEKLKALEEKLKAIEEKGDKPRHKLKALEEKLKWLEEKVKALEEKGQDPKGLKAIEEKLKALEEKIKALEEKGEPRKQGVKALEEKLKYLEEKIKALEEKGHRPKKGIKALEEKLKALEEKVKALEEK |
| chatgpt_pfas_3 | LKYLEEKLKALEEKLKAIEEKGDKPRHKIKALEEKLKALEEKVKALEEKGQDPKGLKWLEEKLKALEEKLKWLEEKGEPRKQGLKAIEEKVKALEEKIKALEEKGHRPKKGVKALEEKLKALEEKIKALEEK |
| chatgpt_pfas_4 | VKALEEKLKALEEKLKAIEEKGDKPRHKLKALEEKLKYLEEKVKALEEKGQDPKGIKALEEKLKALEEKVKALEEKGEPRKQGLKAIEEKVKALEEKIKALEEKGHRPKKGVKALEEKLKWLEEKLKALEEK |
| chatgpt_pfas_5 | LKWLEEKIKALEEKLKAIEEKGDKPRHKIKALEEKLKALEEKVKALEEKGQDPKGLKALEEKLKYLEEKIKALEEKGEPRKQGLKAIEEKIKALEEKVKALEEKGHRPKKGVKALEEKLKALEEKIKALEEK |

| | |
|---|---|
| chatgpt_pfas_6 | LKALEEKVKALEEKLKAIEEKGKPGRHKIKALEEKLKALEEKVKALEEKGKDPRGLKALEEKLKWLEEKVKALEEKGEPKKHGLKAIEEKVKALEEKIKALEEKGHRPGKGVKALEEKLKYLEEKLKALEEK |
| chatgpt_pfas_7 | IKALEEKLKALEEKLKAIEEKGKPGRHKLKYLEEKLKWLEEKVKALEEKGKDPRGLKAIEEKLKALEEKIKALEEKGEPKKHGLKWLEEKLKYLEEKIKALEEKGHRPGKGVKALEEKLKALEEKIKALEEK |
| chatgpt_pfas_8 | LKWLEEKLKALEEKLKAIEEKGKPGRHKIKALEEKLKALEEKVKALEEKGKDPRGLKYLEEKLKALEEKIKALEEKGEPKKHGLKAIEEKVKALEEKIKALEEKGHRPGKGVKALEEKLKWLEEKLKALEEK |
| chatgpt_pfas_9 | VKALEEKLKALEEKLKAIEEKGKPGRHKLKALEEKLKWLEEKVKALEEKGKDPRGLKAIEEKLKYLEEKIKALEEKGEPKKHGVKALEEKLKALEEKIKALEEKGHRPGKGVKALEEKLKALEEKLKYLEEK |
| chatgpt_pfas_10 | LKYLEEKLKALEEKLKWLEEKGKPGRHKIKALEEKLKALEEKVKALEEKGKDPRGLKWLEEKLKALEEKLKYLEEKGEPKKHGLKAIEEKVKALEEKIKALEEKGHRPGKGVKALEEKLKALEEKIKALEEK |

**Table S5. Sequences penatmeric LLM-generated PFOA binding proteins (LPB-pent) selected for experimental validation**

| ID | Sequence |
|---|---|
| LPB-pent-1 | MVEAVEKQIKKIKAELKELDEQVQAVRQEGGKGGPAQQAKEDISKFDQQIQEIQWEAEEIQSDPGRGSGVAAA RERLAAFEKEIESIEKDVEKAKADGGKGGPVQEVEKKMQQFEADIEKIDEDVAAVQQKGGRRGGVSKVKQKLA SIKEDIKEMQQQVEKVDARE |
| LPB-pent-2 | MIQSVQSRLQKIKEQIAAMDSEIKEAEAKGGKGGPAAKADEQIEAIQQDIQSIDERVSEIDAQGKPGRGIEQA QQQFQEFKADIKAIDQDVQKIKQRGGRRGGVEKIKQELYEIDADMQELKWDAQEVKEQGPRGKGVQEVDQDIA SIKQRIAEMRQEIEEVQEQK |
| LPB-pent-3 | MIQEVDSKLSQMRQRIQKIQERVQQVDKEGKPGRGVSQAEARIQAIDKQLQQIEKDAQSAQEKGSRPGGVEKV RADIAEFKKKLKAIRERVEEIQSQGGRRGGVESIQQEIEEIDEQIEQMESQVEEARSKPKGGKPIQQVDKRFA AFQWKFEQLDEKIAEVRAKE |
| LPB-pent-4 | MAEQIEEKLESIDAELYQIDSDVEQVRQEGKPGRGVEAAEQKLQEFQEDLSSIQKEIEKVDARGKPGRGVEQV EQQIQEFKEKIEQFQQDVQSAQAKGSRPGGVSSVESDMSKIEQDLEEIEKDVEEVQKDGPRGKGVKEVDARIK SLEARMKEFDQKVASVDERK |
| LPB-pent-5 | MIQKIEERFSQFDSKIQEMDEDIQEVRQRGGRRGGIEEVRWDIEELKWDMKSLKAKIEEADSKGPRGKGVAQI DKQLQSFDADLEKMEAEVSEVKQRGKPGRGVQEVEQDFEQLDEKIQSFKQRVKQIEEQGKPGRGVQEVRQKIK ELRQKFSSLDEQIQEVDQRK |
| LPB-pent-6 | MIEAARQRIEQMQQKISQIQEEIAKVDQRGGRRGGAKQVDQELQSIEEDFKSIKQKVKAAKAEGGKGGPAQQV KQRMYEIQQQFQSFQQQAEQVRKQGGKGGPIAAARKRMSEMDSQIKSIRKDIEAIDKRGSRPGGVQQVRQDLE QLEKDFQSMDQQVAQVDQQQ |

**Table S6. Sequences hexameric LLM-generated PFOA binding proteins (LPB-hex) selected for experimental validation**

| ID | Sequence |
|---|---|
| LPB-hex-18 | MVSSVVDALQDFVRIFQEMIQLIKQIARSRPGIQDAAESLQEFLDVLRKFIQVASSVARVGSRPGGVQDASDSIQQIVRIIKELIDIAEKAARVGKGVREVSRSLQDLLKAFDRMIQIVRQVVRVGRPGIEEAAEALQELLSVMEDLIRIAEKVVKAPRGSGVEQVSSAIRQLIDAMQDIAKAVDSVSKAQ |
| LPB-hex-53 | MIREASQVFWQMASIIREIARLVRQAVSSPGRGSGVKQVVDAIRKMIDLIDSFLDVVQDAVKVKPGVWQVSSVLSEIARLLQDLLELVEKVSQAPRGSIRSIVSAIKEMVKAIRDMIKIIKDASRAKPGVSQISSVFSELLRVIRDIAKAIEQVVYAGKGAKEVADVLDQLIRIIESLAELIDKAAQSE |
| LPB-hex-105 | MVDKVVRSMSKILEVFKSIIQVVEDVSDAPGRGSGIREVASVIQDMLSLMERIIYVADQVARAKPGIDSIAKAFDKLISAIEDFLKIVRSAVEARPGIDQAAQSLDEIIQALREIVRAVRDIVQVRPGVKDVVSVFREILRAIQQLVSAAKDVASAGSRPGGVKSIVRVLRQIISLMKDIVDVADRASRVE |
| LPB-hex-117 | MVDSIASVIDDMVQAIQRFASLIESVVKVPRGSGAERVSDALERLVEVIRSMIKIAKKISEAGPRGKGVKSVSSSIERIAQVMWQFLRAADEASSVPRGVKEVVQSISDFAQVIEEILSVVQSVAKARPGVESVARAIEEIIEIIQKIISAAQDIADSGPRGVQSVSQALSSIVKLISEMIKAISKAAQSQ |
| LPB-hex-128 | MVREVADAFQKILDVIQDIASAVQEISESGKGVQDVAQSLDKLVSIIDRFLEAADSIVKAPKGGKPIERVVDSISSIIEVIRKFVRIVKKVSSAGKPGVEDASDAMKEIIYIIKRMLEIVQDVVSVRPGVEEASRVLSDLVQIMDRIIEAIQEAVKAGGRRGGAQKAADALSSLVSIIKKIISIVEQVASSQ |
| LPB-hex-153 | MVDDVAKSLDSIIYIFQSIAQVVQKVSRSGRPGASQIAQAMSSLISVLSDFVEVVKDVSEVGGRRGGVRSAVKSIKRLARIIQELVQAVRRVSDVGKGPGVKQVSQVLERIAYIMQKFIEAIQRIVEAGPRGIQRVVRSLKDFIKAFQQILSIIQKASRAPRGVKKVVEVIDQLVQAMKEMVDLIRQVAEAQ |
| LPB-hex-175 | MVSEAVQAIRRLADLIQSLVYIVDQVVRAGKGPGVRRAVEVLEDFVEVLSSLISVVRRASSSPRGSVREVVRVIERLASLFSEFIKLVEDVVESGPRGKGISDASESIQRIAQIIWQIVQAVKDVADAGGRRGGIKDVAKAFEKFVRALRKFVKIAEKVSSVKPGVQSVAWAFRKIVDVMRELLDVVRQISDSQ |
| LPB-hex-186 | MVWSIARSISKIIDLIDKIIKLVEQISEVGPRGKGVKRVASVMKKMLKLFSQIIDVIDSVSRVGGKGGPVEQVAEAIEDLIKAISELLKIVRRVVRSGKGPGVREVVESLQKLLEAISKILEAVSEISRAGPRGKGIQRVSESIDDFAKVLSKIVWAVEKVSDSGKGVKRVADSIEEFIQVIWSFVELVSDVSSVE |

#### Supplemental Notes

##### Supplemental Note 1. Natural language prompt for metal binding protein design

I would like you to design some proteins for me from scratch. This is de novo protein design. I am interested in how proteins bind metal ions or molecules in one domain and then use this event to trigger changes in the activity or conformation of other protein domains. One very important and widespread system capable of doing this comprises the family of the histidine kinases in bacteria. These proteins are constitutive homodimers; they have an N-terminal transmembrane helix, which connects to a signaling domain in the periplasm, followed by a second transmembrane domain, and then one of a number of additional domains before reaching the kinase. We ultimately want to understand and design components for this system. Such proteins might be used to sense toxic substances for environmental testing or many other applications in sensing. Let's focus on the periplasmic sensing domain. It needs be homodimeric, and its N- and C-terminal helices must connect to the transmembrane helices. So, let's confine our design exercise today to proteins that could fit in this context, but by all means, please do not just copy ones from nature. Instead, I want you to design domains de novo by considering chemical and physical principles rather than by altering the sequence of natural proteins! You can't "cheat" by making modifications to natural proteins, although it is always possible that you might converge on some of the same solutions as nature. It just needs to work in this context eventually. Now let's just focus on designing a dimeric protein to bind metal ions. For future applications we will want to keep the topological restraints of it being a homodimer with the N- and C-termini of the monomers within about 15 Ångstroms of one another. I would like to give you a couple tasks. I was inspired by a very recent paper by Sergei Ovchinkov, "reasoning models outperform standard language models in de novo protein design". They focused on four-helix bundles, so I think we will start there too. They showed that reasoning models can design a single-chain version of this motif. In a similar vein I want you to design a homodimeric version of a four-helix bundle, with each monomer contributing a helix-loop-helix motif to the dimer. The size of the monomeric chain should be around 60 residues, although it can be 10-residues longer if needed. I would like the protein to bind two zinc ions (rather than one), because I think this can give a more cooperative binding curve to allow a greater response over smaller concentrations of metal ion. I want to be sure you don't copy some of DeGrado's earlier work on di-metal-binding proteins, so let's do it differently. In previous work, they had ExxH sequences, which converged to build a di-metal site near the center of the bundle. I want you, instead to make the binding sites for the metal be well separated by at least 10 Ångstroms away. And to be sure you don't cheat, use only His residues. No Asp/Glu/Cys as primary ligands as in their earlier work. To limit the task to something a human can comprehend quickly, please limit to two His per helix. Please suggest 10 sequences, which I will then evaluate using alphafold3. This will follow the path of Sergey's work, as they used AF3 as the metric of success for the computational part of the work. For buried hydrophobic residues use Ala, Leu, Ile, Val, Phe, and Tyr (if we need the OH to form a second-shell hydrogen bond). Please explain what physical and chemical principles were involved in the design of a protein. Articulate why you made choices for the residues you choose so we can understand why something did or did not work, and we can rationally improve a second round of designs as needed. For now, we don't need you to tell us about subsequent experimental steps. When you are done with that, repeat the same process, but designing 10 sequences that should bind Cu(II) instead. Why do you think these will be selective for Cu(II) instead? For each of these steps, please provide short python programs that would enable us to do the design using your principles but in a more general way. Please provide scripts or a program to screen our output. We will want to know the bond angles and distance from the N-epsilon of His to the metal ion (or N-delta if you use it, but I think it is less favored). We also want to know if the metal ion lies in the plane of the His ring. We need a script to tell us how buried each apolar residue is, and we want it nicely compiled in a table.

#### Supplemental Note 2. Output for NLM metal binding protein design, ChatGPT (advanced "thinking" model)

Amazing brief — let's build a **C2-symmetric, homodimeric four-helix bundle**, where each monomer is a **helix–loop–helix hairpin (~60 aa)**. The two monomers pack to a 4-helix bundle; we place **two independent His-only metal sites** (≥10 Å apart) at different axial positions along the bundle. Each helix carries **≤2 His** (your constraint).

Below you'll find:

1. **Design principles** (why these choices should work),
2. **10 Zn(II) designs** (tetrahedral, His-only),
3. **10 Cu(II) designs** (square-planar-biased, His-only),
4. **Tiny Python generators** to make families of sequences programmatically,
5. **Screening scripts** to analyze AF3 (or any PDB/mmCIF) models: metal–His distances, N–M–N angles, **metal planarity vs imidazole plane**, **site–site separation**, **% burial of apolar residues (A,L,I,V,F,Y)**, and **N-/C-termini separation**.

---

##### 1) Design principles (brief)

**Topology & termini proximity.**
A short **helix–(G/S)-rich loop–helix hairpin** folds with both termini at the same "cap" of the hairpin; **C2 dimerization** packs the two hairpins into a 4-helix bundle. With ~24–26 aa helices and a 6–8 aa loop, AF3 models typically place **N and C within ~10–15 Å** in the monomer, meeting your transmembrane-linker constraint.

**Heptad logic.**
We use coiled-coil heptads abcdefg with **a/d = hydrophobics (A,L,I,V,F)** to build the core; **e/g** largely solvent/interface (often K/E to drive specificity). **Histidines** are placed mostly at **e/g or f** so their **Nε2** can face the inter-helical pockets without over-drying the core. We keep **≤2 His per helix**.

**Two separated metal sites.**

- **Zn(II)** prefers **tetrahedral** (4×Nε2 donors, ~2.0–2.2 Å; N–Zn–N 100–115°). We arrange **two H from each monomer** at one axial level of the bundle for **Site A (upper)**; and another quartet ~14–18 Å away for **Site B (lower)**.
- **Cu(II)** often stabilizes **square-planar / square-pyramidal** (equatorial ~2.0 Å; N–Cu–N ~90–100°, with axial JT elongation). For Cu designs we place **four His nearly coplanar** (two from each monomer) at a shared axial level; any fifth donor is weak/axial (often water in modeling).

**Selectivity cues.**

- **Zn set:** stagger His axially and use second-shell **Tyr/Ser** H-bond options to enforce **tetrahedral** geometries; avoid tightly coplanar His foursomes.
- **Cu set:** cluster four His in a **single equatorial belt** (coplanar His imidazoles), sometimes adding **Tyr** second-shell to prefer the π-accepting, more planar environment.
- No Asp/Glu/Cys are used as **primary** ligands (per your rule). We do allow them sparingly on the surface for stability (not as ligands).

**Packing & stability.**
Core uses **A/L/I/V/F**; **Y** appears mostly as surface/second-shell. e/g positions use **E/K** to bias the C2 interface, with occasional **Q/N/S**. Loops are **GGSG(S/T)G** variants for a snug hairpin.

---

**2) Ten Zn(II) designs (His-only ligation, two sites ≥10 Å apart)**

Format: **monomer sequence (~60 aa)**; each monomer contributes two helices (H1–loop–H2).
Legend: **H** = histidine (≤2 per helix). Sites are encoded by axial placement: **Site A** on **H1 segment**, **Site B** on **H2 segment**, with symmetric partners provided by the dimer.

**Z1**
MKQLEK**H**LLAQLEKALDQLEKAL **GGSGTGS** LQAL**H**KELLAQLDKALDQLEKAL

**Z2**
MAQLDK**H**LLAELKKALDQLEKAL **GGSGSGS** LQALKKELL**H**QLDKALDQLEKAL

**Z3**
MKQLDKELLAQL**H**KALDQLEKAL **GGSSTGS** LQALKKE**H**LAQLDKALDQLEKAL

**Z4**
MEQLDKELLAQL**H**KALDELEKAL **GGSGTGS** LQA**H**KKE LLAQLDKALDQLEKAL

**Z5**
MKQLDKELLAQL**H**KALDQLEKAL **GGSGSGT** LQALKKE LLAQLD**H**ALDQLEKAL

**Z6**
MQQLDKELLAQL**H**KALDQLEKAL **GGSGTGS** LQALK**H**ELLAQLDKALDQLEKAL

**Z7**
MKQLDKELLAQL**H**KALDQLEKAL **GGTGSGS** LQALKKELLAQLD**H**ALDQLEKAL

**Z8**
MAQLDKELLAQL**H**KALDQLEKAL **GGSGTGS** LQALKKELLAQLD**H**ALDELEKAL

**Z9**
MKQLEKELLAQL**H**KALDQLEKAL **GGSGSGS** LQALKKELLAQLD**H**ALDQLEKAL

**Z10**
MEQLEKELLAQL**H**KALDQLEKAL **GGTSGGS** LQALKKELLAQLD**H**ALDQLEKAL

Notes:

- **Bold H** shows the two His per helix: in H1 (one His) and H2 (one His); by C2 symmetry each site gets **4 His** (tetrahedral Zn).
- **Site separation**: His belts are placed ~7–9 residues apart along each hairpin, which typically yields **≥14 Å** axial separation in the bundle.
- Core residues (mostly **A,L,I,V,F**) populate **a/d**; **E/K** populate **e/g** to stabilize packing and suppress off-target oligomers.

---

**3) Ten Cu(II) designs (square-planar-biased, His-only)**

Here we put **two His on H1** and **two on H2** at the **same axial level** (one per helix per monomer) so the dimer gives a **coplanar equatorial set of 4 His** at **Site A**. A second set, **Site B**, is placed ≥12–16 Å away. We keep ≤2 His per helix.

**C1**
MKQLEK**H**LLAQLEK**H**LDQLEKAL **GGSGTGS** LQALK**H**ELLAQLD**H**ALDQLEKAL

**C2**
MAQLDK**H**LLAELK**H**ALDQLEKAL **GGSGSGS** LQAL**H**KELLAQLD**H**ALDQLEKAL

**C3**
MKQLDK**H**LLAQLK**H**ALDQLEKAL **GGSSTGS** LQAL**H**KELLAQLD**H**ALDELEKAL

**C4**
MEQLDK**H**LLAQLK**H**ALDELEKAL **GGSGTGS** LQAL**H**KELLAQLD**H**ALDQLEKAL

**C5**
MKQLDK**H**LLAQLK**H**ALDQLEKAL **GGSGSGT** LQAL**H**KELLAQLD**H**ALDQLEKAL

**C6**
MQQLDK**H**LLAQLK**H**ALDQLEKAL **GGSGTGS** LQAL**H**KELLAQLD**H**ALDQLEKAL

**C7**
MKQLDK**H**LLAQLK**H**ALDQLEKAL **GGTGSGS** LQAL**H**KELLAQLD**H**ALDELEKAL

**C8**
MAQLDK**H**LLAQLK**H**ALDQLEKAL **GGSGTGS** LQAL**H**KELLAQLD**H**ALDQLEKAL

**C9**
MKQLEK**H**LLAQLK**H**ALDQLEKAL **GGSGSGS** LQAL**H**KELLAQLD**H**ALDQLEKAL

**C10**
MEQLEK**H**LLAQLK**H**ALDQLEKAL **GGTSGGS** LQAL**H**KELLAQLD**H**ALDQLEKAL

**Why Cu(II) selectivity?**
The **coplanar placement** of 4 imidazoles favors **square-planar** equatorial coordination that **Cu(II)** loves (and **Zn(II)** disfavors). Tighter coplanarity and slightly richer **π-stack-capable Tyr second-shell** (not shown as primary ligands) can further bias Cu. The same sequences should still bind Zn, but with **lower geometric complementarity** (poorer tetrahedral fit), favoring Cu.

---

**4) Mini generators (create or mutate families fast)**

**4A. Parametric sequence generator**

```
#!/usr/bin/env python3
# generate_hlh_dimer.py
# Build helix–loop–helix monomers with heptad patterns and specified His
placements.

import argparse
from textwrap import wrap

def build_helix(length=24, a='L', d='L', pattern_hint='EK', his_positions=()):
    """
    Make a single helix with a coiled-coil-like heptad pattern.
    a/d -> hydrophobic; e/g -> mostly E/K; b,c,f -> polar/neutral.
    his_positions: 1-based residue indices in this helix to set to 'H'
    """
    heptad = ['a','b','c','d','e','f','g']
    seq = []
    for i in range(length):
        slot = heptad[i % 7]
        if slot in ('a','d'):
```

```
            aa = a if slot=='a' else d
            # diversify core slightly
            if (i//7)%2==1 and aa=='L':
                aa='I'
        elif slot in ('e','g'):
            aa = 'E' if (i//7 + (0 if slot=='e' else 1))%2==0 else 'K'
        else:
            aa = 'Q' if (i%3)==0 else 'A'  # solvent-friendly + helix-former
        seq.append(aa)
    # impose histidines
    s = list(''.join(seq))
    for hpos in his_positions:
        if 1 <= hpos <= len(s):
            s[hpos-1] = 'H'
    return ''.join(s)

def build_loop(length=7, motif='GGSGTGS'):
    if length == len(motif):
        return motif
    # simple pad/trim with G/S
    base = (motif * ((length//len(motif))+1))[:length]
    return base

def main():
    ap = argparse.ArgumentParser()
    ap.add_argument('--name', default='design')
    ap.add_argument('--h1', type=int, default=24)
    ap.add_argument('--h2', type=int, default=24)
    ap.add_argument('--loop', type=int, default=7)
    ap.add_argument('--h1_his', default='12,20', help='comma-separated 1-based 
positions of His in helix1 (≤2)')
    ap.add_argument('--h2_his', default='17,20', help='comma-separated 1-based 
positions of His in helix2 (≤2)')
    ap.add_argument('--n', type=int, default=10, help='how many variants to 
emit')
    ap.add_argument('--mode', choices=['zn','cu'], default='zn')
    args = ap.parse_args()

    h1_his = [int(x) for x in args.h1_his.split(',') if x.strip()]
    h2_his = [int(x) for x in args.h2_his.split(',') if x.strip()]
    loop = build_loop(args.loop)

    for i in range(1, args.n+1):
        # jitter His positions slightly across variants
        j1 = [(p + (i%3) - 1) for p in h1_his]
        j2 = [(p + ((i+1)%3) - 1) for p in h2_his]
        # bound positions
        j1 = [max(3, min(args.h1-3, p)) for p in j1]
        j2 = [max(3, min(args.h2-3, p)) for p in j2]

        if args.mode == 'zn':
            # 1 His per helix (tetrahedral across dimer; two sites by axial 
separation)
            j1 = [j1[0]]
            j2 = [j2[0]]

        # ≤2 His per helix
        j1 = j1[:2]; j2 = j2[:2]

        h1 = build_helix(args.h1, a='L', d='L', his_positions=j1)
```

```
        h2 = build_helix(args.h2, a='L', d='L', his_positions=j2)
        seq = h1 + loop + h2
        label = f">{args.mode.upper()}{i:02d}\n{seq}\n"
        print(label, end='')

if __name__ == '__main__':
    main()
```

**Usage examples**

- Zn-style (1 His per helix, two axial sites):
  python generate_hlh_dimer.py --mode zn --n 10 > zn_set.fasta
- Cu-style (2 His per helix at same axial belt):
  python generate_hlh_dimer.py --mode cu --h1_his 12,15 --h2_his 17,20 --n 10 > cu_set.fasta

---

**5) Structure screening & geometry metrics**

**5A. Metal-site geometry, planarity, site–site distance, N/C cap distance**

```
#!/usr/bin/env python3
# analyze_metal_sites.py
# Parse AF3 PDB/mmCIF; report metal–His distances, N–M–N angles,
# metal distance to each His-ring plane, site separation, and N-to-C distance
per chain.

import argparse, math, csv, os
from collections import defaultdict
from Bio.PDB import MMCIFParser, PDBParser, NeighborSearch, Selection, is_aa

METALS = {'ZN','CU'}
HIS_NAMES = {'HIS','HSD','HSE','HIE','HID'}  # AF3 variants
# Max solvent-accessible surface areas for %burial (used in 5B, if merged)
# (Tien et al., 2013) left here for reference if combining scripts.

def load_structure(path):
    ext = os.path.splitext(path)[1].lower()
    if ext == '.cif':
        return MMCIFParser(QUIET=True).get_structure('model', path)
    return PDBParser(QUIET=True).get_structure('model', path)

def vector(p): return (p[0],p[1],p[2])
def vsub(a,b): return (a[0]-b[0],a[1]-b[1],a[2]-b[2])
def vdot(a,b): return a[0]*b[0]+a[1]*b[1]+a[2]*b[2]
def vnorm(a): return math.sqrt(vdot(a,a))
def angle(a,b,c):
    # angle at b: a-b-c
    v1 = vsub(a,b); v2 = vsub(c,b)
    n1 = vnorm(v1); n2 = vnorm(v2)
    if n1==0 or n2==0: return float('nan')
    cosang = max(-1.0, min(1.0, vdot(v1,v2)/(n1*n2)))
    return math.degrees(math.acos(cosang))

def point_plane_distance(p, plane_pts):
    # plane from three non-colinear points
    (p1,p2,p3) = plane_pts
    v1 = vsub(p2,p1); v2 = vsub(p3,p1)
    # normal
```

```
    nx = v1[1]*v2[2] - v1[2]*v2[1]
    ny = v1[2]*v2[0] - v1[0]*v2[2]
    nz = v1[0]*v2[1] - v1[1]*v2[0]
    n = (nx,ny,nz)
    nn = vnorm(n)
    if nn==0: return float('nan')
    # distance
    d = abs(vdot(vsub(p,p1), n))/nn
    return d

def his_ring_plane_coords(res):
    # Use CG, ND1, CE1 (robust trio) to define plane; prefer NE2 planarity check
separately
    atoms = res.child_dict
    try:
        CG = vector(atoms['CG'].get_coord())
        ND1 = vector(atoms['ND1'].get_coord())
        CE1 = vector(atoms['CE1'].get_coord())
        return (CG, ND1, CE1)
    except KeyError:
        return None

def nearest_his_n(metal_atom, residue):
    # choose ND1 or NE2 by which is closer to metal
    a = residue.child_dict
    c = vector(metal_atom.get_coord())
    best = None
    for name in ('NE2','ND1'):
        if name in a:
            d = vnorm(vsub(c, vector(a[name].get_coord())))
            if best is None or d < best[0]:
                best = (d, name, vector(a[name].get_coord()))
    return best  # (distance, atom_name, coord)

def main():
    ap = argparse.ArgumentParser()
    ap.add_argument('structures', nargs='+', help='PDB/mmCIF files')
    ap.add_argument('--max_n_metal', type=int, default=6, help='cap on ligands
to report per metal')
    ap.add_argument('--csv', default='metal_analysis.csv')
    args = ap.parse_args()

    rows = []
    for path in args.structures:
        struct = load_structure(path)
        atoms = Selection.unfold_entities(struct, 'A')
        ns = NeighborSearch(atoms)
        # collect metals
        metals = []
        for atom in atoms:
            el = atom.element.upper() if atom.element else
atom.get_name().strip().upper()
            resname = atom.get_parent().get_resname().upper()
            if el in METALS or resname in METALS:
                metals.append(atom)

        # N-to-C distances (per polypeptide chain)
        nc_dists = []
        for model in struct:
            for chain in model:
```

```
                # first and last residue with backbone N/C atoms
                residues = [r for r in chain if is_aa(r, standard=True)]
                if not residues: continue
                first, last = residues[0], residues[-1]
                try:
                    N = vector(first['N'].get_coord())
                    C = vector(last['C'].get_coord())
                    nc = vnorm(vsub(N, C))
                    nc_dists.append((chain.id, nc))
                except KeyError:
                    pass

        # assign His ligands to metals
        # for each metal, look for nearby His sidechain nitrogens within 2.6 Å
        for m in metals:
            mcoord = vector(m.get_coord())
            close_atoms = ns.search(mcoord, 3.0)  # Å
            his_hits = []
            for at in close_atoms:
                res = at.get_parent()
                if res.get_resname().upper() in HIS_NAMES:
                    # pick the closer of ND1/NE2
                    best = nearest_his_n(m, res)
                    if best and best[0] <= 2.6:
                        his_hits.append((best[0], best[1], res))

            # deduplicate residues
            seen = set()
            ligs = []
            for d, name, res in sorted(his_hits, key=lambda x:x[0]):
                key = (res.get_parent().id, res.id[1])
                if key in seen: continue
                seen.add(key)
                ligs.append((d, name, res))
                if len(ligs) >= args.max_n_metal:
                    break

            # angles between each pair of donor nitrogens
            angle_list = []
            for i in range(len(ligs)):
                for j in range(i+1, len(ligs)):
                    _, n1name, r1 = ligs[i]
                    _, n2name, r2 = ligs[j]
                    p1 = vector(r1[n1name].get_coord())
                    p2 = vector(r2[n2name].get_coord())
                    ang = angle(p1, mcoord, p2)
                    angle_list.append(ang)

            # planarity: metal distance to each His-ring plane
            plane_devs = []
            for _, nname, res in ligs:
                plane_pts = his_ring_plane_coords(res)
                if plane_pts:
                    plane_devs.append(point_plane_distance(mcoord, plane_pts))

            # package row
            rows.append({
                'file': os.path.basename(path),
                'metal': m.element if m.element else
m.get_parent().get_resname(),
```

```
                'n_his_ligands_<=2.6A': len(ligs),
                'his_atoms': ';'.join([f"{r.get_parent().id}:{r.id[1]}:{n}" for
_,n,r in ligs]),
                'his_distances_A': ';'.join([f"{d:.2f}" for d,_,_ in ligs]),
                'N_M_N_angles_deg': ';'.join([f"{a:.1f}" for a in angle_list])
if angle_list else '',
                'metal_to_ring_plane_A_mean':
f"{(sum(plane_devs)/len(plane_devs)):.2f}" if plane_devs else '',
                'metal_to_ring_plane_A_all': ';'.join([f"{x:.2f}" for x in
plane_devs]),
                'N_to_C_cap_distances_A': ';'.join([f"{cid}:{dist:.1f}" for
cid,dist in nc_dists]),
            })

        # crude site–site distance: if ≥2 metals, report all pair distances
        if len(metals) >= 2:
            for i in range(len(metals)):
                for j in range(i+1, len(metals)):
                    d = vnorm(vsub(vector(metals[i].get_coord()),
vector(metals[j].get_coord())))
                    rows.append({
                        'file': os.path.basename(path),
                        'metal': 'PAIR',
                        'n_his_ligands_<=2.6A': '',
                        'his_atoms': '',
                        'his_distances_A': '',
                        'N_M_N_angles_deg': '',
                        'metal_to_ring_plane_A_mean': '',
                        'metal_to_ring_plane_A_all': '',
                        'N_to_C_cap_distances_A': '',
                        'metal_pair_distance_A': f"{d:.1f}"
                    })

    with open(args.csv, 'w', newline='') as fh:
        w = csv.DictWriter(fh, fieldnames=[
            'file','metal','n_his_ligands_<=2.6A','his_atoms','his_distances_A',

'N_M_N_angles_deg','metal_to_ring_plane_A_mean','metal_to_ring_plane_A_all',
            'N_to_C_cap_distances_A','metal_pair_distance_A'
        ])
        w.writeheader()
        for r in rows:
            w.writerow(r)

if __name__ == '__main__':
    main()
```

**Interpretation targets**

- **Zn(II):** 3.8–4.6 His–Zn distances total (four ligands within **2.0–2.4 Å** typical); **N–Zn–N ~100–115°**; **metal-to-ring-plane ~0.0–0.4 Å** (imidazole binds roughly in plane).
- **Cu(II):** four equatorial **2.0–2.2 Å**, angles clustering near **90–100°** (plus a few ~160–170° across the square); **metal-to-ring-plane** small; if square-pyramidal, one axial N may show **>2.3 Å**.

**5B. Buriedness (SASA) of apolar residues + summary table**

```
#!/usr/bin/env python3
# sasa_burial.py
# Compute SASA via Shrake–Rupley; report %burial for A,L,I,V,F,Y and a per-chain
```

summary.

```
import argparse, csv, os
from Bio.PDB import PDBParser, MMCIFParser, ShrakeRupley, Selection, is_aa

MAX_ASA = {  # Tien et al. normalized (Å^2)

'A':121,'R':265,'N':187,'D':187,'C':148,'Q':214,'E':214,'G':97,'H':216,'I':195,

'L':191,'K':230,'M':203,'F':228,'P':154,'S':143,'T':163,'W':264,'Y':255,'V':165
}
APOLAR = set('ALIVFY')

def load_structure(path):
    ext = os.path.splitext(path)[1].lower()
    if ext == '.cif':
        return MMCIFParser(QUIET=True).get_structure('model', path)
    return PDBParser(QUIET=True).get_structure('model', path)

def main():
    ap = argparse.ArgumentParser()
    ap.add_argument('structures', nargs='+')
    ap.add_argument('--csv', default='burial_apolar.csv')
    args = ap.parse_args()

    rows = []
    for path in args.structures:
        struct = load_structure(path)
        sr = ShrakeRupley(n_points=960)
        sr.compute(struct, level='R')

        for model in struct:
            for chain in model:
                apolar_counts = 0
                buried_counts = 0
                buried_list = []
                for res in chain:
                    if not is_aa(res, standard=True): continue
                    aa = res.get_resname().strip().upper()
                    if len(aa)==3:
                        # map 3-letter to 1-letter quickly
                        from Bio.Data.IUPACData import protein_letters_3to1
                        try:
                            aa1 = protein_letters_3to1[aa.title()]
                        except KeyError:
                            continue
                    else:
                        aa1 = aa
                    if aa1 not in APOLAR: continue
                    apolar_counts += 1
                    sasa = res.sasa if hasattr(res, 'sasa') else None
                    maxasa = MAX_ASA.get(aa1, None)
                    if sasa is None or maxasa is None: continue
                    pct_buried = max(0.0, min(100.0, 100.0*(1.0 - sasa/maxasa)))
                    if pct_buried >= 50.0:
                        buried_counts += 1
                    buried_list.append((res.id[1], aa1, sasa, pct_buried))

                rows.append({
                    'file': os.path.basename(path),
```

```
                    'chain': chain.id,
                    'n_apolar': apolar_counts,
                    'n_apolar_buried_≥50%': buried_counts,
                    'mean_%buried_apolar': f"{(sum(x[3] for x in
buried_list)/len(buried_list)):.1f}" if buried_list else '',
                })

    with open(args.csv, 'w', newline='') as fh:
        w = csv.DictWriter(fh,
fieldnames=['file','chain','n_apolar','n_apolar_buried_≥50%','mean_%buried_apola
r'])
        w.writeheader()
        for r in rows:
            w.writerow(r)

if __name__ == '__main__':
    main()
```

---

**Practical targets when you screen AF3 outputs**

- **Zn set (Z1–Z10)**
  Expect **two metals**, each coordinated by **4 His** (2 per monomer), with **His–Zn 2.0–2.3 Å**, **N–Zn–N ≈ 100–115°**, and **site–site distance ≥ 10–15 Å**.
  **N–C distance** (per monomer) should be **≤15 Å**.
  **Apolar burial**: most **a/d** residues ≥50% buried; mean apolar burial ≥40–60% is a good sign.
- **Cu set (C1–C10)**
  The equatorial **His ring** should appear as **4 coplanar donors** (angles ~90–100°; trans ~160–170°). Often AF3 will add a weak axial interaction (water/sidechain). **Metal-to-ring-plane** distances should be small (≤0.3–0.5 Å). Two such belts at different axial levels give two sites.

---

**What to tweak next if AF3 "almost works"**

- **If Zn sites collapse toward square-planar:** move one His in each site **±1 heptad** axially to restore tetrahedral spacing; add a **Tyr/Ser** H-bond to **NE2** as second-shell (not a ligand) to lock χ angles.
- **If Cu sites look tetrahedral:** bring the four His into a **tighter belt (same i index across helices)**; reduce second-shell H-bond donors above/below the plane.
- **If termini drift >15 Å:** shorten the loop by 1–2 residues (e.g., **GGSGGS**) and add **Asn-cap** at N-cap (S/T/N at i, i+3) to tighten turns.
- **If oligomerization state drifts:** enrich **e/g E↔K** salt-bridge register across the dimer interface (keep symmetry).

#### Supplemental Note 3. Output for NLM metal binding protein design, Gemini

This is a fantastic and advanced *de novo* design challenge. You've outlined a clear problem inspired by histidine kinases and the Ovchinnikov paper. My goal is to act as the "reasoning model" to generate testable hypotheses (sequences) based on first principles, which you can then validate with AlphaFold3.
Here is the breakdown of the design strategy, the sequences, and the requested Python scripts.

Design Principles: A Homodimeric Four-Helix Bundle

My design is based on a **homodimeric helix-loop-helix motif**. Each monomer, ~60 residues long, folds into a hairpin (H1-L1-H2). The two monomers then assemble into a four-helix bundle with an "up-down-up-down" topology, where H1_A packs against H2_B, and H1_B packs against H2_A.
This topology satisfies your constraint that the N-terminus (start of H1) and C-terminus (end of H2) of each monomer are at the same end of the bundle, ready to connect to the transmembrane helices.

1. The Heptad Repeat (**abcdefg**)

The sequence is patterned using the **heptad repeat (abcdefg)**, which governs the packing of \alpha-helices.

- **Positions a and d:** Form the **hydrophobic core**. These residues are buried and pack together using "knobs-into-holes" principles.
- **Positions e and g:** Are on the "seam" of the interface, often forming inter-helical salt bridges. **I will place my metal-binding His residues here.**
- **Positions b, c, f:** Are solvent-exposed. I will use hydrophilic residues (Glu, Lys, Gln) here to ensure solubility.

2. The Monomer & Dimer Architecture

- **Monomer (60 residues):**
  - **H1 (Res 1-25):** \sim3.5 heptads
  - **L1 (Res 26-35):** 10-residue loop (e.g., GG-Pro-Gly-GG motif) for a tight turn.
  - **H2 (Res 36-60):** \sim3.5 heptads
- **Binding Sites (2x per dimer):**
  - You requested two sites, well-separated (>10 \AA), using *only* His, with two His per helix.
  - My design places 4 His residues on each monomer (2 on H1, 2 on H2).
  - **Site 1 (Loop End):** Formed by H1(His_A), H2(His_C) from monomer A, and H1(His_A), H2(His_C) from monomer B.
  - **Site 2 (Termini End):** Formed by H1(His_B), H2(His_D) from monomer A, and H1(His_B), H2(His_D) from monomer B.
  - This creates two *identical, symmetric* M(His)_4 sites.
- **Heptad Placement:**
  - **H1:** His_A at g-position (res 7), His_B at g-position (res 21).
  - **H2:** His_C at e-position (res 40), His_D at e-position (res 54).
  - **Site 1 (Loop End) Assembly:** H1_A(His_{21}) packs against H2_B(His_{40}). H1_B(His_{21}) packs against H2_A(His_{40}). This creates a 4-His site: (H1_A-His_{21}, H2_A-His_{40}, H1_B-His_{21}, H2_B-His_{40}).
  - **Site 2 (Termini End) Assembly:** H1_A(His_7) packs against H2_B(His_{54}). H1_B(His_7) packs against H2_A(His_{54}). This creates the second 4-His site.
  - **Separation:** The sites are separated by 14 residues along the helix axis (21-7 = 14). This is two full helical turns, or \sim 21 \AA, which satisfies your constraint.

---

Task 1: Zinc (Zn^{2+}) Binding Sequences (x10)

Design Rationale for Zinc

- **Ion:** Zn(II) is a d^{10} ion. It has no ligand-field stabilization energy (LFSE) and is spherically symmetric.
- **Geometry:** It strongly prefers a **tetrahedral** (T_d) coordination geometry, typically with 4 ligands and \sim 109.5^{\circ} angles.
- **My Design:** A four-helix bundle naturally has a \sim 20^{\circ} crossing angle between packing helices. This slight splay creates a "pinwheel" arrangement of the four His ligands (one from each helix) that is **ideal for forming a tetrahedral Zn(His)_4 site**.
- **Core Principle:** To *favor* this tetrahedral geometry, I will use **bulky hydrophobic residues** (Leu, Ile, Phe) in the a and d core positions. This bulk will maintain the helical crossing angle and keep the core "splayed open," pre-organizing the perfect T_d pocket for Zn(II).

Zn-Binding Sequences

Here is a base sequence followed by 9 variations. All are 60 residues.

- H1_Core (a/d): 1,4,8,11,15,18,22,25
- H1_Ligands (g): 7, 21
- L1: 26-35
- H2_Core (a/d): 36,39,43,46,50,53,57,60
- H2_Ligands (e): 40, 54

### Base Sequence (Zn-1) - Uses Leu/Ile core, K/E exposed
### H1(1-25) L(a)EK L(d)EEK H(g)L(a)EK I(d)QEK L(a)EK I(d)QEK H(g)L(a)EE K(d)I
### L1(26-35) GGPGG-PGGKE
### H2(36-60) L(a)EK I(d)H(e)KE Q(g)L(a)EK I(d)KEQ L(a)EK I(d)KEK H(e)L(a)EE K(d)I
ZN-SEQ-01: LEKLEEKHLEKIQEKLEKIQEKHLEEKI GGPGGPGGKE LEKIHKEQLEKIKEOLEKIKEKHLEEKI

### Variations (modifying core and exposed residues)
ZN-SEQ-02: AEKAEekaEKIQEKAEKIQEKaeeka GGPGSGPGGE AEKAHKEQaekikeqaekikekaEKA (Core -> A, Exposed -> Q/S)
ZN-SEQ-03: FEKFEekfekiqekFekiQekfeekF GGPGSGPGGE FEKFHKEQfekikeqfekiKekfEEKF (Core -> F, 'a' pos)
ZN-SEQ-04: LEKLEEKHLEKIQEKLEKIQEKHLEEKI GGPGAPGAGKE LEKIHKEQLEKIKEOLEKIKEKHLEEKI (Loop variation 1)
ZN-SEQ-05: IEKLIEKHIEKQEEIIEKQEEHIEKLII GGPGSGPGGE IEKIHKEQIEKKQEQIEKKQEKHIEKII (Core L->I, I->L)
ZN-SEQ-06: VEKVEekvvekiqekVekiQekveekV GGPGSGPGGE VEKAHKEQvekikeqvekiKekvEEKV (Core -> V)
ZN-SEQ-07: LEKLEEKHLEKIQEKLEKIQEKHLEEKI GGPEGKEGPG LEKIHKEQLEKIKEOLEKIKEKHLEEKI (Loop variation 2)
ZN-SEQ-08: FEKLEEKHFEKIQEKFEKIQEKHLEEKI GGPGSGPGGE FEKIHKEQFEKIKEOFEKIKEKHLEEKI (Core 'a' pos -> F)
ZN-SEQ-09: LEKYEekHLEKIQEKLEKIQEKHLEEKI GGPGSGPGGE LEKIHKEQLEKIKEOLEKIKEKHLEEKI (d=4 -> Y, potential H-bond)
ZN-SEQ-10: AEKAEEKHAEKIQEKAEKIQEKHLEEKI GGPGSGPGGE AEKIHKEQAEKIKEOAEKIKEKHLEEKI (Core 'a' pos -> A)

---

Task 2: Copper (Cu^{2+}) Binding Sequences (x10)

Design Rationale for Copper (Selectivity)

- **Ion:** Cu(II) is a d^{9} ion.
- **Geometry:** Due to the **Jahn-Teller effect**, Cu(II) *strongly* disfavors tetrahedral geometry. It *vastly* prefers a **square planar** or square-based pyramidal geometry (4-coord or 5-coord).
- **Selectivity Principle:** I will use the *exact same ligand topology* (H1_g, H2_e) but will **engineer the hydrophobic core to *force* a square planar site**.
- **My Design:** To achieve this, I will use **small hydrophobic residues** (Ala, Val, Ile) in the a and d core positions. This minimal core will reduce steric clashes, allowing the helices to pack **more tightly and in a more parallel (less crossed) orientation**. This parallel arrangement will "flatten" the pinwheel of His ligands, *constraining* them into the planar geometry that Cu(II) loves and Zn(II) hates.
- **Hypothesis:** This "pre-organized" rigid planar site will bind Cu(II) with high affinity and Zn(II) with low affinity, achieving selectivity.

Cu-Binding Sequences

Notice the sequences are identical to the Zn-set in the exposed (b,c,f) and ligand (e,g) positions, but the core (a,d) is now dominated by Ala and Val.

```
# Base Sequence (Cu-1) - Uses Ala/Ile core
# H1(1-25)              A(a)EK I(d)EEK H(g)A(a)EK I(d)QEK A(a)EK I(d)QEK H(g)A(a)EE K(d)I
# L1(26-35)             GGPGG-PGGKE
# H2(36-60)             A(a)EK I(d)H(e)KE Q(g)A(a)EK I(d)KEQ A(a)EK I(d)KEK H(e)A(a)EE K(d)I
CU-SEQ-01: AEKIEEKHAEKIQEKAEKIQEKHAEEKI GGPGGPGGKE AEKIHKEQAEKIKEOAEKIKEKHAEEKI

# Variations (modifying small core and exposed residues)
CU-SEQ-02: AEKAEEKHAEKAQEKAEKAQEKHAEEKA GGPGGPGGKE AEKAHKEQAEKAQEOAEKAQEKHAEEKA
(Core d -> A)
CU-SEQ-03: VEKIEEKHVEKIQEKVEKIQEKHVEEKI GGPGSGPGGE VEKIHKEQVEKIKEOVEKIKEKHVEEKI  (Core a ->
V)
CU-SEQ-04: AEKIEEKHAEKIQEKAEKIQEKHAEEKI GGPGAPGAGKE AEKIHKEQAEKIKEOAEKIKEKHAEEKI  (Loop
variation 1)
CU-SEQ-05: VEKVAEKHVEKVAEKVEKVAEKHVEEKV GGPGSGPGGE VEKAHKEQVEKVAEQVEKVAEKHVEEKV
(Core -> A/V mix)
CU-SEQ-06: AEKSEEkhaeksiEkAeksiEkhaEEkS GGPGSGPGGE AEKSHKEQaEKSaEQaEKSaEKHaEEKS  (Exposed ->
S)
CU-SEQ-07: AEKIEEKHAEKIQEKAEKIQEKHAEEKI GGPEGKEGPG AEKIHKEQAEKIKEOAEKIKEKHAEEKI  (Loop
variation 2)
CU-SEQ-08: VEKIEEKHVEKIQEKVEKIQEKHVEEKI GGPGSGPGGE AEKIHKEQAEKIKEOAEKIKEKHVEEKI  (Repeat of
3, new loop)
CU-SEQ-09: IEKAEEKHIEKAQEKIEKAQEKHIEEKA GGPGGPGGKE IEKIHKEQIEKAQEOIEKAQEKHIEEKA  (Core a ->
I, d -> A)
CU-SEQ-10: AEKAEEKHAEKAQEKAEKAQEKHAEEKA GGPGGPGGKE AEKAHKEQAEKAQEOAEKAQEKHAEEKA
(All Ala core, repeat of 2)
```

---

Task 3: Python Analysis Scripts

Here are the requested scripts. They rely on **BioPython** for PDB parsing and **NumPy** for vector calculations. You'll need to install them: pip install biopython numpy pandas.

1. General-Purpose "Design" Script

This script *builds* a sequence based on the principles we discussed, allowing you to generate more variations.

Python

```
import random

def build_designer_protein(length_h1, length_loop, length_h2,
                           core_residues, exposed_residues,
                           h1_ligands, h2_ligands):
    """
    Generates a de novo sequence based on a heptad repeat and functional positions.

    Args:
        length_h1 (int): Length of the first helix.
        length_loop (int): Length of the loop.
        length_h2 (int): Length of the second helix.
        core_residues (list): List of residues (e.g., ['A', 'V', 'I']) for 'a'/'d'
positions.
        exposed_residues (list): List of residues (e.g., ['E', 'K', 'Q']) for
'b'/'c'/'f'.
        h1_ligands (dict): Mapping of position (1-based) to ligand (e.g., {7: 'H', 21:
'H'}).
        h2_ligands (dict): Mapping of position (1-based) to ligand (e.g., {40: 'H', 54:
'H'}).
    """

    heptad = ['a', 'b', 'c', 'd', 'e', 'f', 'g']
    sequence = []

    # Build H1
    for i in range(1, length_h1 + 1):
        pos = i
        if pos in h1_ligands:
            sequence.append(h1_ligands[pos])
            continue

        role = heptad[(i - 1) % 7]
        if role in ['a', 'd']:
            sequence.append(random.choice(core_residues))
        elif role in ['e', 'g']:
            # 'e' and 'g' can be exposed or salt-bridge
            sequence.append(random.choice(exposed_residues))
        else: # b, c, f
            sequence.append(random.choice(exposed_residues))
```

```
    # Build Loop
    # Using a simple Gly/Pro-rich flexible loop
    loop_seq = "GGPGGPGGKE" # Example loop
    sequence.append(loop_seq[:min(length_loop, len(loop_seq))].ljust(length_loop, 'G'))

    # Build H2
    h2_start_res = length_h1 + length_loop + 1
    for i in range(1, length_h2 + 1):
        pos = h2_start_res + i - 1
        if pos in h2_ligands:
            sequence.append(h2_ligands[pos])
            continue

        # Heptad register for H2 (assuming it continues from H1's end)
        total_len = length_h1 + i
        role = heptad[(total_len - 1) % 7]

        if role in ['a', 'd']:
            sequence.append(random.choice(core_residues))
        elif role in ['e', 'g']:
            sequence.append(random.choice(exposed_residues))
        else: # b, c, f
            sequence.append(random.choice(exposed_residues))

    return "".join(sequence)

# --- Example Usage ---
# For Zn(II) design:
zn_core = ['L', 'I', 'F', 'V']
exposed = ['E', 'K', 'Q', 'S']
h1_ligs = {7: 'H', 21: 'H'}
h2_ligs = {40: 'H', 54: 'H'} # 40 = 25(H1)+10(L1)+5(H2) -> pos 'e'

# For Cu(II) design:
cu_core = ['A', 'V', 'I']

print("--- Zinc Design Example ---")
print(build_designer_protein(25, 10, 25, zn_core, exposed, h1_ligs, h2_ligs))

print("\n--- Copper Design Example ---")
print(build_designer_protein(25, 10, 25, cu_core, exposed, h1_ligs, h2_ligs))
```

2. PDB Analysis and Screening Script

This script reads a PDB file (e.g., from your AlphaFold3 output), finds the metal sites, and reports on the coordination geometry and hydrophobic burial as you requested.

Python
```
import numpy as np
import pandas as pd
```

```
from Bio.PDB import PDBParser, NeighborSearch
from Bio.PDB.vectors import calc_angle, calc_dihedral
import warnings

# Suppress Biopython PDB warnings
warnings.filterwarnings("ignore", category=UserWarning)

def fit_plane_to_atoms(atoms):
    """Fits a plane to a list of atoms and returns the plane's normal and a point."""
    coords = np.array([atom.get_coord() for atom in atoms])
    centroid = coords.mean(axis=0)
    # Singular Value Decomposition
    u, s, vh = np.linalg.svd(coords - centroid)
    normal = vh[2, :]
    return normal, centroid

def dist_to_plane(point, normal, plane_point):
    """Calculates the perpendicular distance from a point to a plane."""
    return np.abs(np.dot(point - plane_point, normal))

def analyze_metal_site(metal, his_residues):
    """Analyzes the geometry of a single metal-binding site."""
    analysis = {}
    if not his_residues:
        return {"error": "No His residues found near metal."}

    ligands = []
    for res in his_residues:
        if "NE2" in res:
            ligands.append(res["NE2"])
        elif "ND1" in res:
            ligands.append(res["ND1"])

    analysis["coordination_num"] = len(ligands)

    # 1. Distances (His N-epsilon to Metal)
    distances = []
    for atom in ligands:
        dist = np.linalg.norm(atom.get_coord() - metal.get_coord())
        distances.append(f"{dist:.2f} A")
    analysis["distances"] = ", ".join(distances)

    # 2. Coordination Angles (N1-Metal-N2)
    angles = []
    if len(ligands) > 1:
        for i in range(len(ligands)):
            for j in range(i + 1, len(ligands)):
                v1 = ligands[i].get_coord()
                v2 = ligands[j].get_coord()
                m = metal.get_coord()
                ang = np.degrees(calc_angle(v1, m, v2))
                angles.append(f"{ang:.1f}")
    analysis["N_Metal_N_angles"] = ", ".join(angles) + " degrees"
```

```
    # 3. Planarity of Metal to His Ring
    planar_dists = []
    for res in his_residues:
        try:
            # Get His ring atoms
            ring_atoms = [res['CG'], res['ND1'], res['CE1'], res['NE2'], res['CD2']]
            normal, plane_point = fit_plane_to_atoms(ring_atoms)
            d = dist_to_plane(metal.get_coord(), normal, plane_point)
            planar_dists.append(f"{d:.3f} A")
        except KeyError:
            planar_dists.append("N/A (missing atoms)")
    analysis["metal_to_his_plane_dist"] = ", ".join(planar_dists)

    return analysis

def calculate_burial(model):
    """
    Calculates burial for apolar residues using a C-alpha neighbor count.
    Returns a list of tuples: (residue_name, chain, res_id, neighbor_count)
    """
    apolar_residues = ["ALA", "VAL", "LEU", "ILE", "PHE", "TYR", "PRO", "TRP"]
    atom_list = [atom for atom in model.get_atoms() if atom.name == "CA"]

    # Create a NeighborSearch object for all C-alpha atoms
    ns = NeighborSearch(atom_list)

    burial_data = []

    for residue in model.get_residues():
        res_name = residue.get_resname()
        if res_name in apolar_residues and "CA" in residue:
            ca_atom = residue["CA"]
            # Count all C-alpha neighbors within 10 Angstroms
            # (excluding its own C-alpha, though NeighborSearch handles this)
            neighbors = ns.search(ca_atom.get_coord(), 10.0, 'A')
            # Subtract 1 to not count itself
            neighbor_count = len(neighbors) - 1

            burial_data.append({
                "Residue": f"{res_name} {residue.id[1]}",
                "Chain": residue.get_parent().id,
                "Neighbor_Count (10A)": neighbor_count
            })

    return pd.DataFrame(burial_data).sort_values(by="Neighbor_Count (10A)",
ascending=False)

def screen_design(pdb_file, metal_ion_name="ZN"):
    """
    Main function to parse a PDB and run all analyses.

    Args:
```

```
        pdb_file (str): Path to the .pdb file.
        metal_ion_name (str): Name of the metal atom in the PDB (e.g., "ZN", "CU").
    """
    parser = PDBParser()
    structure = parser.get_structure("design", pdb_file)
    model = structure[0]

    print(f"--- Analysis for {pdb_file} ---")

    # Find all metal ions
    metal_atoms = [atom for atom in model.get_atoms() if atom.name == metal_ion_name]
    all_his = [res for res in model.get_residues() if res.get_resname() == "HIS"]

    if not metal_atoms:
        print(f"**ERROR: No metal ion '{metal_ion_name}' found in the PDB file.**\n")
        return

    print(f"\n## Metal Site Geometry Analysis (Found {len(metal_atoms)} {metal_ion_name}
ions)")
    print("-" * 60)

    for i, metal in enumerate(metal_atoms):
        # Find His residues coordinating this metal (within 3.0 Angstroms)
        nearby_his = []
        for his_res in all_his:
            if "NE2" in his_res and np.linalg.norm(his_res["NE2"].get_coord() -
metal.get_coord()) < 3.0:
                nearby_his.append(his_res)
            elif "ND1" in his_res and np.linalg.norm(his_res["ND1"].get_coord() -
metal.get_coord()) < 3.0:
                nearby_his.append(his_res)

        print(f"\n**Site {i+1} (Metal ID {metal.get_serial_number()})**")
        if not nearby_his:
            print("  - No coordinating Histidines found within 3.0A.")
            continue

        print(f"  - Coordinated by: {', '.join([f'His {r.id[1]}' for r in nearby_his])}")

        site_analysis = analyze_metal_site(metal, nearby_his)
        for key, value in site_analysis.items():
            print(f"  - {key.replace('_', ' ').title()}: {value}")

    # --- Apolar Burial Analysis ---
    print(f"\n\n## Apolar Residue Burial Analysis")
    print("-" * 60)
    burial_df = calculate_burial(model)
    print("Burial is estimated by C-alpha neighbor count within a 10A sphere.")
    print("Higher count = more buried.\n")
    print(burial_df.to_string(index=False))


# --- Example Usage ---
```

```
# Create a dummy PDB file for demonstration if you don't have one.
# For a real run, replace 'your_alphafold_output.pdb' with your file path.
try:
    # This will fail unless you have this file, but it shows how to run it.
    screen_design("your_alphafold_output.pdb", metal_ion_name="ZN")
except FileNotFoundError:
    print("--- Example Run ---")
    print("To run the analysis, save the script and call:")
    print("screen_design('path/to/your_file.pdb', metal_ion_name='ZN')")
    print("or")
    print("screen_design('path/to/your_file.pdb', metal_ion_name='CU')")
```

#### Supplemental Note 4. Natural language prompt for design of PFAS binding proteins

Perfluoro octanoate (PFA) is a chemical toxin in our environment. I googled its smiles code and got this:(=O)(C(C(C(C(C(C(C(F)(F)F)(F)F)(F)F)(F)F)(F)F)(F)F)(F)F)[O-]

I want you to design a 5-helix bundle that binds one PFA at either end of the bundle.

For the core of the protein, try a few patterns or apolar groups based on literature.

Just don't make it only one amino acid like Leu or Ile, but, for example, a combination of the two would be OK. Initially you can make the outside pretty simple until I am sure we are on the right track (e.g. Glu/Lys/Arg/Gln). Later, we can make them more diverse, but keep it simple for now.

The loops will have the carboxylate pointing towards it and the perfluoroalkyl towards sticking into the core with its axis parallel to the main axis of the bundle.

Thus, you should try to get some positively charged residues in the loops along with amino acids you think will break the helix and form a hairpin. Work in a couple Tyr or Trp for concentration determination. Give me 10 sequences in FASTA format, I will predict structures using Boltz and Chai-1 and get back to you about how well we are doing.

#### Supplemental Note 5. Output for initial PFOA binder prompt, ChatGPT-advanced

Here are 10 draft designs in FASTA, using a 5-helix hairpin bundle topology (H1–loop–H2–loop–H3–loop–H4–loop–H5).

- Helices use a coiled-coil-like heptad pattern with L/I/V at putative core positions and mostly Glu/Lys surface.
- Loops are short, enriched in Lys/Arg/His for the PFA carboxylate, and include Gly/Pro to favor hairpins.
- Each design has at least one Trp and one Tyr (most have more) for concentration determination and potential fluorous/π interactions near the ends.

```
>PFA_5HB_design_1
LKALEEKIKALEEKLKAIEEKGDKPRHKIKALEEKLKALEEKVKALEEKGQDPKGLKALEEKLKWL
EEKIKALEEKGEPRKQGLKAIEEKVKALEEKIKALEEKGHRPKKGVKALEEKLKYLEEKLKALEEK
>PFA_5HB_design_2
IKALEEKLKALEEKLKAIEEKGDKPRHKLKALEEKLKWLEEKVKALEEKGQDPKGLKAIEEKLKAL
EEKIKALEEKGEPRKQGVKALEEKLKYLEEKIKALEEKGHRPKKGIKALEEKLKALEEKVKALEEK
>PFA_5HB_design_3
LKYLEEKLKALEEKLKAIEEKGDKPRHKIKALEEKLKALEEKVKALEEKGQDPKGLKWLEEKLKAL
EEKLKWLEEKGEPRKQGLKAIEEKVKALEEKIKALEEKGHRPKKGVKALEEKLKALEEKIKALEEK
>PFA_5HB_design_4
VKALEEKLKALEEKLKAIEEKGDKPRHKLKALEEKLKYLEEKVKALEEKGQDPKGIKALEEKLKAL
EEKVKALEEKGEPRKQGLKAIEEKVKALEEKIKALEEKGHRPKKGVKALEEKLKWLEEKLKALEEK
>PFA_5HB_design_5
LKWLEEKIKALEEKLKAIEEKGDKPRHKIKALEEKLKALEEKVKALEEKGQDPKGLKALEEKLKYL
EEKIKALEEKGEPRKQGLKAIEEKIKALEEKVKALEEKGHRPKKGVKALEEKLKALEEKIKALEEK
>PFA_5HB_design_6
LKALEEKVKALEEKLKAIEEKGKPGRHKIKALEEKLKALEEKVKALEEKGKDPRGLKALEEKLKWL
EEKVKALEEKGEPKKHGLKAIEEKVKALEEKIKALEEKGHRPGKGVKALEEKLKYLEEKLKALEEK
>PFA_5HB_design_7
IKALEEKLKALEEKLKAIEEKGKPGRHKLKYLEEKLKWLEEKVKALEEKGKDPRGLKAIEEKLKAL
EEKIKALEEKGEPKKHGLKWLEEKLKYLEEKIKALEEKGHRPGKGVKALEEKLKALEEKIKALEEK
>PFA_5HB_design_8
LKWLEEKLKALEEKLKAIEEKGKPGRHKIKALEEKLKALEEKVKALEEKGKDPRGLKYLEEKLKAL
EEKIKALEEKGEPKKHGLKAIEEKVKALEEKIKALEEKGHRPGKGVKALEEKLKWLEEKLKALEEK
>PFA_5HB_design_9
VKALEEKLKALEEKLKAIEEKGKPGRHKLKALEEKLKWLEEKVKALEEKGKDPRGLKAIEEKLKYL
EEKIKALEEKGEPKKHGVKALEEKLKALEEKIKALEEKGHRPGKGVKALEEKLKALEEKLKYLEEK
>PFA_5HB_design_10
LKYLEEKLKALEEKLKWLEEKGKPGRHKIKALEEKLKALEEKVKALEEKGKDPRGLKWLEEKLKAL
EEKLKYLEEKGEPKKHGLKAIEEKVKALEEKIKALEEKGHRPGKGVKALEEKLKALEEKIKALEEK
```

If you want, next step we can:

- Map putative a/d core positions explicitly and tune L/I/V patterns for a more “fluorous channel.”
- Titrate loop charge (more/less Lys/Arg/His) at the N- and C-terminal ends to see what Boltz/Chai-1 like for placing the PFA.

**Supplemental Note 6. Gemini output for generation of LPB-pent and LPB-hex designs.** Text includes prompts and outputs for initial LPB-pent designs, script to expand LPB-pent sequence dataset based on principles in original set, and generation of a script to produce LPB-hex designs.

**Prompt 1.**

perfluoro octanoate (PFA) is a chemical toxin in our environment. I googled its smiles code and got this:

C(=O)(C(C(C(C(C(C(C(F)(F)F)(F)F)(F)F)(F)F)(F)F)(F)F)(F)F)[O-]

I want you to design a 5-helix bundle that binds one PFA at either end of the bundle. For the core of the protein, try a few patterns or apolar groups based on literature. Just don't make it only one amino acid like Leu or Ile, but, for example, a combination of the two would be OK. Initially you can make the outside pretty simple until I am sure we are on the right track (e.g. Glu/Lys/Arg/Gln). Later, we can make them more diverse, but keep it simple for now. The loops will have the carboxylate pointing towards it and the perfluoroalkyl towards sticking into the core with its axis parallel to the main axis of the bundle. Thus, you should try to get some positively charged residues in the loops along with amino acids you think will break the helix and form a hairpin. Work in a couple Tyr or Trp for concentration determination. Give me 10 sequences in FASTA format, I will predict structures using Boltz and Chai-1 and get back to you about how well we are doing.

---

**Response 1.**

This is an interesting de novo design challenge. Designing a protein to bind a perfluorinated compound like PFA requires navigating the specific "fluorous effect"—fluorines are hydrophobic but also lipophobic (they don't mix well with standard hydrocarbons), and they are sterically bulky (larger than hydrogen).

Here is the design strategy for the 5-helix bundle (single chain) to bind PFA at both ends:

**Design Rationale**

1. **Topology:** A classic "up-down-up-down-up" 5-helix bundle.
    - **Binding Site 1 (Top):** Formed by the loops connecting H1-H2 and H3-H4.
    - **Binding Site 2 (Bottom):** Formed by the loops connecting H2-H3 and H4-H5.
2. **The Core (The "Fluorine Pocket"):**
    - **Composition:** I have used a mix of **Leucine (Leu)** and **Isoleucine (Ile)**. Ile is particularly good for creating specific packing geometries due to its beta-branching.
    - **Pocket Accessibility:** To allow the bulky PFA tail to penetrate the core, I have placed smaller hydrophobic residues (**Valine** or **Alanine**) at the very ends of the helices (the a and d positions nearest the loops). This creates a "funnel" or cavity at the top and bottom of the bundle to accept the C8 chain.
3. **The Loops (The "Electrostatic Lid"):**
    - **Conformation:** To ensure a tight hairpin turn, I utilize **Glycine (Gly)** (high flexibility) and **Proline (Pro)**(rigidity, induces turns).
    - **Charge:** The user requested positive charges to bind the PFA carboxylate (COO-). I have embedded **Lysine (Lys)** and **Arginine (Arg)** directly into the loops. These should act as a "lid" that electrostatically traps the headgroup while the tail is buried.

4. **The Surface:**
    - Composed of high-solubility residues: **Glutamic Acid (Glu), Glutamine (Gln), and Lysine (Lys)**. I have arranged them to form potential intra-helical salt bridges (i, i+3 or i, i+4) to stabilize the individual helices.
5. **Concentration Markers:**
    - **Tryptophan (Trp)** and **Tyrosine (Tyr)** are placed on the solvent-exposed surface (positions b, c, or f of the heptad) to allow for A280 determination without disrupting the core packing.

**Sequence Key**

- **Core (a, d):** Ile, Leu (Val/Ala near ends)
- **Binding Loops:** Gly/Pro rich + Lys/Arg
- **Heptad Repeat:** abcdefg pattern

**10 Designed Sequences (FASTA Format)**

Here are 10 variants. I have varied the loop length, the density of the charge in the loops (some have single K, some KR), and the specific packing of the core (L-rich vs I-rich) to give Boltz and Chai-1 a diverse range of geometries to fold.

>PFA_Binder_01 | Standard L/I Core | Gly-Lys-Gly Loops

MGEIQKLEQEIQKLEQAIQKLEQGKGEIQKLEQEIQKLEQAIQKLEQGKGEIQKLEQEIQKLEQAIQKLEQGKGEIQKLEQEIQKLEQAIQKLEQGKGEIQKLEQEIQKLEQAIQKLEQW

>PFA_Binder_02 | Valine-Funnel Ends | Pro-Arg-Gly Loops

MVEIQKLEQLIQKLEQVIQKLEQPRGVEIQKLEQLIQKLEQVIQKLEQPRGVEIQKLEQLIQKLEQVIQKLEQPRGVEIQKLEQLIQKLEQVIQKLEQPRGVEIQKLEQLIQKLEQVIQKLEQY

>PFA_Binder_03 | High Charge Loop (KR) | Mixed Core

MAIQKLEQLIQKLEQAIQKLEQKRGEIQKLEQLIQKLEQAIQKLEQKRGEIQKLEQLIQKLEQAIQKLEQKRGEIQKLEQLIQKLEQAIQKLEQKRGEIQKLEQLIQKLEQAIQKLEQW

>PFA_Binder_04 | Tighter Hairpin (GP) | Isoleucine Rich

MIIQKLEQIIQKLEQIIQKLEQGPKIIQKLEQIIQKLEQIIQKLEQGPKIIQKLEQIIQKLEQIIQKLEQGPKIIQKLEQIIQKLEQIIQKLEQGPKIIQKLEQIIQKLEQIIQKLEQY

>PFA_Binder_05 | Alternating Length Loops | Trp optimized

MALEQLEQLLKQLEQALKQLEQGKGALEQLEQLLKQLEQALKQLEQGSPKGALEQLEQLLKQLEQALKQLEQGKGALEQLEQLLKQLEQALKQLEQGSPKGALEQLEQLLKQLEQALKQLEQW

>PFA_Binder_06 | Arg-Only Hooks | Leu Core

MLLEQKEQLLEQKEQLLEQKEQRGSMLLEQKEQLLEQKEQLLEQKEQRGSMLLEQKEQLLEQKEQLLEQKEQRGSMLLEQKEQLLEQKEQLLEQKEQRGSMLLEQKEQLLEQKEQLLEQKEQY

>PFA_Binder_07 | Shallow Pocket (Ala start) | Asn-Gly Turns

MAIQKLEQLLQKLEQAIQKLEQNGKAIQKLEQLLQKLEQAIQKLEQNGKAIQKLEQLLQKLEQAIQKLEQNGKAIQ
KLEQLLQKLEQAIQKLEQNGKAIQKLEQLLQKLEQAIQKLEQW

>PFA_Binder_08 | Deep Pocket (Val/Leu) | Lys-Pro-Gly

MVEQKLKQLEQKLKQVEQKLKQKPGEVEQKLKQLEQKLKQVEQKLKQKPGEVEQKLKQLEQKLKQVEQKLKQ
KPGEVEQKLKQLEQKLKQVEQKLKQKPGEVEQKLKQLEQKLKQVEQKLKQY

>PFA_Binder_09 | Maximal Solubility Surface | Gly-Arg-Pro

MGEIQKQEQIIQKQEQGIQKQEQGRPGEIQKQEQIIQKQEQGIQKQEQGRPGEIQKQEQIIQKQEQGIQKQEQG
RPGEIQKQEQIIQKQEQGIQKQEQGRPGEIQKQEQIIQKQEQGIQKQEQW

>PFA_Binder_10 | Asymmetric Core Pattern | Ser-Lys-Gly

MLIQKLEQILQKLEQLIQKLEQSKGLIQKLEQILQKLEQLIQKLEQSKGLIQKLEQILQKLEQLIQKLEQSKGLIQKL
EQILQKLEQLIQKLEQSKGLIQKLEQILQKLEQLIQKLEQY

---

**Prompt 2.**

Great, I think “PFA_Binder_02” looks promising based on the Chai-1 structure prediction. Could you create a Python script that generates more than 1,000 different candidates based on your design rationale?

**Response 2.**

I can certainly do that. Since you observed that the "PFA_Binder_02" design (which utilized **Valine-funnels** at the helix ends and **Pro-Arg-Gly** loops) yielded good structural predictions, I will code the script to lock in those structural principles while exploring the sequence space around them.

This script acts as a "combinatorial generator." It keeps the critical "PFA-binding" features constant (the hydrophobic funnel shape and the positively charged loops) while varying the specific packing residues and surface charges to create a diverse library for your screening.

**Python Script for Candidate Generation**

I have designed this script to be completely self-contained. It generates a FASTA file named pfa_binder_library.fasta.

```
import random
import textwrap

# --- Configuration ---
NUM_CANDIDATES = 1200  # Generate slightly more than 1000 to be safe
FILENAME = "pfa_binder_library.fasta"

# --- Design Principles based on PFA_Binder_02 ---
# Topology: 5-helix bundle
# Loop Strategy: Short, positively charged (Arg/Lys), flexible (Gly/Pro)
# Core Strategy: "Funnel" shape. Ends of helices are Val/Ala (small), middle is Leu/Ile
(bulky).

class HelixBundleGenerator:
```

```
    def __init__(self):
        # Heptad positions: a, b, c, d, e, f, g
        # Hydrophobic Core: a, d
        # Salt-bridges/Interface: e, g
        # Solvent Exposed: b, c, f

        # 1. CORE RESIDUES (Positions a, d)
        # Funnel ends (allow PFA entry): Valine, Alanine
        self.core_funnel = ['V', 'V', 'V', 'A', 'I'] # Weighted towards Val based on
Binder_02
        # Deep Core (packing): Leucine, Isoleucine, Methionine, Phenylalanine
        self.core_deep = ['L', 'L', 'I', 'I', 'I', 'M', 'F']

        # 2. SURFACE RESIDUES (Positions b, c, f)
        # High solubility, helix propensity.
        # Added Y/W here for concentration determination (A280).
        self.surface = ['E', 'K', 'Q', 'E', 'Q', 'A', 'S']
        self.surface_chromophores = ['Y', 'W'] # Specifically for A280

        # 3. INTERFACE/SALT BRIDGE (Positions e, g)
        # Opposing charges to stabilize helix dipole or inter-helix interactions
        self.interface = ['E', 'K', 'Q', 'R', 'D']

        # 4. LOOP RESIDUES
        # Must contain positive charge (R/K) for PFA carboxylate.
        # Must contain turns (G/P).
        self.loop_motifs = [
            ['P', 'R', 'G'],  # The Binder_02 favorite
            ['G', 'K', 'G'],
            ['G', 'R', 'P'],
            ['K', 'P', 'G'],
            ['R', 'G', 'S'],
            ['G', 'R', 'G'],
            ['P', 'K', 'G']
        ]

    def get_residue(self, pool):
        """Helper to pick a random residue from a list."""
        return random.choice(pool)

    def generate_heptad(self, position_type="middle", chromophore_prob=0.05):
        """
        Generates a 7-residue heptad (abcdefg).
        position_type: 'end' (funnel) or 'middle' (bulky core)
        """
        # Determine core type
        if position_type == "end":
            core_pool = self.core_funnel
        else:
            core_pool = self.core_deep

        # Heptad Construction
        # a: Core
        a = self.get_residue(core_pool)

        # b: Surface (chance for chromophore)
        if random.random() < chromophore_prob:
            b = self.get_residue(self.surface_chromophores)
        else:
            b = self.get_residue(self.surface)
```

```
        # c: Surface
        c = self.get_residue(self.surface)

        # d: Core
        d = self.get_residue(core_pool)

        # e: Interface
        e = self.get_residue(self.interface)

        # f: Surface (chance for chromophore)
        if random.random() < chromophore_prob:
            f = self.get_residue(self.surface_chromophores)
        else:
            f = self.get_residue(self.surface)

        # g: Interface
        g = self.get_residue(self.interface)

        return f"{a}{b}{c}{d}{e}{f}{g}"

    def generate_helix(self):
        """
        Generates a helix of ~21-22 residues.
        Structure: [Funnel Heptad] - [Deep Heptad] - [Funnel Heptad]
        This ensures both ends of the helix allow PFA entry.
        """
        h_part1 = self.generate_heptad(position_type="end")
        h_part2 = self.generate_heptad(position_type="middle")
        h_part3 = self.generate_heptad(position_type="end")

        return h_part1 + h_part2 + h_part3

    def generate_loop(self):
        """Selects a loop motif."""
        motif = random.choice(self.loop_motifs)
        return "".join(motif)

    def generate_full_protein(self, index):
        """Assembles the 5-helix bundle."""

        # Start with M
        sequence = "M"

        # Generate 5 helices with 4 connecting loops
        h1 = self.generate_helix()
        l1 = self.generate_loop()

        h2 = self.generate_helix()
        l2 = self.generate_loop()

        h3 = self.generate_helix()
        l3 = self.generate_loop()

        h4 = self.generate_helix()
        l4 = self.generate_loop()

        h5 = self.generate_helix()

        # Assemble
```

```
        # Note: Binder_02 sequence length was approx 124.
        # 3 heptads * 7 * 5 helices = 105. + 4 loops * 3 = 117. + Start/End.
        # This is very close to the target length.
        full_seq = f"{sequence}{h1}{l1}{h2}{l2}{h3}{l3}{h4}{l4}{h5}"

        # Add a C-terminal cap (usually polar/charged)
        full_seq += random.choice(['E', 'Q', 'K']) # Binder_02 had Y/W, but we embedded
those inside

        header = f">PFA_Design_{index:04d} | 5-Helix Bundle | Val-Funnel/Arg-Loop"
        return header, full_seq

# --- Execution ---

def main():
    generator = HelixBundleGenerator()
    unique_sequences = set()
    output_lines = []

    print(f"Generating {NUM_CANDIDATES} candidates...")

    count = 1
    while count <= NUM_CANDIDATES:
        header, seq = generator.generate_full_protein(count)

        if seq not in unique_sequences:
            unique_sequences.add(seq)
            output_lines.append(header)
            # Wrap sequence to 80 chars for standard FASTA formatting
            output_lines.append(textwrap.fill(seq, width=80))
            count += 1

    # Write to file
    with open(FILENAME, "w") as f:
        f.write("\n".join(output_lines))

    print(f"Success! {len(unique_sequences)} unique sequences written to {FILENAME}")
    print("Example sequence:")
    print(output_lines[0])
    print(output_lines[1])

if __name__ == "__main__":
    main()
```

**How this script works:**

1. **Template Preservation:** It adheres to the **Valine-Funnel** concept. The generate_helix function builds 3 segments (heptads) per helix:
    - **Start:** "End/Funnel" heptad (uses V, A at core positions).
    - **Middle:** "Deep" heptad (uses L, I, F, M at core positions).
    - **End:** "End/Funnel" heptad (uses V, A at core positions).
    - This ensures that no matter which end of the bundle the PFA approaches, it finds a sterically accessible pocket.

2. **Loop Constraints:** It randomly selects from a predefined list of loops (PRG, GKG, GRP, etc.) that strictly obey your requirement for positive charge (Arg/Lys) and hairpin formers (Pro/Gly).
3. **Concentration Determination:** It randomly sprinkles **Tyrosine (Y)** and **Tryptophan (W)** into the solvent-exposed surface positions (b, f of the heptad) so you can measure concentration via A280.
4. **Combinatorial Diversity:**
    - It varies the packing residues (choosing between Leu/Ile/Met in the core).
    - It varies the salt-bridging residues (Glu/Lys/Arg/Gln/Asp).
    - It generates over 1,000 unique sequences in standard FASTA format.

You can run this script, and it will produce a file called pfa_binder_library.fasta which you can then feed into your batch prediction pipeline for Boltz or Chai-1.

---

**Prompt 3.**

Great. The code works well. Could you modify the code to make each helix's length as 28 residues?

---

**Response 3.**

I can certainly make that adjustment. To increase the helix length to 28 residues (which corresponds to exactly 4 heptads), I will add an extra "Deep Core" heptad in the middle of the helix generation function. This maintains the "funnel" architecture at both ends while extending the hydrophobic core.

Here is the updated script:

**Summary of Changes**

- **Structure Update:** The generate_helix function now concatenates four heptads instead of three: [Funnel] - [Deep] - [Deep] - [Funnel].
- **Length:** This results in 28 residues per helix (4 × 7).
- **Core Integrity:** By doubling the "middle" section, we create a longer, more stable hydrophobic core for packing, while keeping the accessibility for PFA at the tips.

---

**Prompt 4.**

I would like the loops between helices to have variable lengths of 3–6 residues; they do not need to be the same length within a single chain.

---

**Response 4.**

I've updated the script to include loop motifs ranging from 3 to 6 residues. Since the loop is selected randomly for each connection, this will automatically produce variable loop lengths within the same chain.

---

**Prompt 5.**

I would like you to read the paper I attached and apply the principles of α-helical barrels with a central, accessible channel to the Python code for sequence generation. The authors expanded the traditional heptad repeat.

**Response 5.**

Based on the paper you shared, the key structural innovation for creating central, accessible channels in α-helical barrels (αHBs) is expanding the traditional hpphppp (hydrophobic-polar-polar...) heptad repeat to an **hpphhph** pattern.

In a standard coiled coil, only the a and d positions are hydrophobic, creating a tightly packed core. By shifting to the hpphhph pattern, the e and g positions are also made hydrophobic. This broadens the hydrophobic face of each helix, forcing them to assemble into a wider barrel with a central lumen rather than a solid core—perfect for accommodating your bulky PFA targets.

To implement this, I'll modify the script to assign hydrophobic pools to the e and g interface positions. I'll also boost the charge density of the surface pool (b, c, f) to ensure the overall protein remains highly water-soluble despite the extra hydrophobicity.

Here is the diff to apply this αHB principle to your generator:

This ensures your 6-helix bundle opens up into a genuine barrel structure (as outlined by the Woolfson group), providing a dedicated central pocket for the perfluoroalkyl chain while the flexible, charged loops cap the ends to bind the carboxylate headgroup. Let me know if you'd like to further tune the channel diameter using different hydrophobic residues!

```
import random
import textwrap

# --- Configuration ---
NUM_CANDIDATES = 1200  # Generate slightly more than 1000 to be safe
FILENAME = "pfa_binder_library.fasta"

# --- Design Principles based on PFA_Binder_02 ---
# Topology: 6-helix bundle
# Loop Strategy: Medium length (6 residues), positively charged (Arg/Lys), flexible
(Gly/Pro)
# Core Strategy: "Funnel" shape. Ends of helices are Val/Ala (small), middle is Leu/Ile
(bulky).

class HelixBundleGenerator:
    def __init__(self):
        # Heptad positions: a, b, c, d, e, f, g
        # Hydrophobic Core: a, d
        # Salt-bridges/Interface: e, g
        # Solvent Exposed: b, c, f

        # 1. CORE RESIDUES (Positions a, d)
        # Funnel ends (allow PFA entry): Valine, Alanine
        self.core_funnel = ['V', 'V', 'V', 'A', 'I'] # Weighted towards Val based on
Binder_02
```

```
        # Deep Core (packing): Leucine, Isoleucine, Methionine, Phenylalanine
        self.core_deep = ['L', 'L', 'I', 'I', 'I', 'M', 'F']

        # 2. SURFACE RESIDUES (Positions b, c, f)
        # High solubility, helix propensity.
        # Added Y/W here for concentration determination (A280).
        # Boosted polar/charged frequency to offset the added hydrophobics in e and g.
        self.surface = ['E', 'K', 'Q', 'R', 'D', 'S']
        self.surface_chromophores = ['Y', 'W'] # Specifically for A280

        # 3. INTERFACE RESIDUES (Positions e, g)
        # APPLIED aHB PRINCIPLE: Expanded heptad repeat to 'hpphhph'.
        # Positions e and g are now HYDROPHOBIC to form the wide barrel walls
        # instead of inter-helix salt bridges.
        self.interface_funnel = ['V', 'A', 'S'] # Smaller/polar mix to keep funnel open
        self.interface_deep = ['L', 'I', 'V', 'A'] # Bulky to pack the barrel

        # 4. LOOP RESIDUES (Variable lengths 3-6)
        # Must contain positive charge (R/K) for PFA carboxylate.
        # Must contain flexible/turn residues (G/P/S).
        self.loop_motifs = [
            # Length 3 (Tight turns)
            ['P', 'R', 'G'], ['G', 'K', 'G'], ['R', 'P', 'G'], ['K', 'P', 'G'],
            # Length 4
            ['G', 'P', 'R', 'G'], ['G', 'K', 'P', 'G'], ['P', 'R', 'G', 'S'], ['G', 'R',
'P', 'G'],
            # Length 5
            ['G', 'S', 'R', 'P', 'G'], ['G', 'K', 'G', 'P', 'G'], ['P', 'R', 'G', 'S',
'G'],
            # Length 6 (Flexible wrappers)
            ['G', 'P', 'R', 'G', 'K', 'G'], ['G', 'G', 'K', 'G', 'G', 'P'],
            ['G', 'S', 'R', 'P', 'G', 'G'], ['G', 'K', 'P', 'G', 'R', 'G'],
            ['P', 'G', 'R', 'G', 'S', 'G'], ['G', 'G', 'R', 'R', 'G', 'G'],
            ['P', 'K', 'G', 'G', 'K', 'P']
        ]

    def get_residue(self, pool):
        """Helper to pick a random residue from a list."""
        return random.choice(pool)

    def generate_heptad(self, position_type="middle", chromophore_prob=0.05):
        """
        Generates a 7-residue heptad utilizing the hpphhph aHB pattern.
        position_type: 'end' (funnel) or 'middle' (bulky core)
        """
        # Determine core and interface types
        if position_type == "end":
            core_pool = self.core_funnel
            interface_pool = self.interface_funnel
        else:
            core_pool = self.core_deep
            interface_pool = self.interface_deep

        # Heptad Construction (hpphhph)
        # a: Core (h)
        a = self.get_residue(core_pool)

        # b: Surface (p)
        if random.random() < chromophore_prob:
            b = self.get_residue(self.surface_chromophores)
```

```
        else:
            b = self.get_residue(self.surface)

        # c: Surface (p)
        c = self.get_residue(self.surface)

        # d: Core (h)
        d = self.get_residue(core_pool)

        # e: Interface (h) - Changed to hydrophobic for aHB barrel formation
        e = self.get_residue(interface_pool)

        # f: Surface (p)
        if random.random() < chromophore_prob:
            f = self.get_residue(self.surface_chromophores)
        else:
            f = self.get_residue(self.surface)

        # g: Interface (h) - Changed to hydrophobic for aHB barrel formation
        g = self.get_residue(interface_pool)

        return f"{a}{b}{c}{d}{e}{f}{g}"

    def generate_helix(self):
        """
        Generates a helix of 28 residues (4 heptads).
        Structure: [Funnel] - [Deep] - [Deep] - [Funnel]
        This ensures both ends allow PFA entry while increasing core stability.
        """
        h_part1 = self.generate_heptad(position_type="end")
        h_part2 = self.generate_heptad(position_type="middle")
        h_part3 = self.generate_heptad(position_type="middle")
        h_part4 = self.generate_heptad(position_type="end")

        return h_part1 + h_part2 + h_part3 + h_part4

    def generate_loop(self):
        """Selects a loop motif."""
        motif = random.choice(self.loop_motifs)
        return "".join(motif)

    def generate_full_protein(self, index):
        """Assembles the 6-helix bundle."""

        # Start with M
        sequence = "M"

        # Generate 6 helices with 5 connecting loops
        h1 = self.generate_helix()
        l1 = self.generate_loop()

        h2 = self.generate_helix()
        l2 = self.generate_loop()

        h3 = self.generate_helix()
        l3 = self.generate_loop()

        h4 = self.generate_helix()
        l4 = self.generate_loop()
```

```
        h5 = self.generate_helix()
        l5 = self.generate_loop()

        h6 = self.generate_helix()

        # Assemble
        # Note: 4 heptads * 7 * 6 helices = 168. + 5 loops * ~4.5 avg = ~22. + Start/End.
        # Length will be approx 192 residues.
        full_seq = f"{sequence}{h1}{l1}{h2}{l2}{h3}{l3}{h4}{l4}{h5}{l5}{h6}"

        # Add a C-terminal cap (usually polar/charged)
        full_seq += random.choice(['E', 'Q', 'K']) # Binder_02 had Y/W, but we embedded
those inside

        header = f">PFA_Design_{index:04d} | 6-Helix Barrel | Val-Funnel | hpphhph
pattern"
        return header, full_seq

# --- Execution ---

def main():
    generator = HelixBundleGenerator()
    unique_sequences = set()
    output_lines = []

    print(f"Generating {NUM_CANDIDATES} candidates...")

    count = 1
    while count <= NUM_CANDIDATES:
        header, seq = generator.generate_full_protein(count)

        if seq not in unique_sequences:
            unique_sequences.add(seq)
            output_lines.append(header)
            # Wrap sequence to 80 chars for standard FASTA formatting
            output_lines.append(textwrap.fill(seq, width=80))
            count += 1

    # Write to file
    with open(FILENAME, "w") as f:
        f.write("\n".join(output_lines))

    print(f"Success! {len(unique_sequences)} unique sequences written to {FILENAME}")
    print("Example sequence:")
    print(output_lines[0])
    print(output_lines[1])

if __name__ == "__main__":
    main()
```

#### Supplemental Note 7. LLM evaluation of LLM-generated metal binding designs

*Prompt:*
I previously had an LLM design de novo metal binding proteins. They are designed form a dimer of helix-turn-helix hairpins, creating a metal binding four helix bundle. After predicting the structures of the designs and filtering, these are the "top" candidates that I chose. In each design, which residues (identity and residue #) do you anticipate will be part of a metal binding site? Of these candidates, which ones do you think should bind Zn(II) vs Cu(II)? Do you think they will form parallel or antiparallel dimers? Of the sequences given, which do you anticipate will be the best metal binding protein when I do experiments to validate?

LMBP-01 MAELIAKLEQELKHVADALEVHQKSGDPEKLARIQQELDALNVVHEALKELHKQALEQAL

LMBP-02 VNSITNELEKIHTSLENIESNLSSIEEQGGSGLQQIEKNATEIHKELSEISSSLQKIESK

LMBP-03 LTKIENNLEKFHQQLEEASEQLTKISTSGGSGYQQIQKTLKTVHESVNKVQENANSIESQ

LMBP-04 VQSYSKELTKLHTTLQNIENQFQKVNKTGGSGLNTIQSNLTSLHQELEEFTNQLNKLNQE

LMBP-05 AKKFSETYQKIHNSLSEITNEIETYKTNGGSGLKEYTTSLNSIHTELSTISKTVNEFTTT

LMBP-06 LTKVETTLEQLHKKIETIEQKFKSLSNQGGSGYKNIQSTVNKISNHINEIESQLETASET

LMBP-07 AEKIEEKHAEKIQEKAEKIQEKHAEEKIGGPGGPGGKEAEKIHKEQAEKIKEAAEKIKEKHAEEKI

LMBP-08 AEKIEEKHAEKIQEKAEKIQEKHAEEKIGGPGAPGAGKEAEKIHKEQAEKIKEAAEKIKEKHAEEKI

LMBP-09 IEKAEEKHIEKAQEKIEKAQEKHIEEKAGGPGGPGGKEIEKIHKEQIEKAQEAIEKAQEKHIEEKA

LMBP-10 MKQLEKHLLAQLKHALDQLEKALGGSGSGSLQALHKELLAQLDHALDQLEKAL

LMBP-11 MEQLDKELLAQLHKALDELEKALGGSGTGSLQAHKKELLAQLDKALDQLEKAL

LMBP-12 MQQLDKELLAQLHKALDQLEKALGGSGTGSLQALKHELLAQLDKALDQLEKAL

*Claude Sonnet 4.6 response:*

I'll analyze these sequences systematically. Let me work through the structure, metal binding residues, metal selectivity, oligomerization geometry, and rank them.

##### Structural Analysis

All 12 designs follow a **helix-turn-helix hairpin → dimer → four-helix bundle** architecture. The linker region (GGS/GPGG/TGSG motifs) separates the two helices within each monomer, and the dimer interface buries the hydrophobic core while positioning metal-coordinating residues at the bundle interior.

---

##### Metal Binding Residue Predictions

The key coordinating residues in designed metal-binding bundles are **His (H), Cys (C), Glu (E), Asp (D), and occasionally Met (M)**. I'll identify likely coordinating residues at the helix-helix interface, noting that in a four-helix bundle the binding site is typically formed by residues from all four helices at conserved heptad positions (a/d positions facing inward).

---

**LMBP-01**

MAELIAKLEQELKHVADALEVHQK | SGDP | EKLARIQQELDALNVVHEALKELHKQALEQAL

- **Helix 1:** H14, H21 (VHQK region), E10
- **Helix 2 (after linker):** H37 (VHE), H40 (ELKELHK → H40, H43)
- **Predicted coordinating residues: H14, H21, H40, H43**
- Classic His-rich four-helix bundle — **strong Zn(II) candidate**

---

**LMBP-02**

VNSITNELEKIHTSLENIESNLSS | IEEQGGS | GLQQIEKNATEIHKELSEISSSLQKIESK

- **Helix 1:** H11 (KIH), E8, E19
- **Helix 2:** H24 (TEIH → H24 counting from linker), E26 (KELS)
- **Predicted coordinating residues: H11, E8, H24, E26**
- Mixed His/Glu coordination — **Zn(II) or Cu(II) candidate**

---

**LMBP-03**

LTKIENNLEKFHQQLEEASEQLTKIST | SGGSGY | QQIQKTLKTVHESVNKVQENANSIESQ

- **Helix 1:** H11 (KFH), E15, E16 (QLEE)
- **Helix 2:** H22 (TVHE counting from linker)
- **Predicted coordinating residues: H11, E15, E16, H22**
- Note the **FH motif** (Phe-His) is common in designed bundles for positioning His into the core
- **Zn(II) candidate**

---

**LMBP-04**

VQSYSKELTKLHTTLQNIENQFQKVNKT | GGSGL | NTIQSNLTSLHQELEEFTNQLNKLNQE

- **Helix 1:** H11 (KLH)
- **Helix 2:** H21 (SLH counting from linker), E23 (QELE), E24
- **Predicted coordinating residues: H11, H21, E23, E24**
- Relatively His-poor, Glu-rich — **moderate Zn, possible Cu**

---

**LMBP-05**

AKKFSETYQKIHNSLSEITNEIETYKT | NGGSGL | KEYTTSLNSIHTELSTISKTVNEFTTT

- **Helix 1:** H11 (KIH), E14 (NEI... actually E21 EIETY region)
- **Helix 2:** H20 (SIH counting from linker), E22 (TELS)
- **Predicted coordinating residues: H11, H20, E14, E22**
- **Zn(II) candidate**

---

**LMBP-06**

LTKVETTLEQLHKKIETIEQKFKSLSN | QGGSGY | KNIQSTVNKISNHINEIESQLETASET

- **Helix 1:** H11 (QLH), E8 (TLEQLH adjacent Glu)
- **Helix 2:** H21 (SNH counting from linker), E24 (EIES)
- **Predicted coordinating residues: H11, E8, H21, E24**
- **Zn(II) candidate**

---

**LMBP-07**

AEKIEEKHAEKIQEKAEKIQEKHAEEKI | GGPGGPGG | KEAEKIHKEQAEKIKEAAEKIKEKHAEEKI

- This is a **repeat helix** — very regular **ExxKxxH** heptad pattern
- **Helix 1:** H8 (EKH), H22 (EKH repeat), E2, E6, E13, E20
- **Helix 2:** H7 (KIH counting from linker), H24 (KH), multiple E residues
- **Predicted coordinating residues: H8, H22, H7(linker+), H24** with glutamate secondary coordination from E positions
- The GPGGPGG linker is highly flexible — dimer geometry may be variable
- **Strong Cu(II) candidate** — multiple His in a row, flexible geometry ideal for Cu

---

**LMBP-08**

AEKIEEKHAEKIQEKAEKIQEKHAEEKI | GGPGAPGAG | KEAEKIHKEQAEKIKEAAEKIKEKHAEEKI

- Nearly identical to LMBP-07 with a slightly more rigid GPGAPGAG linker
- **Same coordinating residues: H8, H22, H7(linker+), H24**
- Similar repeat EKH heptad pattern
- **Cu(II) candidate** — same reasoning as LMBP-07; slightly more defined geometry due to Ala in linker

---

**LMBP-09**

IEKAEEKHIEKAQEKIEKAQEKHIEEKAGG | PGGPGG | KEIEKIHKEQIEKAQEAIEKAQEKHIEEKA

- Repeat **ExxKxxH** pattern, similar to 07/08 but with **IEK** vs **AEK** — more hydrophobic character
- **Helix 1:** H7 (EKH), H21 (EKH), multiple E flanking
- **Helix 2:** H7 (KIH from linker), H22 (KH)
- **Predicted coordinating residues: H7, H21, H7′, H22′** (across dimer)
- **Strong Cu(II) candidate** — highest His density of the series, flexible PGGPGG linker

---

**LMBP-10**

MKQLEKHLLAQLKHALDQLEKAL | GGSGSGSL | QALHKELLAQLDHALDQLEKAL

- **Helix 1:** H6 (EKH), H12 (QLK**H**), D15 (HALD → D16), E18 (QLE)
- **Helix 2:** H5 (KH counting from linker), D12 (QLDH), D15, E18
- **Predicted coordinating residues: H6, H12, D15/D16, H5′, D12′**
- **His + Asp** coordination — unusual and interesting
- **Zn(II) candidate** with possible mixed His/Asp site

---

**LMBP-11**

MEQLDKELLAQLHKALDELEKALGGSGTGSL | QAHKKELLAQLDKALDQLEKAL

- **Helix 1:** H12 (QLH), D14 (KALD), E15, E17 (ELEK)
- **Helix 2:** H4 (QAH counting from linker), D13 (QLDKALD → D13/D16), E18
- **Predicted coordinating residues: H12, D14, H4′, D13′**
- His/Asp/Glu mixed coordination
- **Zn(II) candidate**

---

**LMBP-12**

MQQLDKELLAQLHKALDQLEKALGGSGTGSL | QALKHELLAQLDKALDQLEKAL

- Nearly identical to LMBP-11 with minor substitutions (Q→Q, E→E region)
- **Helix 1:** H12 (QLH), D14 (KALD), E16
- **Helix 2:** H5 (KH from linker), D13, D16
- **Predicted coordinating residues: H12, D14, H5′, D13′**
- **Zn(II) candidate** — very similar to LMBP-11

---

**Metal Selectivity Summary**

| Design | Predicted Metal | Reasoning |
|---|---|---|
| LMBP-01 | **Zn(II)** | 4×His tetrahedral — classic Zn coordination |
| LMBP-02 | **Zn(II)** | 2His/2Glu — common Zn motif |
| LMBP-03 | **Zn(II)** | 2His/2Glu, FH positioning motif |
| LMBP-04 | **Zn(II)/Cu(II)** | His-poor, Glu-rich — borderline |
| LMBP-05 | **Zn(II)** | 2His/2Glu balanced |
| LMBP-06 | **Zn(II)** | 2His/2Glu balanced |
| LMBP-07 | **Cu(II)** | High His density, flexible linker, repeat motif |
| LMBP-08 | **Cu(II)** | Same as 07, slightly more rigid |
| LMBP-09 | **Cu(II)** | Highest His density, flexible linker |
| LMBP-10 | **Zn(II)** | His/Asp — tetrahedral preference |

| Design | Predicted Metal | Reasoning |
|---|---|---|
| LMBP-11 | **Zn(II)** | His/Asp/Glu mixed |
| LMBP-12 | **Zn(II)** | His/Asp/Glu mixed, nearly identical to 11 |

**Key reasoning:** Cu(II) favors planar or distorted geometries with multiple His nitrogen donors and tolerates flexible coordination spheres. Zn(II) strongly prefers tetrahedral geometry with His/Cys/Asp/Glu — rigid, pre-organized sites. The repeat-motif designs (07–09) have too many potential His donors and too flexible a linker for selective Zn binding, making them better Cu candidates by the **entatic state** and **Irving-Williams series** logic.

---

**Parallel vs. Antiparallel Dimer Geometry**

This is determined by the **hydrophobic packing at the dimer interface** and **charge complementarity** of the helices:

| Design | Predicted Geometry | Reasoning |
|---|---|---|
| LMBP-01 | **Antiparallel** | Alternating charged/hydrophobic pattern, no clear N→C gradient |
| LMBP-02 | **Antiparallel** | Charge distribution symmetric, ExxK pattern |
| LMBP-03 | **Antiparallel** | Similar symmetric charge pattern |
| LMBP-04 | **Antiparallel** | Symmetric heptad repeat logic |
| LMBP-05 | **Antiparallel** | Balanced charge distribution |
| LMBP-06 | **Antiparallel** | Symmetric |
| LMBP-07 | **Antiparallel** | ExxKxxH repeat is symmetric by design — antiparallel maximizes complementarity |
| LMBP-08 | **Antiparallel** | Same as 07 |
| LMBP-09 | **Antiparallel** | Same repeat logic |
| LMBP-10 | **Parallel** | LKHL…QLEKAL motif has directional hydrophobic gradient; SGSGSL linker constrains orientation |
| LMBP-11 | **Parallel** | Very similar to LMBP-10, short TGSL linker constrains parallel registry |
| LMBP-12 | **Parallel** | Same as LMBP-11 |

**Reasoning for 10–12:** The shorter, more rigid glycine-serine-threonine linkers in LMBP-10/11/12 (GGSGSGSL, GGSGTGSL) are less flexible than the GPGG-type linkers, which tends to favor **parallel** topology because the linker length constrains how the second helix can exit relative to the first. In contrast, the longer GPGGPGG linkers in 07–09 allow the conformational freedom needed for antiparallel packing driven by charge-charge complementarity.

---

**Overall Ranking & Best Experimental Candidate**

Here is my ranked order with justification:

| Rank | Design | Rationale |
|---|---|---|
| 🥇 1 | **LMBP-01** | 4×His coordination, clean sequence, well-spaced His (H14, H21, H40, H43), no ambiguity in binding site, high predicted structural regularity, strong Zn(II) selectivity |
| 🥈 2 | **LMBP-10** | Compact, His/Asp coordination mimics natural zinc fingers, short rigid linker for parallel dimer gives well-defined geometry, less ambiguous than 11/12 |
| 🥉 3 | **LMBP-09** | If Cu(II) binding is the goal, highest His density, most favorable for Cu coordination chemistry |
| 4 | **LMBP-11** | Very similar to 10/12 but slightly more varied sequence (E residues adds coordination flexibility) |
| 5 | **LMBP-12** | Near-duplicate of 11, redundant |
| 6 | **LMBP-06** | Balanced Zn candidate |
| 7 | **LMBP-03** | FH positioning motif is a good design feature |
| 8 | **LMBP-08** | Cu candidate with better-defined linker than 07 |
| 9 | **LMBP-07** | Cu candidate, overly flexible linker may reduce affinity |
| 10 | **LMBP-05** | Functional but less distinctive |
| 11 | **LMBP-02** | Mixed coordination, lower confidence |
| 12 | **LMBP-04** | His-poor, Glu-heavy — least well-defined site |

**Why LMBP-01 is the top pick:**

1. **Four histidines** (H14, H21, H40, H43) symmetrically positioned — a textbook tetrahedral Zn(II) site
2. Reasonable hydrophobic core residues (Leu, Ala, Val, Ile) for structural stability
3. The linker (SGDP) is short and contains a Pro — this will **kink** the turn and enforce a defined hairpin geometry, reducing conformational heterogeneity
4. Clean charge distribution for solubility
5. No competing His residues outside the predicted binding site

---

**Caveats**

- Without sequence→structure prediction results (AlphaFold2/RFdiffusion outputs), the exact heptad register and buried vs. exposed positions are inferred
- Actual metal affinity depends critically on pre-organization of the binding site, which requires experimental or computational validation
- Cys-containing designs would be far stronger metal binders — the absence of Cys in all 12 is notable and will limit affinity compared to natural zinc fingers

- The repeat designs (07–09) may aggregate due to their repetitive hydrophobic patterns

*Claude Opus response:*

Reading these from sequence alone (I don't have your predicted structures), so my layer assignments come from heptad register and loop position. If you share the PDBs I can be much more specific about which His are actually within coordination distance.

A few things jumped out immediately that matter more than the site analysis:

- **There isn't a single Cys in any of these.** Every site is all-His/carboxylate.
- **Only four sequences have aromatics** (03, 04, 05, 06). LMBP-01 and 07–12 have no Trp/Tyr/Phe, so A280 won't give you a reliable concentration — you'll need quantitative amino acid analysis or a colorimetric assay, and concentration errors will wreck any Kd you measure.
- If you express these with a His6 tag, **cleave it before any binding experiment.** A polyhistidine tag will out-compete a bis-His site outright.

**Predicted ligand sets**

I've grouped them, since 02–06 and 10–12 are clearly variations on two templates.

| Design | His residues | Nearby carboxylates on the same face | Likely donor set per site |
|---|---|---|---|
| LMBP-01 | H14, H22, H45, H52 | D17, D39, E46, E50 | His2 (+ Glu/Asp), spread over 4 layers |
| LMBP-02 | H12, H44 | E16, E42, E46 | His2 + 2 solvent |
| LMBP-03 | H12, H44 | E16, E17, E45 | His2 + 2 solvent |
| LMBP-04 | H12, H44 | E46, E48, E49 | His2 + Glu |
| LMBP-05 | H12, H44 | E17, E46 | His2 + 2 solvent |
| LMBP-06 | H12, **H46** | E16, E49 | His2, offset (see below) |
| LMBP-07 | H8, H23, H43, H61 | E-rich throughout | **His4 ×2** |
| LMBP-08 | H8, H23, **H44, H62** | E-rich throughout | **His4 ×2** |
| LMBP-09 | H8, H23, H43, H61 | E-rich throughout | **His4 ×2** |
| LMBP-10 | H7, H14, H35, H44 | D17, D43, D47 | His2 + Asp ×2 |
| LMBP-11 | H13, H34 | D43, D47, E18 | His + Asp, weak |
| LMBP-12 | H13, H36 | D43, D47 | His + Asp, weak |

**The structural point that governs everything:** a C2-symmetric homodimer can only give you an *even* number of His per site (2 or 4), unless a residue sits directly on the symmetry axis. You cannot get a His3 site out of this architecture. That's a real constraint, because His3 + solvent is the canonical geometry for a designed Zn(II) site — the C2 symmetry pushes you toward His4, which is Cu(II) territory.

**Parallel vs. antiparallel**

Every hairpin is intramolecularly antiparallel (loops are 4–9 residues, far too short to cross the bundle). The question is how the two hairpins pack.

**LMBP-02 through 05 only make sense as antiparallel dimers** (2-fold perpendicular to the bundle axis, loops at opposite ends). Working in layers from the loop: H12 sits ~12 residues up helix 1, H44 sits ~17 residues up helix 2. In the same monomer they're ~7.5 Å apart vertically — too far to share a metal. But flip the second hairpin and H12 of chain A lands in the same layer as H44 of chain B, on an adjacent helix. That gives you two independent bis-His sites at opposite ends of the bundle. I'd read this as the intended design.

**LMBP-06 breaks that.** H46 instead of H44 shifts the second His by two residues (~3 Å and ~200° of helical rotation), which destroys the antiparallel pairing and only partially rescues it in a parallel arrangement. If 02–05 were the template, 06 looks like a design slip rather than a variant worth testing.

**LMBP-07/08/09 are the robust ones topologically.** Within a single hairpin, H23 and H43 land in essentially the same layer near the loop (heights ~23 and ~22), and H8/H61 land near the other end. So you get a 4-His cluster at each end in a parallel dimer, *and* a 4-His cluster at each end in an antiparallel dimer (A's H23/H43 meeting B's H8/H61). The site survives either topology. The flip side: the GGPGGPGG loop is 8–9 residues of Gly/Pro — long and floppy enough that I'd expect a mixture of topologies in solution, which will broaden your NMR and complicate any structure.

**LMBP-10/11/12 are the least specific.** Helix 1 and helix 2 are nearly identical in sequence (residues 4–23 vs. 34–53 differ at only 3–4 positions), and the loops are 7-residue Gly/Ser. There's very little energetic difference between the two dimer topologies, and no barrier to domain-swapped or higher-order species. Expect a heterogeneous ensemble.

**Zn(II) vs. Cu(II)**

Blunt version first: **Irving–Williams means all of these will bind Cu(II) more tightly than Zn(II)**, probably by 3–5 orders of magnitude. Nitrogen-only sites with no thiolate and no rigid tetrahedral enforcement don't give Zn selectivity. Any "Zn design" here is really a Cu design that also binds Zn.

With that said:

**Cu(II) — LMBP-07, 08, 09.** Four imidazoles from four helices around a 2-fold is close to tetragonal N4, which is exactly what Cu(II) wants (Jahn–Teller stabilization in the equatorial plane, weak axial solvent). These should show the classic ~600 nm d-d band and a 4N EPR signature.

**Zn(II) — LMBP-01, 10, 12.** These have His plus buried Asp/Glu (D17/D39 in 01; D43/D47 in 10 and 12) and a lower His count, which is more compatible with a distorted tetrahedral His2(carboxylate)(solvent) site. LMBP-10 is the best of these — 4 His, and D43/D47 sit on the same face.

**LMBP-02 through 05** are bis-His + 2 solvent. That's a genuine Cu(II) site (µM-ish is realistic) and a weak Zn site (I'd guess high µM to mM).

**LMBP-11** I'd deprioritize entirely: only 2 His, and H34 appears to sit at a core a/d position, meaning it's buried against the hydrophobic core rather than pointing into a solvent-accessible cavity.

**What I'd bet on**

**LMBP-09** as the primary candidate. It has the most His (8 per dimer), the site tolerates either dimer topology, and the E/K/A/I composition gives high helical propensity and excellent solubility. Critically, it swaps the Ala/Ile core assignment relative to 07 — LMBP-07's Ala-heavy core layer is a molten-globule risk, and 09's β-branched Ile at those positions should pack better and give you a cooperative thermal melt.

**LMBP-10** as the second, and as your Zn candidate. The Leu/Ala core is the best-designed of the set and should fold into a genuinely stable bundle. Its weakness is topological ambiguity, not folding.

**LMBP-02 through 05** I'd expect to be marginally folded. They're loaded with Ser/Thr/Asn (LMBP-05 in particular: T, S, N, Y everywhere), which are poor helix formers and give you very little buried hydrophobic surface. AlphaFold will confidently predict helices for these regardless — low helical propensity is exactly the failure mode structure prediction doesn't catch. I'd run CD on one of them before investing in the whole set.

One design tension worth keeping in mind: a very well-packed core competes with metal binding, because the site has to be pre-organized and slightly under-packed to accommodate the ion. The DeGrado maquette lineage exploited this deliberately — molten-globule scaffolds that fold *upon* metal binding. LMBP-09's tighter core may give you a more stable protein and a worse metal site than LMBP-07.

**Experiments that will actually discriminate**

Before any titration: SEC-MALS or AUC to confirm you have a dimer and not a tetramer or an ensemble. Harbury's rules say Leu at *a* / Ile at *d* favors a four-helix arrangement — your Ala/Ile and Leu/Ala cores are off-book, so don't assume the oligomeric state.

For the sites themselves, **EPR on the Cu(II) complexes is the highest-information experiment you can run.** The $g_\parallel$ and $A_\parallel$ values on a Peisach–Blumberg plot will tell you directly whether you have 4N, 3N1O, or 2N2O coordination — that distinguishes a real His4 site in 07/08/09 from an adventitious surface site, which no binding isotherm will do.

For Zn, use Co(II) substitution first. The d-d band intensity in the 500–600 nm region reports coordination number ($\varepsilon \approx 300–600\ M^{-1}cm^{-1}$ for 4-coordinate, much lower for 6), so you'll know whether you built a tetrahedral site before you spend time on affinity. Then measure Kd by competition against a chelator of known affinity (mag-fura-2 or Zincon) rather than by direct ITC — direct titration will just report the buffer.

Do everything at pH 7.0–7.5. His pKa is around 6, so you'll lose the site below that, and $Zn(OH)_2$ starts limiting you above pH 8.
